# Reverse engineering environmental metatranscriptomes clarifies best practices for eukaryotic assembly

**DOI:** 10.1101/2022.04.25.489326

**Authors:** Arianna I. Krinos, Natalie R. Cohen, Michael J. Follows, Harriet Alexander

## Abstract

Diverse communities of microbial eukaryotes in the global ocean provide a variety of essential ecosystem services, from primary production and carbon flow through trophic transfer to cooperation via symbioses. Increasingly, these communities are being understood through the lens of omics tools, which enable high-throughput processing of communities of multiple species. Metatranscriptomics in particular offers an understanding of real-time gene expression in microbial eukaryotic communities, providing a window into community metabolic activity. However, these approaches are rarely validated. A systematic validation of metatranscriptome assembly and annotation methods is necessary to appropriately assess (1) the fidelity of our community composition measurements and (2) the confidence of novel taxonomic and functional content discovered with metatranscriptomics. Here, we present *euk* `rhythmic`, an automated and customizable multi-assembler workflow for eukaryotic metatranscriptome assembly, annotation, and analysis, and validate the ability of the pipeline to recapitulate simulated eukaryotic community-level expression data treated as a gold standard. Finally, we apply this pipeline to two previously published metatranscriptomic datasets, showing both recapitulation of previous results as well as enhanced taxonomic and functional gene discovery.

## Introduction

Eukaryotic microbes play diverse and important roles in global ecosystems (***Massana and Pedrós-Alió, 2008***), including grazing processes, primary production, and acting as hosts for diverse and essential symbionts (***Worden et al., 2015***). In ocean ecosystems in particular, the literature on the role of eukaryotic microbes in ecosystem processes continues to expand (***Caron et al., 2017***). This literature provides further evidence that eukaryotic microbes are as important as prokaryotic counterparts when it comes to nutrient cycling, and have outsized influence on food webs and community ecology (***Caron et al., 2009***; ***Lawler and Morin, 1993***; ***Stoecker, 1998***; ***Sherr and Sherr, 2002***), necessitating renewed efforts of understanding why.

The ecological relevance of eukaryotic microbes requires careful study of their ecology and distribution, but this can be difficult to execute, both *in situ* and in the laboratory. Taxonomic diversity can be catalogued in the field using 18S rRNA gene amplicons or even count data, but this completely neglects functional annotation, which can be leveraged to create biogeochemical understanding. Furthermore, the diversity of natural eukaryotic assemblages makes comprehensive surveys difficult to perform in a taxon-specific manner. Moreover, many eukaryotic microbes are not easily cultivatable in the lab (***Massana and Pedrós-Alió, 2008***; ***Del Campo et al., 2016***). Thus, relying solely on the subset of eukaryotic microbes that we can grow in the laboratory may apply a biased filter to our understanding of these organisms (***Del Campo et al., 2013***). For this reason, sequencing techniques have become a popular and successful method of uncovering new taxonomic and functional diversity in populations of eukaryotic microbes in diverse environments in the field (***Del Campo et al., 2016***).

Metatranscriptomics has become a widespread and promising approach to answer questions about microbial communities in the environment without prior knowledge or bias (***Gifford et al., 2011***), and may be used to identify underlying genetic mechanisms driving global phenomena like ocean biogeochemistry (***Becker et al., 2021***; ***Salazar et al., 2019***; ***Stewart et al., 2012***). Metatranscriptomes provide an accessible means to look at the full suite of genes being expressed by a group of organisms, which may be partitioned by size, site, or phylogenetic origin (***John et al., 2009***). However, despite the potential of this approach, the field is relatively new and immature. The first environmental transcriptome, targeting bacterioplankton, was sequenced in 2005 (***Poretsky et al., 2005***), and environmental metatranscriptomes began to appear in the literature around 2008 (***Gilbert et al., 2008***; ***John et al., 2009***). Metatranscriptomes offer a snapshot of the whole community at the time of sequencing, but the relative proportion of transcripts and their detectability may not always provide meaningful insights into true biological processes, in particular when sequencing depth is low or references are missing from the database (***Gifford et al., 2011***). For this reason, databases must be compiled and new computational approaches must and continue to be developed to process and interpret metatranscriptomic data. The collation of laboratory transcriptomic data to a single location and format by the Marine Microbial Eukaryote Transcriptome Sequencing Project (MMETSP) (***Keeling et al., 2014***; ***Caron et al., 2017***) began as a repository effort and became one of the most important databases enabling the identification of marine microbial eukaryotes from metatranscriptomic sequences (e.g. ***Krinos et al. (2021)***; ***Carradec et al. (2018)***; ***Alexander et al. (2015)***). Substantial discoveries have been made using sequenced metatranscriptomes, including novel explanations for persistent gaps in ecological understanding, such as coexistence within a seemingly narrow niche (***Alexander et al., 2015***), discovering new genes or putative organisms from previously unknown sequences (***Gilbert et al., 2008***), developing a molecular understanding of the basis of coral disease (***Daniels et al., 2015***), and decoding the complexities of deep-sea hydrothermal vent microbial communities (***Lesniewski et al., 2012***). Metatranscriptomic data availability, in particular for eukaryotic phytoplankton, has been transformed by the *Tara* Oceans Project, which provided an unprecedented amount of metagenomic and metatranscriptomic information, enabling us to better interpret the ocean genetic landscape on a global scale (***Richter, 2017***). Still, metatranscriptome analysis tends to vary substantially between studies, and interpretation can suffer from biases inherent to the technology. While metatranscriptomes offer a snapshot of the whole community at the time of sequencing, the relative proportion of transcripts and their detectability may not always provide meaningful insights into true biological processes.

Reliable, reproducible, and broadly available approaches to metatranscriptome analysis have been lacking, particularly in eukaryotic microbial community assessment. Early transcriptome pipelines were designed for conventional, well-studied organisms, such as humans (e.g. ***Leimena et al. (2013)***). These pipelines are unlikely to include user-downloadable software, and often are focused on annotation and do not include a mechanism for *de novo* assembly and processing (***Leimena et al., 2013***). Soon thereafter, the first pipeline for uncharacterized microbial communities emerged, but it was a description of the steps needed for metatranscriptome analysis, rather than as software products available to users (***Davids et al., 2016***). The Simple Annotation of Metatranscriptomes by Sequence Analysis (SAMSA) tool, and its second released version, SAMSA2, are among the most recently updated metatranscriptome analysis tools (***Westreich et al., 2018***). While this tool is a complete package that can be downloaded and used by scientists, it focuses on rRNA removal steps, and does not include assembly steps (***Westreich et al., 2018***). In fields such as microbial oceanography, we often require *de novo* assembly of transcriptome sequences, as the identity of the organisms in environmental samples is not always known, and even for well-understood organisms, comprehensive references may not be available. To date, metatranscriptome pipelines have either lacked accompanying software products or assembly steps necessarily for *de novo* environmental analysis. As a consequence, the scientific community is in need of a reliable metatranscriptome analysis tool that is downloadable, reproducible, and includes *de novo* transcriptome assembly. Further, a protocol for validating popular metatranscriptomic assessments, and a set of recommendations for how best to balance and manage the challenge of minimizing the computational cost of metatranscriptome assembly and post-processing, is necessary to maximize the ecological insight that can be extracted from these powerful data.

The landscape of *de novo* transcriptome assembly tools is wide, and often there is disagreement about which tool is best to use for a particular application or the average expression level for a sequenced transcript (***Vijay et al., 2013***). The Oyster River Protocol (ORP) software was published as a method to optimize single-organism transcriptome assembly via the integration of many assemblers and *k*-mer sizes (***MacManes, 2018***).The ORP, however, is a standalone approach to transcriptome assembly, and does not allow the user to simultaneously process multiple samples, nor was it designed to accommodate metatranscriptomes or accommodate downstream annotation. More recently, it has been demonstrated that *de novo* co-assembly using multiple transcriptome assemblers improves the quality of single-organism transcriptome assembly (***Ortiz et al., 2021***). This was shown using a *de novo* transcriptome assembly pipeline with non-model organism RNAseq data as input to recapitulate the transcriptome of a single species. The authors of this study used a BUSCO score (or the relative presence of lineage-specific single-copy genes (***Simão et al., 2015***; ***Jauhal and Newcomb, 2021***)) quality threshold of 50% recovery in order to assess the recovery of single organism transcriptomes (***Ortiz et al., 2021***). The authors note that multiple assemblers used at once for a larger co-assembly contribute to higher quality transcriptomic assemblies of RNAseq data, especially when some subset of the highest-performing assemblers is used (***Ortiz et al., 2021***). The results from these two co-assembly studies may help inform multi-organism metatranscriptomic community data, but they require a transition from consideration of single-organism BUSCO metrics to identification of key features of multiple organisms present in an environmental community, or the use of a known, simulated community treated as a gold standard.

The question that remains from single-organism co-assembly studies is why individual transcriptomic assemblers sometimes produce higher-quality or more complete results, and whether redundancy within each transcriptomic assembly skews quality assessment. In order to answer this question, the assembled content shared in the output of multiple assemblers needs to be compared to the new content offered by combining assembly tools. Here, we assess the ability of metatranscriptomic assembly methods, and specifically our co-assembler, multiple-sample assembly approach to recover *all* transcripts included from existing single-organism transcriptome assemblies. Rather than testing the recovery of identified database genes, we compare our metatranscriptome assemblies to annotated “designer” metatranscriptomes constructed from diverse transcriptome assemblies from a database created using a collection of transcriptomes from marine microbial eukaryotes, the Marine Microbial Eukaryote Transcriptome Sequencing Project (MMETSP) (***Keeling et al., 2014***; ***Caron et al., 2017***). We used database sequences to evaluate whether collecting and assembling metatranscriptomes from a mixed environmental community like the ocean is comparable to isolating and sequencing particular species or strains of eukaryotic marine microbes. Our validation workflow is designed to answer the questions: Do studies that use metatranscriptomics to understand community diversity in eukaryotic microbes found in the environment (a) adequately recapitulate the taxonomic and functional diversity found in those communities? and (b) reproduce consistent *sequences* which could be reliably recovered with repeated sampling and assembly? Further, we test whether combining assembly methods and samples improves the skill of transcript recovery.

## Results

Throughout this paper, we use: “designer metatranscriptomes” to refer to the “gold standard” `jEUKebox`-simulated metatranscriptomic contigs generated from MMETSP reference transcriptomes with known taxonomic annotations, “simulated raw reads” to refer to simulated raw metatranscriptomic reads, and “reassembled products” to refer to the combined output of metatranscriptome assembly using the *euk* `rhythmic` pipeline. In brief, simulated raw reads created using the `jEUKebox` pipeline described below were trimmed, underwent quality estimation, and were assembled using multiple software tools which were identified or shown in previous studies to perform well with transcribed mRNA sequences, metagenomic data, or both. These assemblies were combined via multiple clustering steps and then functionally and taxonomically annotated with `eggNOG-mapper` and `EUKulele`, respectively (***Huerta-Cepas et al., 2017***; ***Krinos et al., 2021***). The details of the `jEUKebox` and *euk* `rhythmic` pipelines are expanded upon in the Materials and Methods (Section “Eukrhythmic pipeline”).

### jEUKebox pipeline generates simulated eukaryotic metatranscriptomes with varying sequence diversity

We developed the `jEUKebox` pipeline to facilitate the rapid and reproducible creation of comprehensive mock metatranscriptomic datasets that may be be used to benchmark pipelines and software. Here, we construct marine eukaryotic metatranscriptomes with differing sequence diversity and community complexity leveraging reference data from the MMETSP (***Keeling et al., 2014***; ***Caron et al., 2017***). We treat the `jEUKebox`-simulated datasets as a gold standard to assess the performance of the *euk* `rhythmic` pipelines and the assemblers that it uses (***Meyer et al., 2021***). More details about how the pipeline simulates raw reads that resemble the data type generated by marine metatranscriptomic surveys can be found in the Materials and Methods (Section “Simulation of eukaryotic communities using `jEUKebox`”). We randomly chose two distinct groups of laboratory transcriptomes from the MMETSP (***Keeling et al., 2014***) for the simulations to ensure that the results were not a product of the specific organisms we selected (Figure 2). The only part of the selection process that was not entirely random was requiring that some subset of the organisms that went into the communities had to have a highly similar partner in the same community per computed nucleotide similarity score (Section “Simulation of eukaryotic communities using `jEUKebox`”). We also designed the `jEUKebox` pipeline to include a balanced fraction of common transcripts that had an ortholog expressed by multiple organisms, and we implemented six distinct community configurations so as to simulate a range of species richness and evenness (Figure 2).

**Figure 1.**
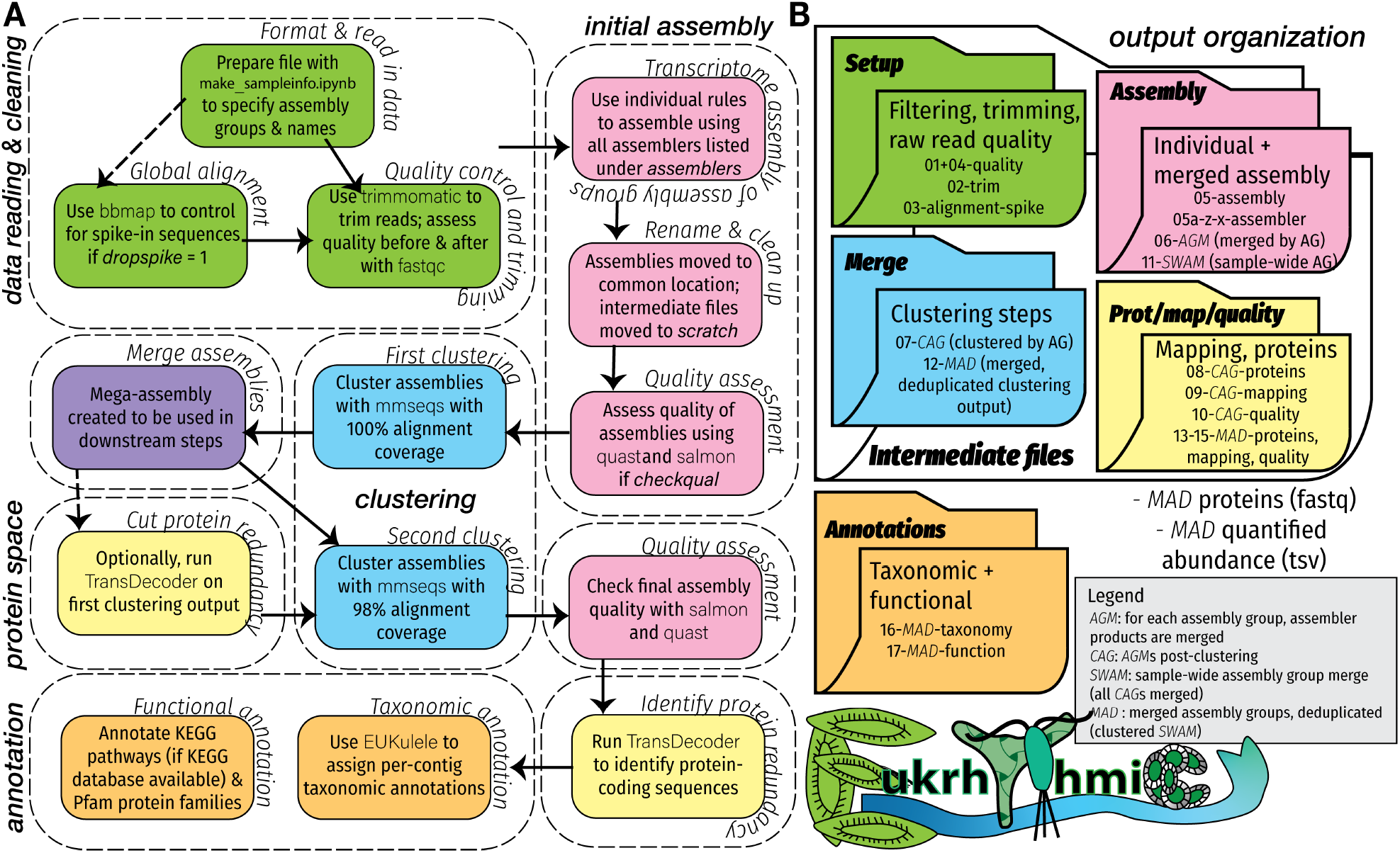
Conceptual diagram of *euk* `rhythmic` workflow, including (A) major and minor pipeline steps and (B) expected output of the pipeline.

**Figure 2.**
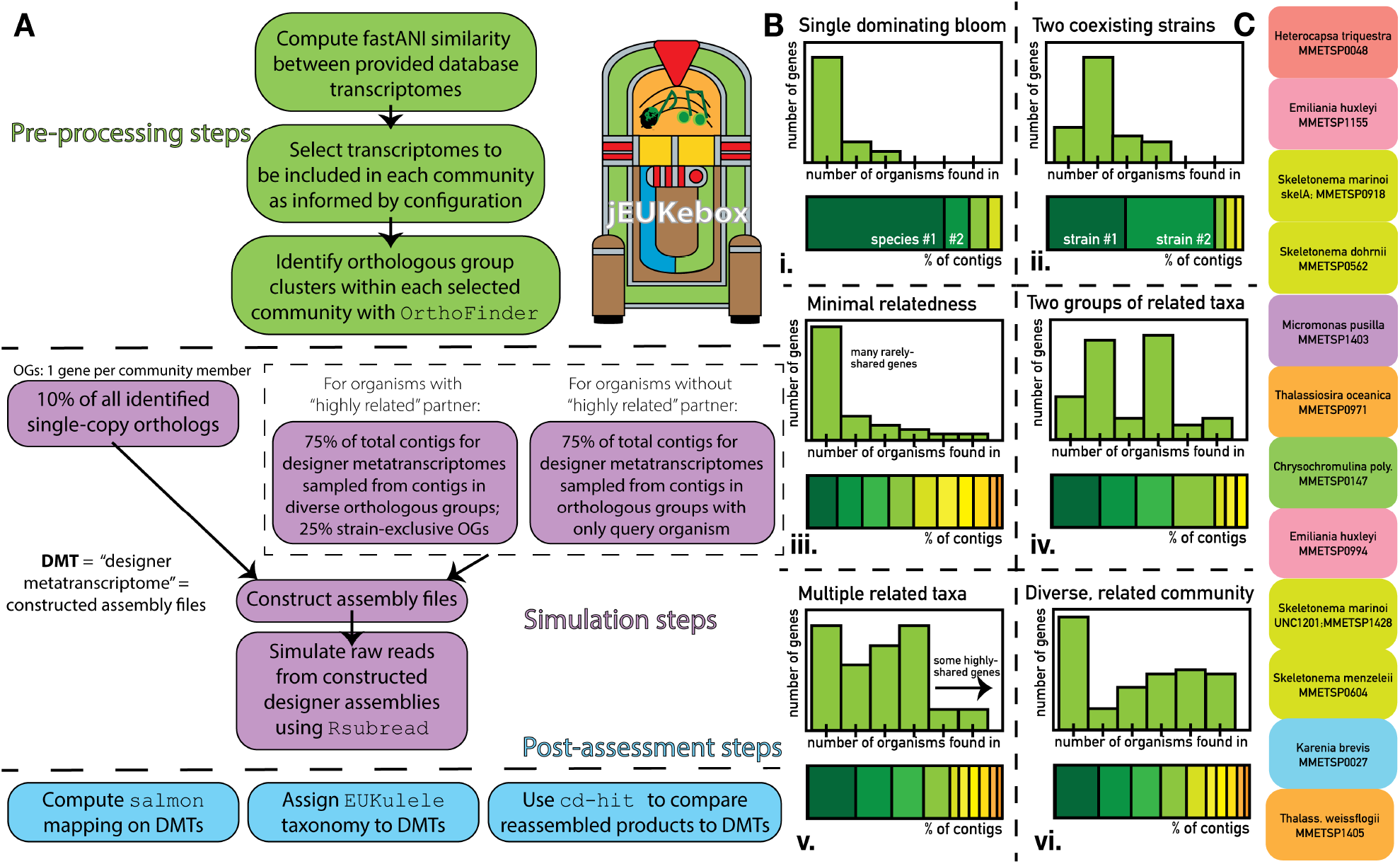
(A) the steps of the `jEUKebox` workflow for simulating environmental metatranscriptomes from individual transcriptomes from the MMETSP database. (B) Conceptual representation of the six targeted community composition simulations using MMETSP Group A from the paper.

### Eukrhythmic products accurately represent the raw reads

The *euk* `rhythmic` pipeline produced *de novo* assemblies that performed well as compared to the designer assemblies with respect to percentage mapping. The mapping of simulated raw reads against the *euk* `rhythmic` reassembled products was lower than against the designer metatranscriptomes against which they were simulated, with 87.5 ± 2.0% of simulated raw reads mapping against the *euk* `rhythmic` reassembled products and 96.0 ± 0.2% against the designer assembly (Figure 4). This discrepancy is likely due to the error introduction step in the raw reads or conflicts between different raw read placements in candidates for reassembled products which could not be resolved by the assembler. These patterns were reproduced in the environmental dataset we tested (***Alexander et al., 2015***): both the MAD (82.1±3.8%) assembly and the multi-assembler clustered assembly (“CAG”; 77.6 ± 4.5% mapped) outperformed any individual assembler with respect to percentage mapping (Figure 4E). In our simulated data, `rnaSPAdes` had the highest average percentage mapping of any assembler, and `MEGAHIT` had the lowest (Figure 4D), but patterns were slightly different in the environmental dataset (***Alexander et al., 2015***). While `MEGAHIT` still underperformed relative to the other assemblers with respect to percentage mapping (Figure 4D-E), comparisons between the remaining assemblers were less straightforward. `rnaSPAdes` showed the highest individual performance (75.0 ± 4.7% mapped), followed by `SPAdes` (70.5 ± 5.4% percent mapped). However, `Trinity` performed better in some samples than in others, hence showed higher dispersion in percentage mapping values (67.8 ± 8.6%).

**Figure 3.**
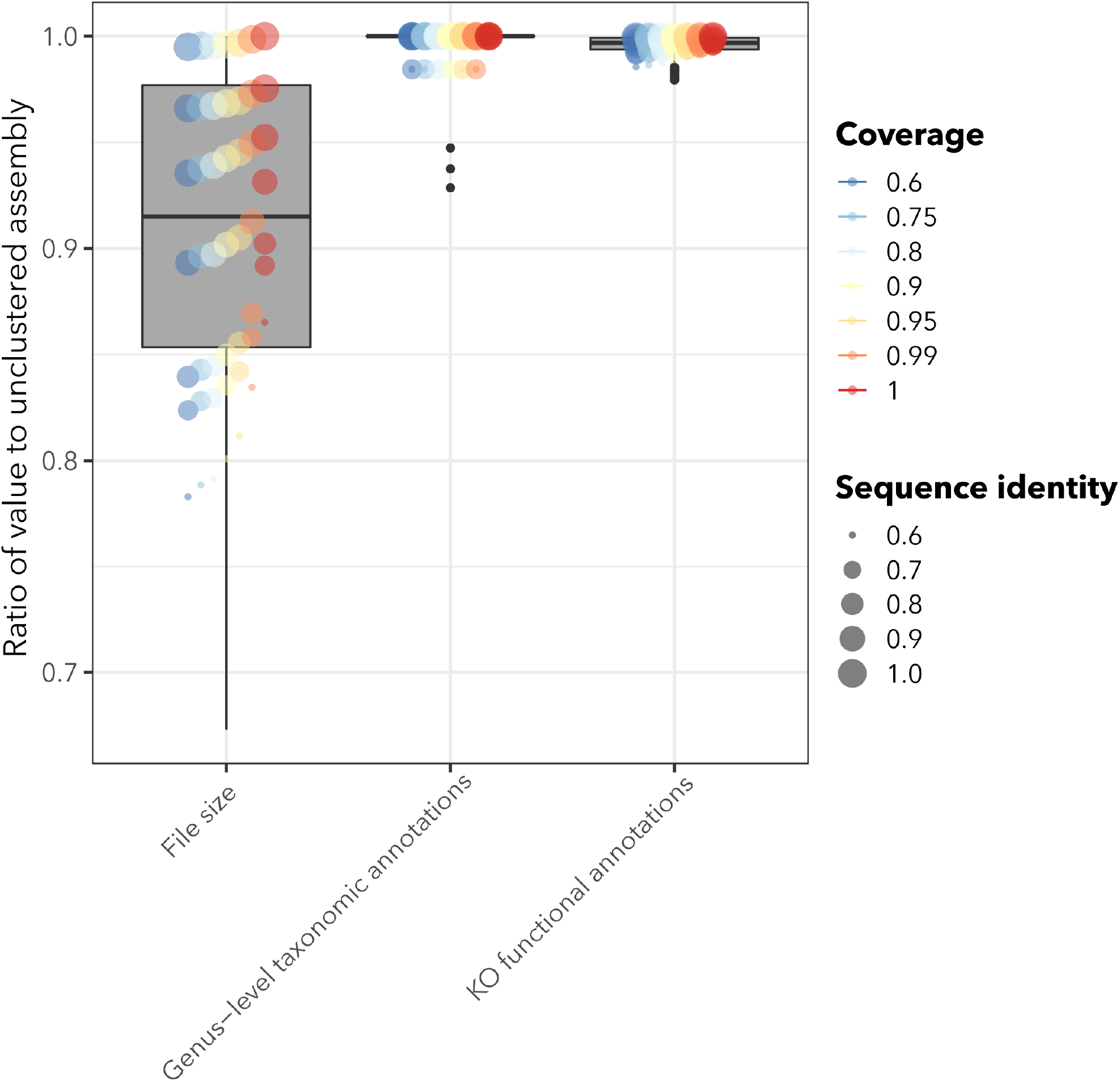
Effect of clustering designer assembly on assembly size and annotations. Clustering was performed on the original “designer metatranscriptome” set of contigs from the MMETSP references using the `mmseqs2` tool ***Mirdita et al. (2019)***. Eukrhythmic uses a coverage level of 0.98 and sequence identity of 1 for `mmseqs2` clustering. See Supplementary Figure 1 for a more detailed graphical summary of the influence of sequence identity and coverage on the size of the recovered assembly and its functional and taxonomic annotations.

**Figure 4.**
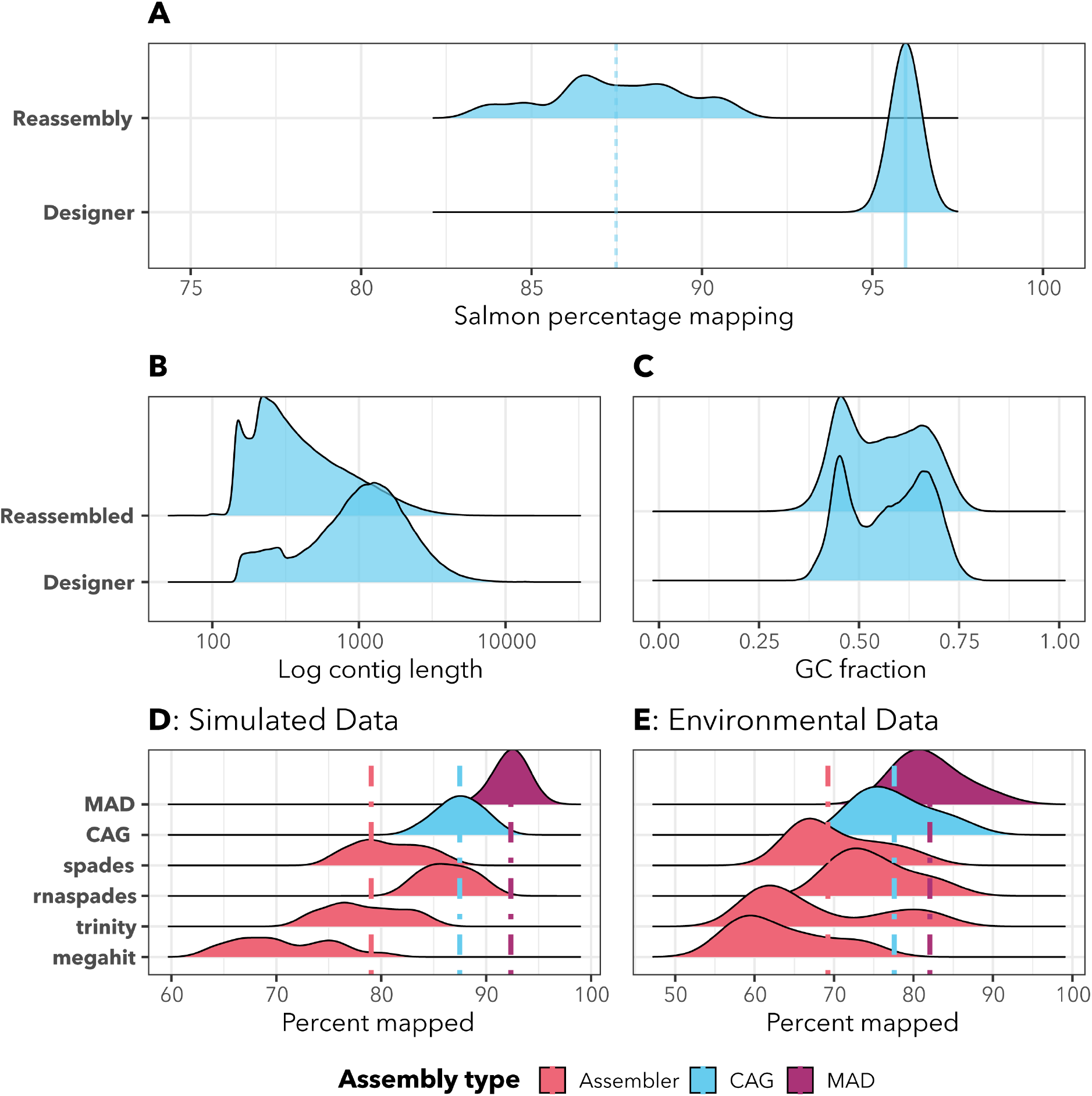
Basic assembly statistics for the eukrhythmic reassemblies (per-sample) as compared to the designer metatranscriptomes. A: `Salmon` percentage mapping distribution for the designer vs. reassembled metatranscriptomes. B: Log-normalized contig length distributions compared between designer and reassembled. C: Per-sequence fraction of GC-content for the designer as compared to the reassemblies. D: Environmental data from Narragansett Bay as a comparison to Panel C.

Average contig length tended to be significantly shorter in the eukrhythmic reassemblies as compared to the designer metatranscriptomes, although there was considerable variability (Figure 4). The average length of open reading frames (ORFs) predicted by the `TransDecoder` tool was also smaller in the *euk* `rhythmic` reassemblies as compared to the original sequences retrieved from the MMETSP transcriptomes (Table 1; Figure 4). Although still substantially shorter than the designer metatranscriptomes, sequences in the *euk* `rhythmic` products that were recovered by more than one assembler according to `mmseqs2` clustering had progressively longer length (median length 334 base pairs for clusters represented by a single assembler, median length 960 base pairs for clusters represented by all four assemblers; between-distribution t-test *p <* 0.001; Figure 4). These longer contigs had high fidelity with the raw reads, as evidenced by the agreement of multiple assembly approaches, hence were likely to be longer sequences interrupted by fewer conflict instances.

**Table 1.** Comparison of the average length of sequences in the designer metatranscriptomes as compared to the *euk* `rhythmic` reassemblies. Both the average length of nucleotide sequences and protein sequences as predicted by `TransDecoder` are provided, as well as the average fraction of GC-content for nucleotide sequences.

| Assembly | Sequence Type | Average Length | Average Fraction GC Content |
| --- | --- | --- | --- |
| Designer Assemblies | Nucleotide | 1278.7 ± 1027.4 | 0.57 ± 0.10 |
| Reassembled Products | Nucleotide | 539.9 ± 564.3 | 0.56 ± 0.10 |
| Designer Assemblies | Protein | 276.3 ± 261.7 | - |
| Reassembled Products | Protein | 165.1 ± 157.7 | - |

### Less stringent clustering slightly decreases identified annotations

*euk* `rhythmic` reduces the redundancy of the identified contigs for the merged assembly via clustering, hence reducing computational complexity of downstream operations on the smaller multi-assembler, multi-sample assembly file. Applying clustering directly to the designer metatranscriptomes revealed that substantial protein-space clustering only slightly decreases the unique annotations extracted from the dataset. Using a sequence identity threshold of 0.6 and a coverage threshold of 0.6 in coverage mode 1 using `mmseqs2` reduced the number of contigs in the assembly by an average of 23.7% and reduced the assembly file size by an average of 21.7%, but only reduced average identified KEGG database functional annotations by and average of 1.4% and did not result in the loss of any species from the dataset via clustering (Figure 3). By default, *euk* `rhythmic` uses a conservative approach of 100% sequence identity and 98% coverage for the most lenient clustering step, but we found in this test that values of 80% for both coverage and sequence identity could considerably reduce total file size without considerably changing unique annotations (Supplementary Figure 1). Given this substantial reduction in file size without loss of the majority of annotations, more stringent clustering thresholds may be warranted, especially in datasets with many samples or high sequencing depth.

### Eukaryotic metatranscriptome assembly accurately recapitulates simulated taxonomic diversity

On the whole, all assemblers performed well with respect to recovery of the major taxonomic annotations from the simulated metatranscriptomes. 94.8±2.2% of all recovered contigs were assigned genus-level annotations by the `EUKulele` tool that matched genera found in the selected MMETSP transcriptomes used to simulate the metatranscriptomes (97.7±2.2% of annotated contigs). In general, the number of annotations in conflict with the genus-level annotation assigned based on the MMETSP was similar in the designer metatranscriptomes as compared to the reassemblies generated by eukrhythmic. The total abundance of each genus-level annotation as assessed by `Salmon` quantification matched well between the designer metatranscriptomes and the reassembled products from eukrhythmic. The computed linear regression between the genus-level annotations from the designer assemblies and the eukrhythmic re-assemblies was nearly one-to-one: Reassembly = −1353 + 1.02(Designer); *R* = 0.95; *p* =*<* 8.2*e* − 184; note that the intercept is relative to total abundances on the order of 10^5^.

Despite this performance, some genus-level annotations were missing based on the contigs provided from the MMETSP. Between all trials, an average of 1.3 ± 1.9 genera (averaged per-trial integer numbers) out of an average of a total 6.1 ± 2.7 genera were not recovered by the *euk* `rhythmic` re-assembly, despite being present in the MMETSP transcriptomes which were used to create each community (see Table 4). As many of these annotations were also missing from the `EUKulele` annotations on the contigs from the MMETSP themselves (1.9±1.9 genera), these contigs may simply have not been sufficiently distinct from the transcriptomes of other organisms in the database to be annotated, potentially due to sequence length or specificity. Using the `EUKulele` annotations instead of the taxonomic annotation of the transcriptome from which the original contigs were taken, 2.8 ± 1.7 genera were not found in the eukrhythmic reassembled results as compared to the original `EUKulele` annotations of the designer assemblies. An average of 39.3 ± 12.9 distinct genus-level annotations were assigned in the eukrhythmic output as compared to 6.1 ± 2.7 distinct MMETSP genera being used to generate the samples due to being successfully annotated as similar genera present in the MMETSP. These spuriously taxonomically-annotated contigs constituted both a minority of the total assembled contigs as well as estimated abundance from the simulated raw reads (Supplementary Figure 5), and the occurrence of these spurious annotations could be reduced with more stringent `EUKulele` parameters, though at the expense of some correct annotations.

**Table 2.**
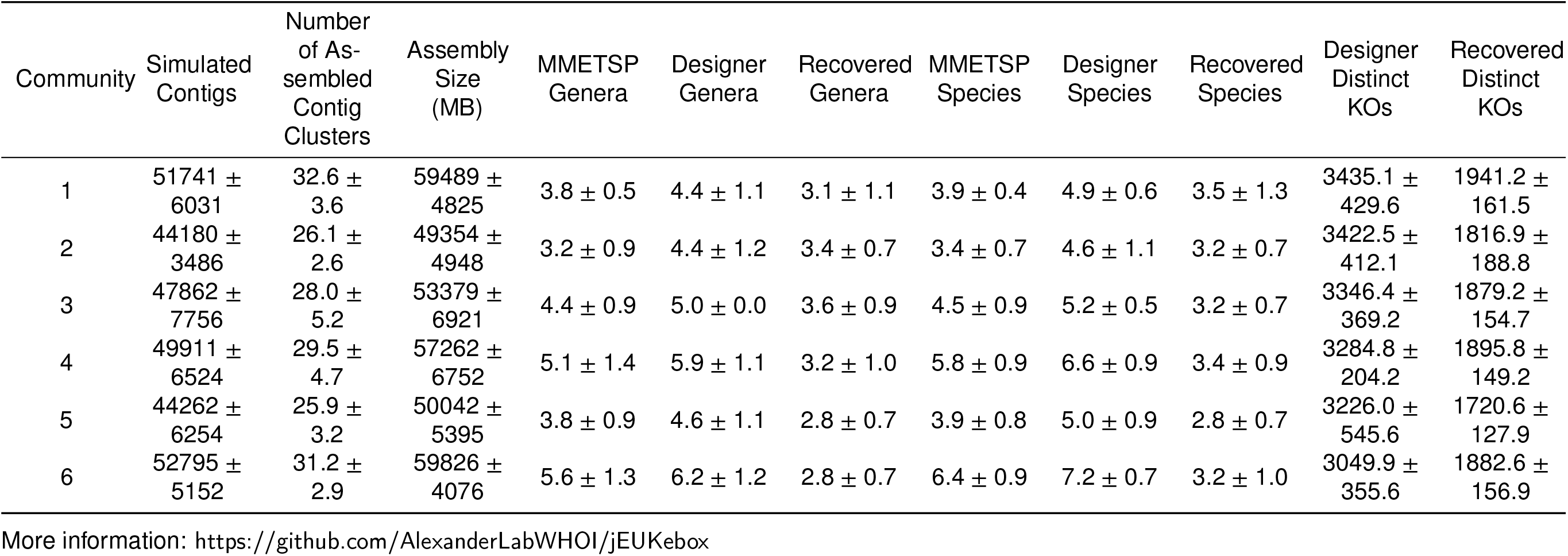
Resulting assembly size and taxonomic, functional, and core content recovery of `jEUKebox` outputs after raw read simulation and re-assembly with *euk* `rhythmic`. Four assemblers were used in this analysis which were then clustered together using default *euk* `rhythmic` settings. The mean and standard deviation of four trials of each community and list of MMETSP IDs is presented. We also show the number of genera that were (1) originally included via transcriptomes leveraged from the MMETSP (2) identified using `EUKulele` within the simulated metatranscriptomes and (3) recovered in the reassembled data after application of the *euk* `rhythmic` pipeline. For functional KO IDs, only the designer assemblies and the *euk* `rhythmic` reassembled products could be compared. A version of this table in which the two distinct communities designed from the MMETSP (combining the two contributes to relatively high standard deviation) are presented separately is provided as a supplement.

**Table 3.** Calculated diversity metrics for the six simulated MMETSP-based communities used in the analysis. The `sourmash` composite score is an abundance-weighted average of the `sourmash` distance between two MMETSP transcriptomes. The Shannon diversity index is computed according to ***Shannon (1948)***, and the richness is the number of MMETSP transcriptomes included in the community metatranscriptomes (species richness).

| Community | *sourmash*<br>composite<br>score | Shannon | Richness |
| --- | --- | --- | --- |
| 1 | $0.9 \pm 10^{-16}$ | $15.8 \pm 0.2$ | 4 |
| 2 | $1.3 \pm 10^{-17}$ | $18.3 \pm 0.7$ | 5 |
| 3 | $2.0 \pm 10^{-16}$ | $28.3 \pm 0.7$ | 8 |
| 4 | $2.3 \pm 0$ | $36.6 \pm 0.5$ | 10 |
| 5 | $1.8 \pm 10^{-17}$ | $25.2 \pm 0.7$ | 7 |
| 6 | $2.4 \pm 10^{-16}$ | $35.6 \pm 0.5$ | 12 |

The sequence annotations were classified according to whether they did or did not align with genus-level annotations from the MMETSP (Figure 5). There was no statistically significant difference between the per-sample summed abundance of incorrectly-annotated contigs between the designer and the *euk* `rhythmic` reassembled products (T=-0.084; *p*=0.93), however correctly-annotated contigs were significantly more abundant in the designer assemblies (T=-5.28; *p*=8.3e-7) and unannotated contigs were significantly more abundant in the *euk* `rhythmic` reassemblies (T=5.43; *p*=4.5e-7).

**Figure 5.**
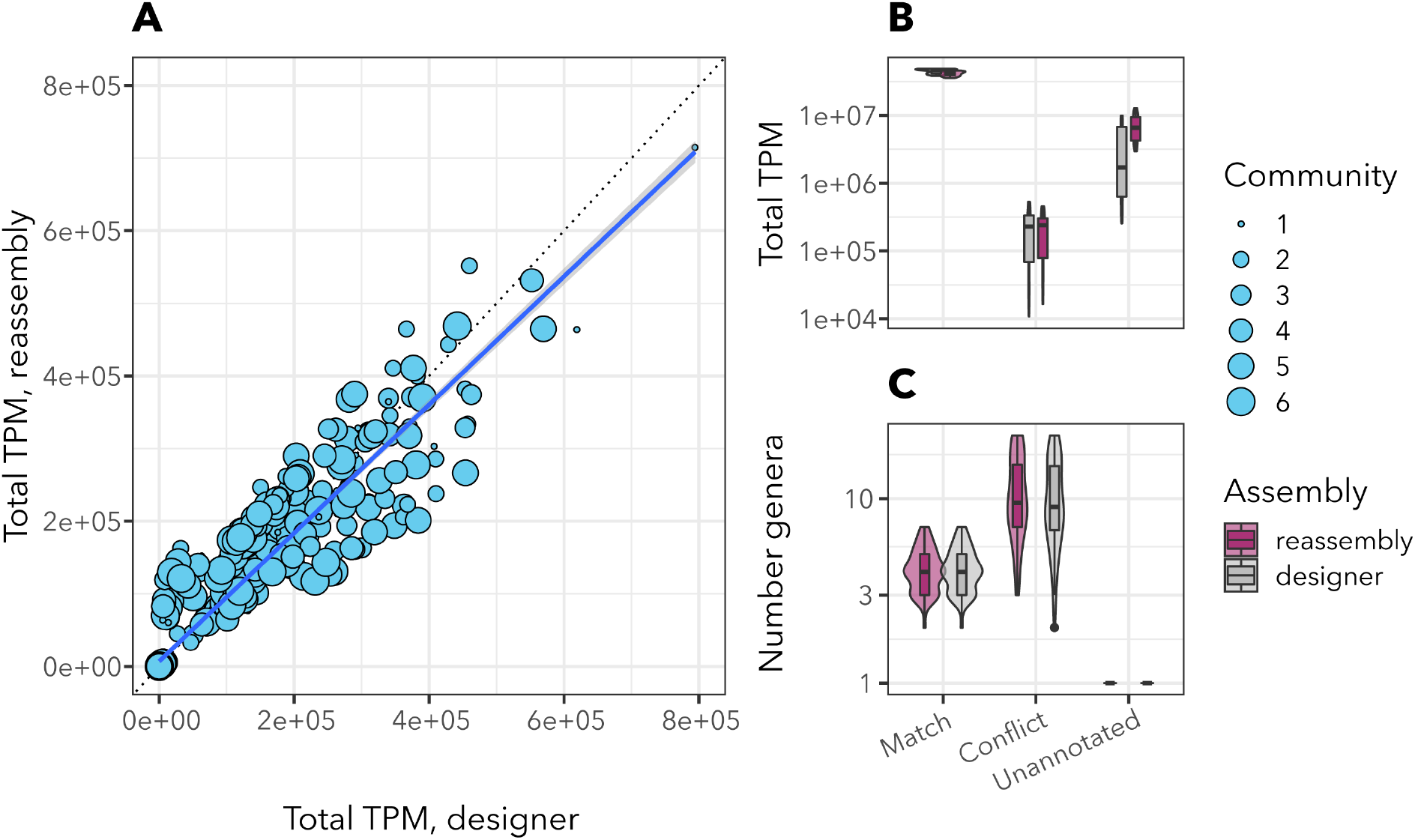
Visual summary of findings of taxonomic annotations from eukrhythmic. A: summed transcripts per million (TPM) as reported by `Salmon` from the designer assembly compared to the eukrhythmic reassembly. Each point represents a genus; the dotted line is a 1-to-1 line (*y* = *x*), or collection of reference transcriptomes from the MMETSP. Circle size corresponds to community type as described in the text; of note is that some communities have very highly abundant genera, such as the smallest circles corresponding to Community 1. B: Sum of total TPM in the designer vs. reassemblies that corresponded to genera which (1) matched genera from the original MMETSP transcriptomes, (2) conflicted or did not match genera from the original MMETSP transcriptomes, or (3) were not annotated, according to `EUKulele`. C: The number of genera that matched (true positives), did not match (false positives), or were unannotated (false negatives according to database precision). As shown in panel B, unannotated contigs at the genus level were more abundant in the reassemblies than the designer metatranscriptomes. There were also statistically significantly more matches in the designer metatranscriptomes than the reassemblies from eukrhythmic. However, false positives were occurred at a similar rate between the two assembly types, indicating that these were more likely a product of the original quality of the contigs from the MMETSP or their ability to be uniquely classified.

### Functional annotations from metatranscriptome assembly match abundance and diversity of functions in designer transcriptomes

Functional annotations were recovered with similar frequency and relative abundance in the *euk* `rhythmic` reassembled products as compared to the designer assemblies (Figure 6). As an overall average across MMETSP groups and samples, 5820.6±349.6 KEGG orthology terms (KOs) were correctly recovered from the designer assemblies, 820.3±163.7 were “false positives” that were recovered in the eukrhythmic reassemblies but not in the original designer assemblies, and 473.8±107.6 were identified in the designer assembly but not recovered by *euk* `rhythmic`. However, the false positive and unrecovered KOs tended to have low abundance compared to those which were correctly identified: on average, there were 1566.5±321.3 total occurrences of annotations of false positive KOs per sample in the eukrhythmic reassemblies and 107.6±204.7 total occurrences of annotations of KOs that were not found in the eukrhythmic reassemblies in the designer assemblies, as compared to and average of 132751.9±10176.5 occurrences in the designer assembly and 116489.5±9961.0 occurrences in the eukrhythmic reassemblies of KOs that were mutually recovered before and after the reassembly process. A linear regression with an imposed *y*-intercept of zero as calculated in R (***R Core Team, 2021***) revealed a relationship of Reassembled KO abundance = Designer KO abundance · 0.96 with an adjusted *R*^2^ of 0.85 (*p* = 2.2*e*^−16^), indicating a nearly one-to-one relationship between the abundances of each KO in the designer assembly and in the reassembled products (including false positives and KOs missing from the eukrhythmic reassemblies; Figure 6).

**Figure 6.**
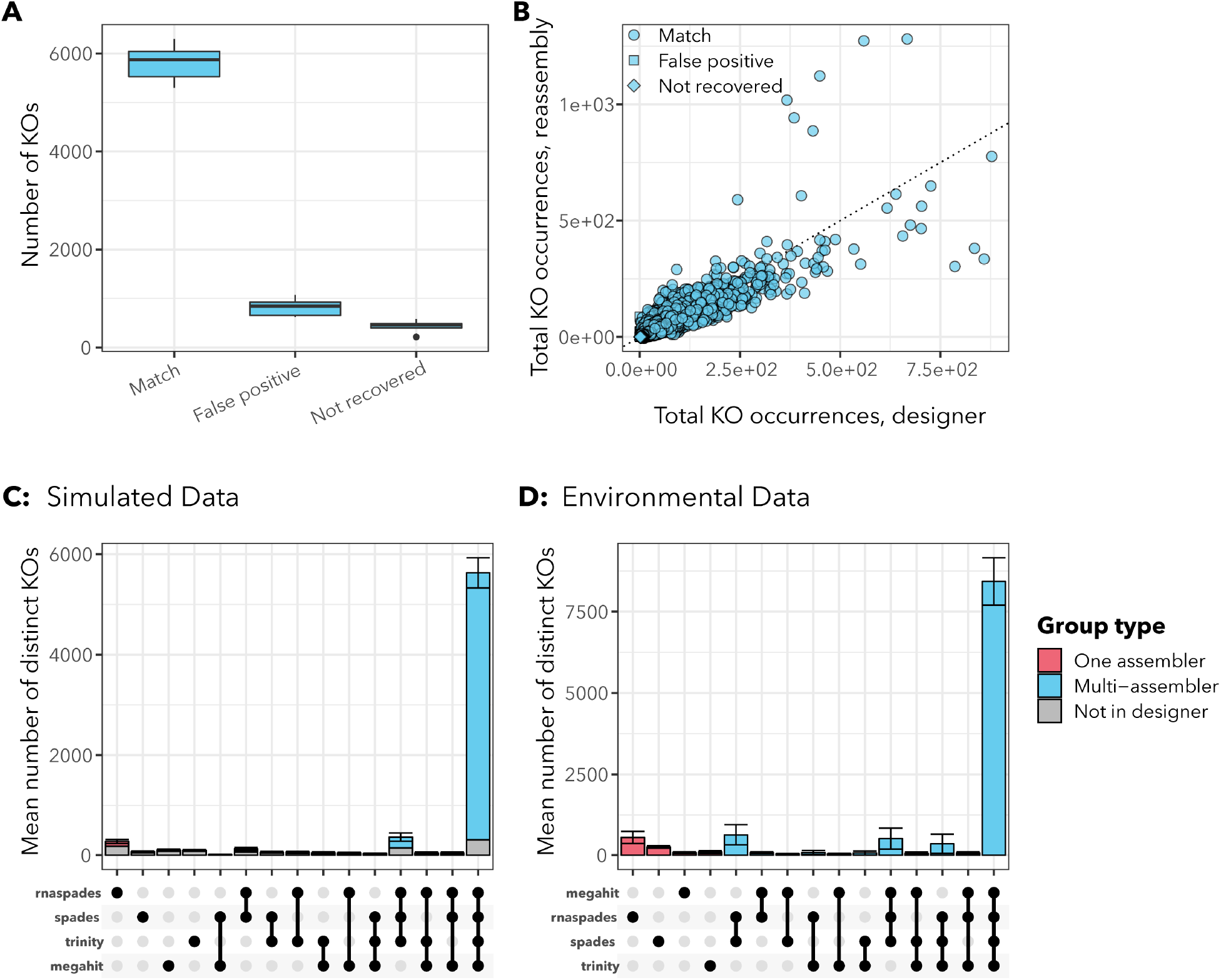
Visual summary of functional annotation findings from reassembling the simulated raw reads from the designer metatranscriptomes. A: on the far left, the total number of KEGG orthology terms (KOs) recovered from the designer metatranscriptomes that were also identified in annotations from the designer transcriptomes (“Match”) is shown. The other two boxes represent total number of KOs that were found in the eukrhythmic reassemblies despite not actually being present in the original designer assemblies (“false positives”) and those that were present in the designer assemblies but not recovered via *euk* `rhythmic` (“not recovered”). B: the number of occurrences of each KO is compared between the designer metatranscriptomes (horizontal axis) and the *euk* `rhythmic` reassemblies (vertical axis). C: this incidences of each KO in the designer assemblies and the *euk* `rhythmic` reassemblies are broken up by the individual assemblies that each KO was recovered from (“incidence count” is the number of KOs meeting each category). The majority of all recovered KOs are shown to be recovered by all four assemblers as well as present in the designer metatranscriptomes. Portions of the bar colored in gray indicate that these KOs were recovered by all of the assemblers listed, but were not found in the desinger assembly. D: Environmental data for KOs from Narragansett Bay as a comparison to Panel C.

The vast majority of KOs also recovered in the designer assemblies were identified by all four assembly tools (5326.6±247.9 KOs across all samples). `rnaSPAdes` individually recovered the highest number of unique KOs that were also found in the designer assembly of any assembler (96.0±20.4), but `rnaSPAdes` also generated the highest number of KOs that were not found in the designer assemblies (176.4±26.5), nearly double the number that it uniquely recovered (Figure 6).

### Applying the *euk*rhythmic pipeline to environmental metatranscriptomic datasets

To benchmark the *euk* `rhythmic` pipeline and to provide examples of the potential biological insights that can be extracted from the assembly approach, we assembled and annotated samples from two metatranscriptomic datasets. First, we chose two sets of samples from the *Tara* Oceans project as a representative general oceanographic dataset: one set from the Southern Ocean and one from the Mediterranean Sea, two ocean basins with contrasting levels of diversity. We found that *euk* `rhythmic` expands the total amount of protistan coding sequence data that can be recovered from anywhere in the global ocean. We also assembled a metatranscriptomic dataset from a previously published study in Narragansett Bay as a coastal example with a dominant taxonomic group (i.e. a canonical “bloom” scenario). We observe that while *euk* `rhythmic` recapitulates many of the general patterns from a direct read mapping-based study, the assembly approach outperforms direct read mapping with respect to the number of distinct diatom representatives recovered, the dominant taxonomic group (*Bacillariophyta*) in the samples.

### *Tara* Oceans *euk* rhythmic assemblies contain coding sequences that lack representation in the “MATOU” gene atlas

More than one in six (17.5%) of the coding sequences we recovered from our multi-assembler *euk* `rhythmic` assemblies from the *Tara* Oceans metatranscriptomes had no significant hits to the “MATOU” composite gene atlas across all *Tara* Oceans metatranscriptomes curated by ***Carradec et al. (2018)*** (Figure 9). An average of 16.1% of all Mediterranean Sea coding sequences and 18.8% of all Southern Ocean sequences did not have any match to previously recovered coding sequence content in the MATOU database, which includes coding sequences from all major global ocean basins. These results indicate the expansion of coding sequences achieved by using *euk* `rhythmic`, but also that total number of coding sequences is not evenly expanded across samples (Figure 9E); while in some samples >75% of coding sequences did not have a match to the MATOU database, in others it was <10%.

**Figure 7.**
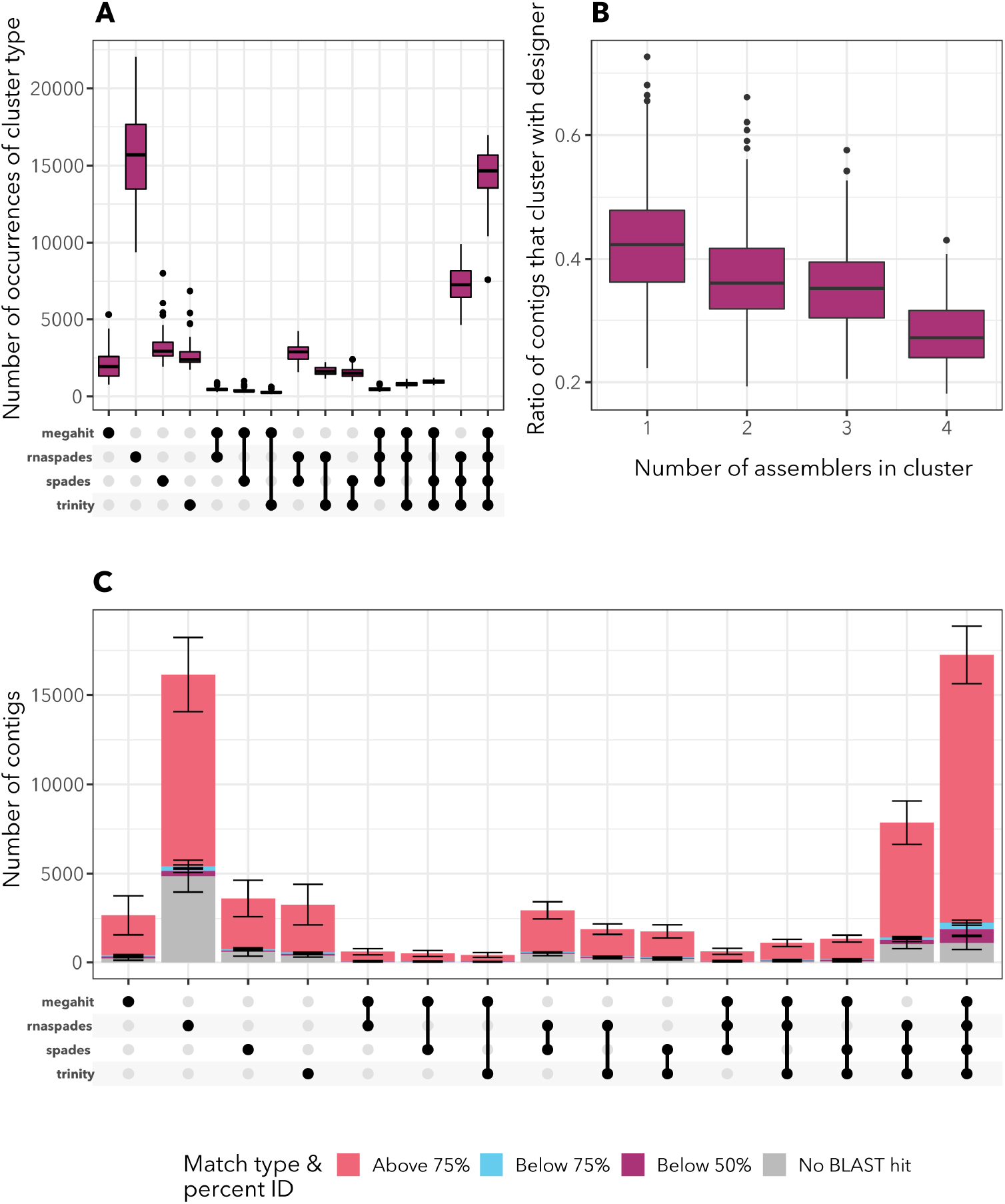
Summary of `mmseqs2` clustering within eukrhythmic to collapse redundant sequences and then secondarily to compare extracted ORFs to the designer assembly proteins, and `BLAST` hit matching to assess the similarity of extracted protein content from the reassemblies to the original designer contigs. Panel A: total number of contigs per sample as separated by the assemblers that were recovered from. `rnaSPAdes` produced the highest number of contigs overall independently, which was a higher overall number than the contigs which were produced by all four assemblers (far right boxplot in panel A). Panel B: The ratio of `mmseqs2` clusters of proteins that did versus did not cluster with proteins from the designer assembly as a function of the number of assemblers the protein product was found by. Protein products supported by assembly by all four assemblers were least likely to be “spurious”, or not recoverable from the designer assembly. Panel C: Number of contigs that had no protein ORF assigned to them via `TransDecoder` (black) as compared to contigs with proteins having `BLAST` matches according to some percentage identity. The first stacked bar corresponds to contigs that both had a detected ORF and a BLAST match with percentage identity *>*75% at an e-value threshold of 10^−2^. Supplementary Figure 7.4 shows the contigs from the designer assembly which originally did not have an identified ORF.

**Figure 8.**
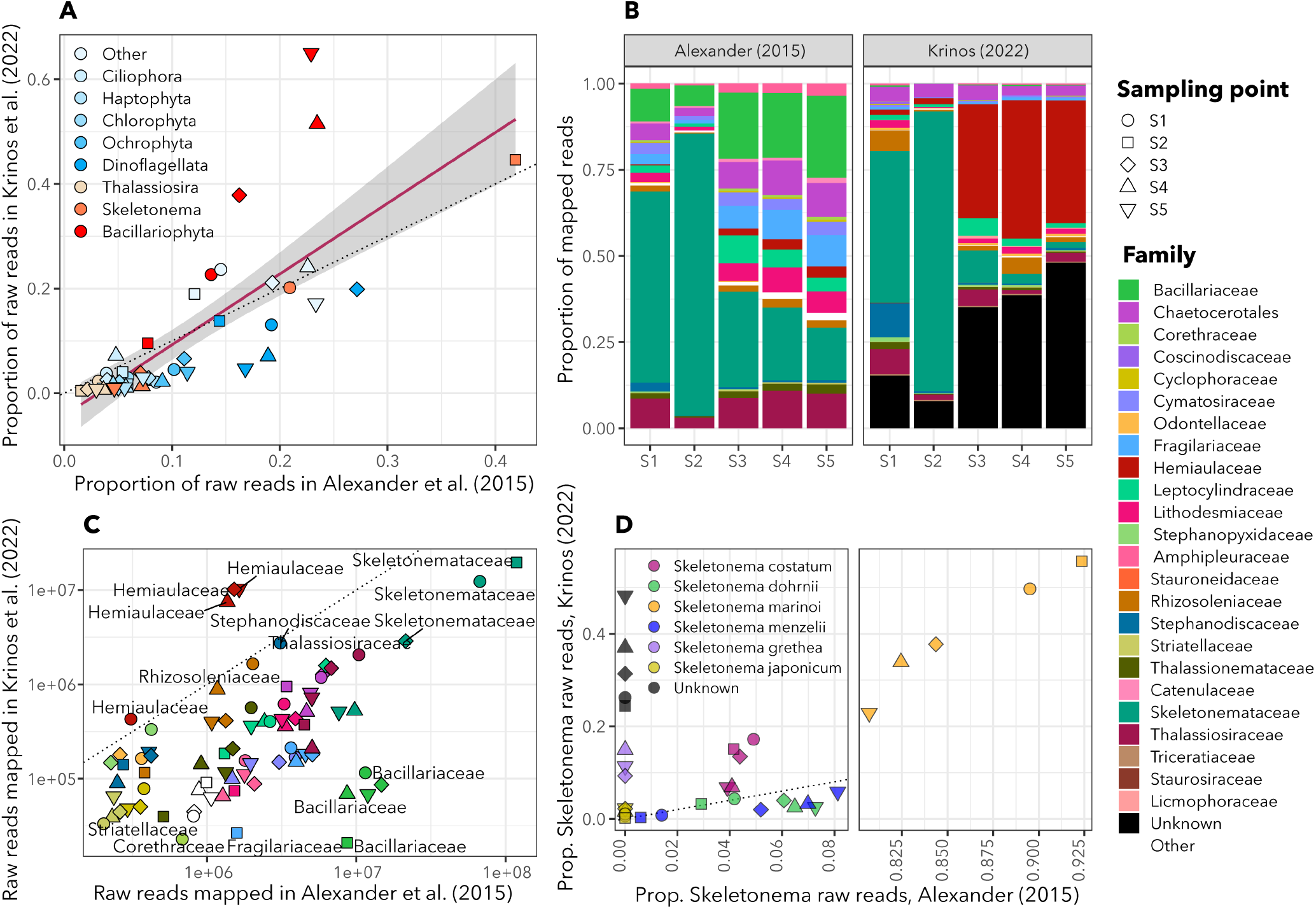
Narragansett Bay dataset from ***Alexander et al. (2015)*** assembled using *euk* `rhythmic`. A: The correspondence between the proportion of total raw reads in (y) this study vs. (x) ***Alexander et al.*** (***2015***). Each point represents a sampling time, and *Bacillariophyta* aggregates all non-*Skeletonema* and non-*Thalassiosira* diatoms. B: Family-level taxonomic breakdown of ***Alexander et al.*** (***2015***)’s raw read mapping (left) as compared to this study. C: Log-normalized raw reads mapped to each taxonomic family compared between the two studies. D: *Skeletonema* species represented in the *euk* `rhythmic` reassembly representing some of the diversity within this genus known to show seasonal dominance in Narragansett Bay.

**Figure 9.**
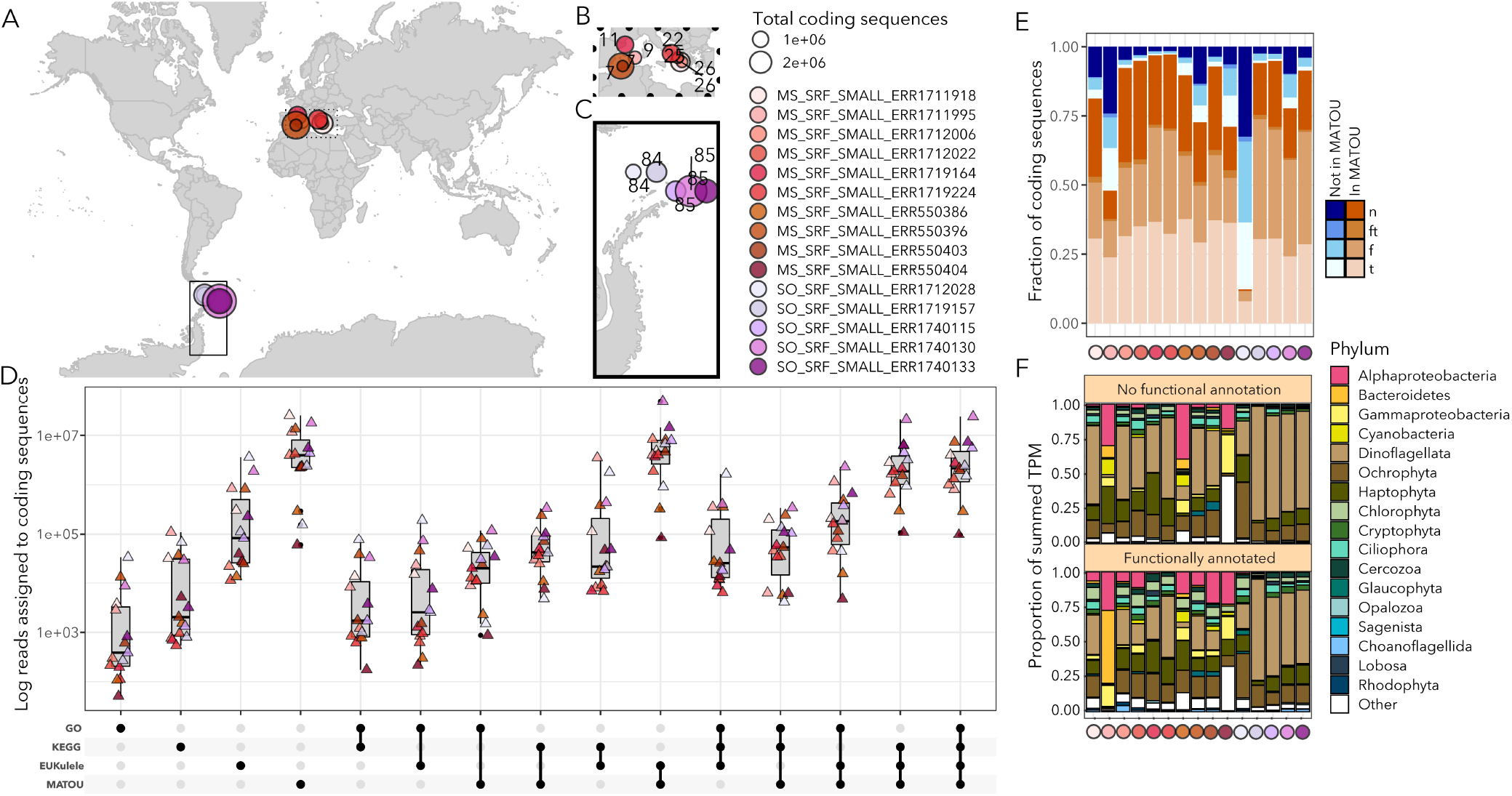
*Tara* Oceans reassemblies using *euk* `rhythmic`. A: Map showing the locations of reassembled *Tara* Oceans samples. Boxes over regions are expanded in Panels B and C. B: Mediterranean Sea samples. Numbers indicate *Tara* Oceans stations. C: Southern Ocean samples. As in Panel B, numbers indicate *Tara* Oceans stations. D: between-assembler overlap of the reads assigned to coding sequences. The x-axis indicates the annotations assigned to each of the coding sequences, and the y-axis shows the between-sample sum of reads assigned to coding sequences for that category. E: Fraction of coding sequences that did or did not have a match to the MATOU database. Shades of blue indicate coding sequences recovered only by this study. The top segment indicates coding sequences without functional or taxonomic annotations, following by the proportion of sequences with functional and taxonomic annotations (“ft”), the proportion with only functional annotations (“f”), and the proportion with only taxonomic annotations (“t”). The same is shown in shades of orange for the assembld coding sequences from this study that did have a significant match to the MATOU database. The y-axis shows the color-coded *Tara*Oceans sample. F: The fraction of TPM assigned to coding sequences with recovered taxonomic annotations. These are from the “Not in MATOU” “ft” and “t” bars in Panel E. Dinoflagellated dominate many of the Southern Ocean samples, particularly for those coding sequences which could not be taxonomically annotated.

As much as 41.3% of the coding sequence products of the assembly for each sample could not be assigned a taxonomic annotation via `EUKulele`, but more so in samples from the Mediterranean Sea (Supplementary Figure 9.2); mean Mediterranean Sea: 34.8 ± 5.2%, mean Southern Ocean 28.0 ± 4.4%). Among the fraction of sequences that had `EUKulele` taxonomic annotations and were not found in the MATOU database, dinoflagellates dominated the recorded number of coding sequences recovered in both basins (mean Mediterranean Sea: 12.0 ± 7.7%, mean Southern Ocean: 34.0 ± 10.1%; Figure 9F). Dinoflagellates also dominated the taxonomically-annotated coding sequences in terms of average proportion of total TPM (SO: 30.3±11.1%; MS: 8.6±6.6%), but not in terms of mean raw TPM assigned in the Southern Ocean (SO: Ochrophytes had the highest assigned TPM at 32853.1±66098.9, while dinoflagellates had 31853.4±82808.6 assigned TPM), because some samples dominated by dinoflagellates also had a relatively low number of reads assigned to coding sequences not found in the MATOU database (Figure 9E-F). All of the taxonomic annotation information for the fraction of the *euk* `rhythmic` sequences that had a `EUKulele` annotation is summarized in Supplementary Figure 9.3.

Our efforts expand the total coding sequence content available from global ocean metatranscriptomes, but also highlight the ongoing need for intercomparison of approaches. The average length of coding sequences that did not have a match to the MATOU database was 466.9±243.1 bases, while the average length of the coding sequences that did have a match was 613.5±438.0 bases. The average length of the coding sequences with a match (i.e., recovered by both assembly efforts) was significantly longer (*t*=720.86; *p*<1e-16). The use of a *k*-mer size of 63 with the `velvet` (***Zerbino and Birney, 2008***) assembler by ***Carradec et al. (2018)*** may also have contributed to this result - the `rnaSPAdes` assembler, for example, takes into account the varying coverage level of expression data by using a *k*-mer size that varies dynamically with read length (***Bushmanova et al., 2019***). This approach occasionally increases misassembly rate, but also protects rarely-expressed genes from being missed (***Bushmanova et al., 2019***). Because eukaryotic communities in the microbial ocean may be sparse and contain rare taxa, we argue that a more exhaustive assembly approach is warranted, even if the average length of assembled sequences is reduced.

### *Tara* Oceans assembly highlights potential importance of non-coding sequences

*Tara* Oceans assemblies from the Mediterranean Sea and Southern Ocean varied in their overall composition as well as the accuracy of their recovery in the assembly process. While we focused the balance of our analysis on predicted coding sequences to compare to the analysis by ***Carradec et al. (2018)***, we note that via `Salmon` mapping, an average of 30.1±10.7% of the raw reads for the Mediterranean Sea samples mapped back to the coding sequences, as compared to 51.5±13.3% for the full assembly, while in the Southern Ocean samples, 51.5±11.6% of the raw reads mapped back to coding sequences as compared to 76.4±10.3% for the full assembly (Supplementary Figure 9.1). This indicates that in both cases, a substantial fraction (*>*20%) of the original raw reads can be assembled into contigs, but appear to be non-coding. These non-coding sequences may be involved in important regulatory processes (***Budak et al., 2020***; ***Rogato et al., 2014***), such as nutrient stress in diatoms (***Lopez-Gomollon et al., 2014***), hence should not be excluded from consideration.

### Multiple assemblers improve metatranscriptome assembly of Narragansett Bay phytoplankton

We benchmarked the *euk* `rhythmic` pipeline using a previously analyzed marine metatranscriptomic dataset (***Alexander et al. (2015)***; Figure 8). In particular, we were able to recapitulate the taxonomic composition of the diatom-dominated community described in ***Alexander et al. (2015)***. Across all assemblers, representative from the phylum *Ochrophyta* were suggested to be dominant members of the community (Figure 8), and moreover the genera *Skeletonema* and *Thalassiosira* were recovered in expected proportions, with Skeletonema producing a numerically-dominant bloom determined via cell counts obtained from microscopy in sample S2 (Supplementary Figure 8.2). Notably, our assembly recovered a greater diversity of diatom species than the raw read mapping method used previously (Figure 8; ***Alexander et al. (2015)***), including the recovery of multiple species of *Skeletonema* known to be present in this ecosystem (***Canesi and Rynearson (2016)***; Figure 8D).

While broad patterns in taxonomic annotations were indistinguishable between the different assemblers and the majority of KEGG Orthology (KO) IDs were recovered by all four assemblers (Figure 8B,C), the assemblers showed some differences with respect to the abundance of each functional annotation. In particular, `MEGAHIT` reported fewer instances of each functional gene grouping than `rnaSPAdes`, and fewer than `Trinity` approximately half of the time (Figure 8; Supplementary Figure 8.1). rnaSPAdes appeared to report a lower overall abundance of diatoms when the normalized TPM metric returned by `Salmon` ***Patro et al. (2017)*** was used, but this pattern did not hold when non-normalized raw reads were used instead (Figure 8; Supplementary Figures 8.2-4). Contigs generated by the assemblers that were successfully annotated as *Skeletonema* or some other diatoms appeared to have longer average length than the average among all taxa (mean length of *Skeletonema* contigs with standard error of the mean: 618.6 ± 0.7; overall mean: 396.6 ± 0.07; two sample t-test t = 310.17 *p <* 2.2*e* − 16; Supplementary Figure 8.5). `rnaSPAdes` produced a disproportionately high number of contigs relative to the other assemblers, many of these contigs belonging to non-diatom taxa (Supplementary Figure 8.6). These contigs also tended to be shorter in the `rnaSPAdes` assemblies, both for non-diatom taxa (`rnaSPAdes` mean ± standard error = 377.7 ± 0.1; overall mean = 421.1 ± 0.09; t-test t=-367.17, *p <* 2.2*e* − 16) and for unannotated contigs (`rnaSPAdes` mean ± standard error = 264.7 ± 0.7; overall mean = 300.8 ± 0.07; t-test t=-498.5, *p <* 2.2*e* − 16). While these differences were universal for annotated non-diatom and unannotated contigs, `rnaSPAdes` produced shorter *Skeletonema* contigs than `Trinity` (t=-101.1; *p <* 2.2*e* − 16), but longer *Skeletonema* contigs than both `MEGAHIT` (t=41.6; *p <* 2.2*e* − 16) and `SPAdes` (t=64.0; *p <* 2.2*e* − 16).

## Discussion

Metatranscriptome analysis has become a widespread approach in the literature for extracting taxonomic and functional information from diverse environmental protistan communities (***Becker et al., 2021***; ***Salazar et al., 2019***; ***Stewart et al., 2012***; ***Gifford et al., 2011***; ***Alexander et al., 2015***; ***Vorobev et al., 2020***). Here, we designed a multi-assembler pipeline for metatranscriptomic assembly, and have evaluated the performance of this pipeline for environmental metatranscriptomes by simulating community data using reference transcriptomes from the MMETSP (***Keeling et al., 2014***). In doing this, we explored the relative performance of commonly-used assemblers, and determined that a multi-assembler approach improves the outcomes of metatranscriptome assembly with regard to recapitulating proteins and their taxonomic and functional annotations. Further, we applied the multiassembler pipeline, *euk* `rhythmic`, to metatranscriptomes from two published environmental datasets (***Alexander et al., 2015***; ***Carradec et al., 2018***).

### Do metatranscriptomes capture the diversity of protistan communities?

Environmental metatranscriptomes are a community mosaic of ephemeral RNA-based signals of expression. Metatranscriptomes are increasingly a routine diagnostic tool for making important conclusions about community composition and function within marine systems (***Gilbert et al., 2010***; ***Vorobev et al., 2020***; ***Alexander et al., 2015***; ***Nowinski et al., 2019***; ***Becker et al., 2021***; ***Salazar et al., 2019***; ***Gifford et al., 2011***), and are being applied to establish comparison at global scales (***Vorobev et al., 2020***; ***Carradec et al., 2018***) and over long time periods (***Vislova et al., 2016***; ***Ollison et al., 2021***; ***Hu et al., 2018***). Despite this, best practices for physical collection, molecular processing, and bioinformatic analyses have yet to be established. Towards the standardization of computational approaches to marine protistan metatranscriptomics, we have shown that a multi-tiered eukaryotic metatranscriptome assembly pipeline recapitulates annotated contigs from a mock transcriptome community. Notably, the contigs produced from multiple assemblers tend to be of the highest quality with regard to their similarity to the original contigs from the transcriptome assemblies via clustering, taxonomic, and functional annotations.

Our reassembly of environmental metatranscriptomic datasets further highlights the power of the multi-assembler approach in recovering novel gene content. In the samples from diatom-dominated Narragansett Bay (***Alexander et al., 2015***), we recovered a greater diversity of diatoms than raw read mapping alone, a level of diversity which aligns with other studies from the region (***Canesi and Rynearson, 2016***). From the *Tara* Oceans samples, we found novel protein sequences not recovered and included in a comprehensive, global analysis effort using a single assembler (***Carradec et al., 2018***), more than half of which had functional and/or taxonomic annotations. Even when the final coding sequences were clustered and only contigs of sufficient length were retained following ***Carradec et al. (2018)***, all samples contained previously unknown coding sequences, and some samples contained more unknown than known sequences. Though not all of these coding sequences could be annotated, recent progress has been made towards annotating genes of unknown function (***Vanni et al., 2021***), which can be highly abundant in metatranscriptomic data. These results demonstrate the value in reassembling previously analyzed datasets using multiple tools with different underlying algorithms.

Identifying the most salient and appropriate metrics for the claim that a single organism has been accurately identified and its functions accurately described from a metatranscriptome poses a significant challenge for the field. This is particularly true for environmental community data in which taxonomic boundaries might not be fully resolved in the first place, and culture representatives may not be available. To further complicate matters, even when assembly products can be shown to be “accurate” relative to commonly used metrics such as contig length, percentage of the raw sequencing reads mapping back to the assembly, and the presence of annotated genes with homology to “core” reference genes, they are not guaranteed to offer the best solution to the assembly problem because of the lack of database representatives (***Bushmanova et al., 2019***). One important note from our analysis is that even the best resource that we have available for consensus-based taxonomic annotation of *de novo* mixed community metatranscriptome assemblies constrains our efforts before we begin: laboratory-derived sequenced transcriptomes of single organisms cannot be fully reverse-annotated. In other words, even when we use sequence search tools to recover the taxonomic annotation of a contig present in the database, some of these sequences are too short or share a non-negligible percentage of sequences between organisms and cannot be annotated, even in their unmodified state. In these cases, the fact that we can recover many, but not all, of originally annotated contigs after reverse-engineering the community tells us more about the limits of taxonomic annotation via sequences than it does about the pitfalls of the assembly process. Hence, it is critical that we continue to consider the shortcomings of the annotation process as we analyze and re-analyze metatranscriptomic datasets.

### Is there one best assembler for eukaryotic environmental metatranscriptomes?

An additional goal of our analysis was to compare the performance of different assemblers on eukaryotic metatranscriptome data, and to determine whether the use of multiple assemblers is warranted. According to our results, no single assembler we assessed (*MEGAHIT* (***Li et al., 2015***), `rnaSPADes` (***Bushmanova et al., 2019***), `metaSPAdes` (***Nurk et al., 2017***), and `Trinity` (***Haas et al., 2013***)) is universally the best choice. *De novo* sequence assembly has both technical and practical considerations. Beyond simply balancing run time, memory requirements, and accuracy, assembler performance metrics is are not obvious. In particular in community metatranscriptome assembly, low sequencing depth, potentially unevenly across community members, may complicate typical algorithmic approaches used to reduce the impact of sequencing error. In our study, two assemblers - `MEGAHIT` (***Li et al., 2015***) and `rnaSPAdes` (***Bushmanova et al., 2019***) - stood out as the bookends of the spectrum of assembly approaches. `MEGAHIT` produced longer contigs, but had the proportion of raw reads recruited to the assembly, while `rnaSPAdes` routinely had the highest raw read mapping percentage and number of functional annotations (Figure 8; Supplementary Figure 8.1), but had shorter contigs on average, and a high incidence of transcripts that did not appear to be coding. These patterns of performance held in both simulated and environmental datasets (Figures 4 and 6).

The spectrum of approaches adopted by individual assemblers have a significant effect on the interpretation of the metatranscriptome assembly. The average length of contigs recovered by an assembler (e.g. `MEGAHIT` and `rnaSPAdes`) can directly skew interpretation of community composition. Shorter, potentially fragmented transcripts will recruit fewer reads, but will appear more abundant when a normalization that takes sequence length into account is used (***Wagner et al., 2012***). Because assemblers that work like `rnaSPAdes` produce a greater number of shorter contigs that may not be annotated, organisms or individual contigs or predicted genes with longer transcript length will appear comparatively less abundant when reads are normalized (Supplementary Figure 8.3-5). For example, in the Narragansett Bay samples, we observe that contigs annotated as diatom *Skeletonema* appears to have higher mean length. However, conventional community composition metrics like TPM that normalize to contig length will penalize the recruitment of raw reads to these longer-than-mean contigs. However, as has been well described for transcriptomics, using raw reads leaves interpretation vulnerable to biases related to sequencing depth, sequencing approach, and transcript length, intuitively because longer transcripts are expected to recruit a greater number of raw reads by virtue of their size (***Wagner et al., 2012***). In a mixed community sample, normalizations need to take the heterogeneity of the community into account.

Taken together, these results support the potential utility of merging the subtly different approaches taken by different assembly tools, in order to maximize gene recovery whilst also retaining the distinct signatures that make community composition interpretable. `rnaSPAdes`-like approaches improve functional recovery, while `MEGAHIT`-like assemblers produce longer sequences, which possibly have higher fidelity to the observed community. This observation further raises the question of how we can or should be extracting community composition insights from metatranscriptomes, especially when samples cannot or have not been normalized to housekeeping or spike-in sequences.

### Should we be downsizing our metatranscriptomic assemblies?

Computational constraints continue to limit the scale of metatranscriptomic analyses, as downstream tools for e.g. abundance quantification and functional annotation may have sizable memory requirements for excessively large assembly files (***Shakya et al., 2019***). Here, we advocate for a multi-assembler approach to metatranscriptome assembly. As we have discussed, the multi-assembler approach generates a greater number of total predicted coding sequences, and many of the additional coding sequences assembled from our simulated dataset are similar taxonomically, functionally, and via sequence identity to coding sequences from the designer assembly (Figures 5, 6, 7). However, using a multi-assembler approach will create larger assemblies, and users need to be cognizant of the complexity of their dataset and memory usage requirements downstream. The size of the assembly may be addressed by: (1) intentionally limiting the assembly to downsized, high-quality content (i.e., content recovered by all assemblers), or (2) by more strictly clustering assembly products.

Broadly, the contigs that were recovered by all assemblers were most likely to contain detectable open reading frames and to closely resemble “true” sequence content in a real-world sample via `blast` sequence search and `mmseqs2` clustering (Figure 7). Researchers may choose to maximize confidence in assembly products by using only the contigs discovered by more than one assembler. While the intention of *euk* `rhythmic` is to combine the outputs of multiple independently-contributing tools, in an analysis in which the goal is to extract only the products which can be assumed to be of the highest quality, the smaller intersection between assembly tools may be retained. This would also substantially reduce the number of sequences, improving the computational feasibility of downstream analyses. For example, if a researcher was interested specifically in generating a core set of high-confidence genes for a site and then mapping raw reads back to a combined assembly to detect changes in expression over time and space, multiple assembly may furnish a set of transcripts more likely to both be accurately retrieved from the original samples as well as ecologically relevant for mapping. However, it is important to note two pitfalls of this approach. First, this decreases the proportion of the raw reads that are represented in the final contigs after assembly (Figure 7). Secondly, while some assemblers produce a higher number of sequence products that *do not* have detectable similarity to the “true” contigs from which raw reads were simulated, they also produce a number of *unique* sequences that *are* detectable in the original assembly and, crucially, not identified by any of the other assembly approaches (e.g., `rnaspades`, Figure 7).

Researchers may also opt to cluster the resulting assembly in accordance with their desired final file size or level of sequence redundancy. Using `mmseqs2` clustering (***Mirdita et al., 2019***), we found that for our combined assemblies, the choice of clustering parameters is an important one, with potentially significant reductions in file size being possible without appreciable impact on the functional and taxonomic profile of the assembled metatranscriptome (Figure 3; Supplementary Figure 1). Approaches are now being developed to more reliably and efficiently cluster predicted genes of both known and unknown function, for example using tools like `mmseqs2` coupled to functional domain information and probabilistic modeling (***Vanni et al., 2021***). Such approaches are particularly useful for fitting coding sequences from a single assembly into the context of expansive datasets in space and/or time, from which many millions of total coding sequences can be extracted, and computational processing becomes exceptionally limiting (***Vanni et al., 2021***).

### *euk*rhythmic: an approach for optimized multi-assembler metatranscriptome assembly

Metatranscriptome quality cannot be assessed using either genomic or single-organism metrics. Instead, the assembly products need to be considered as potentially novel genetic content when assessing assembly success. Here, we presented *euk* `rhythmic`, a workflow for assembling environmental metatranscriptomes of eukaryotic communities by leveraging multiple assemblers. We evaluate our pipeline both using existing environmental metatranscriptomes and simulated community data that we generated using a second pipeline, `jEUKebox`. The flexible `jEUKebox` pipeline can be reused as additional reference sequences become available in order to test community ecology hypotheses for cultured and uncultured organisms. Simulating communities and testing their ability to be recovered is an essential step in ensuring the fidelity of metatranscriptome studies as the volume of taxonomic and functional data available to make predictions grows. In particular, we envision the construction of metacommunity data using uncultured organisms inferred from metagenomic sequences (metagenome-assembled genomes (MAGs); ***Alexander et al. (2021)***). Our inability to annotate some unmodified contigs from the original simulated community highlights crucial questions about the limits of annotation. Are some genes destined to remain difficult to annotate, either because they vary too dramatically between organisms, hence an organism-specific and highly complete genome is needed to accurately identify them, or because they are part of an indistinguishable cluster of highly similar genes? Can we be sure that these are true genes, or could they be artifacts of the assembler originally used to generate the reference assemblies? Rigorous simulations of communities may help to identify these difficult-to-annotate genes and to set thresholds that prevent erroneous annotations, in conjunction with new approaches for annotating unknown genes (***Vanni et al., 2022***). Computational simulations should be paired with laboratory curation of cultivated communities and accompanying metatranscriptomic sequencing which can be compared with count data. Promising plans for executing these steps are already in place (***Berube, 2022***).

Critical assessment of the accuracy and quality of metatranscriptome assembly and quantification of technical impacts, such as clustering similarity or the algorithms used to construct contigs, provide confidence for ecological interpretations. The *euk* `rhythmic` pipeline represents a reproducible roadmap towards assembling new eukaryotic environmental metatranscriptomes, and reassembling the growing repository of existing eukaryotic environmental metatranscriptomes with multiple assemblers. This flexible tool that researchers can use to standardize the crucial steps of metatranscriptome analysis is a step towards the standardization and validation eukaryotic metatranscriptome assembly. With the consistent use of software tools and pre- and post-processing steps that *euk* `rhythmic` enables, metatranscriptome assembly has the potential to unlock the functional roles of largely uncharacterized eukaryotic microbes that drive biogeochemistry across diverse natural ecosystems. Standardized workflows for eukaryotic metatranscriptome assembly like *euk* `rhythmic` and community simulations as powered by tools like `jEUKebox` are a vital means to validate these discoveries.

## Materials and Methods

### Eukrhythmic pipeline

#### Data cleaning and trimming

Trimming is performed using `Trimmomatic`, a flexible tool that is specifically suited to paired-end next-generation sequencing data, with user-specifiable parameters (***Bolger et al., 2014***). Optionally, the user may also choose to filter out spike-in sequences, if they were added during extraction, with `bbmap` (***Bushnell, 2014***).

#### Assembly

One major advantage of using the *euk* `rhythmic` pipeline is the flexibility to use as many (or as few) transcriptome assemblers as is appropriate for the data. Many different metatranscriptome assemblers are available to researchers and commonly used, and it can be challenging to select the appropriate assembler, given that each often has its own advantages and disadvantages (***Honaas et al., 2016***; ***Clarke et al., 2013***). Among the assemblers considered are MetaVelvet (***Namiki et al., 2012***), the String Graph Assembler (SGA) (***Simpson and Durbin, 2012***), Trinity (***Grabherr et al., 2011***), and MEGAHIT ***Li et al. (2015)***. In *euk* `rhythmic`, the user may select any combination of assemblers, and the assembly process is conducted in parallel, as resources allow.

#### Merging and Clustering

The consolidation of outputs from the constituent metatranscriptome assemblers is performed in two steps. First, assemblies from the same sample or user-defined “assembly group” (considered a single unit due to some shared characteristic) are concatenated. Inspired by the process adopted by Cerveau et al. (2016) (***Cerveau and Jackson, 2016***), we used the `MMSeqs` clustering tool (***Mirdita et al., 2019***) to eliminate similar contigs from the combined assembly, first using a similarity threshold of 100% for the shorter sequence in a local alignment to remove identical contigs recovered by multiple assemblers. Next, the pipeline branches into two output types. For the first output type, individual samples/assembly groups (“CAG”, or “clustered by assembly group”), which then undergo a second round of `MMSeqs` clustering to remove similar contigs at a 98% similarity threshold (defined the same way as above), accounting for potential sequencing errors (***Cerveau and Jackson, 2016***). Additionally, samples already merged from the assembly process are then merged between samples, such that one combined assembly is produced with all available data, labeled “multiple assembly consolidation” or abbreviated to “MAD” (“multi-assembler deduplicated assemblies”) in the text. We then cluster the combined assembly at the 98% level of similarity using `MMSeqs2` as previously described.

#### Protein Translation

To accommodate protein-space downstream analysis, such as protein families database (Pfam) annotation (***El-Gebali et al., 2019***), protein translation with TransDecoder (***Haas and Papanicolaou, 2021***) is supported as part of *euk* `rhythmic`. Both the output individual sample/assembly group files from the two clustering steps and the single combined assembly are translated to protein sequences.

#### Annotation

While *euk* `rhythmic` is primarily designed for assembly, the user may optionally elect to annotate the assembly output as part of the pipeline. Presently, the pipeline provides annotation tools including phylogenetic assessment using `EUKulele` (***Krinos et al., 2021***), and basic taxonomic assessment using established gene and protein annotation databases and/or companion tools.

Briefly, using the Hidden Markov Model-powered `hmmsearch` function from the `HMMSEARCH` package, the Pfam database is used to assign contigs to likely protein families (***Johnson et al., 2010***; ***El-Gebali et al., 2019***). In addition, the contigs are assigned a likely match to the KEGG Database (***Kanehisa et al., 2002***) using `DIAMOND` (***Buchfink et al., 2015***) and a simple set of scripts, `arKEGGio`, which we adapted to manage parsing of the database and its modules and pathways (***Krinos and Alexander, 2020***). To characterize KEGG annotations, we grouped results by Kegg Orthology ID (KO) and only considered the hit with the highest percentage identity to the query contig. When multiple relevant pathways were associated with a single hit, we assigned counts evenly to the assigned pathways.

### Design of simulated community schema

#### Communities

The six simulated communities were designed to have differing complexity and to represent community ecotypes that might be encountered in real-world metatranscriptomic studies. These configurations are summarized visually in Figure 2 and in terms of their complexity in Table 3. Community 1 was designed to resemble a community dominated by a single organism, thus has the lowest Shannon diversity index and species richness (see calculations in Section). Community 2 has a similar species richness value and only marginally higher diversity, since two strains of the same species make up the majority of the sample. Community 3 has the highest number of genes which are not shared between any of the organisms in the sample, but lower diversity than communities 4 and 6, which have the highest total species diversity. Community 4 has more genes shared between two closely-related groups. Community 5 has the highest total number of reasonably related organisms and shared genes. For MMETSP group B, the list of MMETSP IDs to choose from was selected randomly, and individual community pairings were determined using `fastANI` similarity (see Section).

#### Metrics for assessing community complexity

The Shannon diversity index of each community was calculated using the following formula ***Shannon (1948)***:

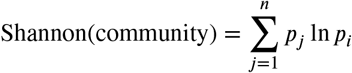

where *n* is the total number of “types” of community members, and *p* is their proportion in their community. Total species richness was reported as the total number of types present in the community.

We used `sourmash` to compute the pairwise similarity of each MMETSP transcriptome within each community (***Brown and Irber, 2016***). We additionally introduce another diversity metric to account for the potential similarity of the transcriptomes beyond their taxonomic annotations:

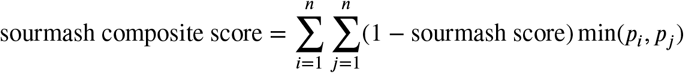

In other words, for each pair of transcriptomes in the community, we weight the `sourmash` similarity score of the pair of transcriptomes by the abundance of the less-abundant transcriptome in the pair. We report the sum of these weighted scores for each community in Table 3.

### Simulation of eukaryotic communities using jEUKebox

#### Selection of transcriptomes

For each set of simulated eukaryotic communities, 12 transcriptomes from the MMETSP ***Keeling et al. (2014)***; ***Caron et al. (2017)*** were used. These are summarized in Table 4 for the two selected communities. For “community A”, the IDs, but not the contigs selected, were included based on their features, including some MMETSP IDs of the same species and some of closely related strains. For “community B”, the MMETSP transcriptomes were selected randomly by `jEUKebox`, with the only constraint being the inclusion of some closely-related taxa.

#### Similarity computation using fastANI

In order to select “closely-related” transcriptomes for the community specifications, we used `fastANI` (***Jain et al., 2018***) to calculate the average nucleotide-based sequence identity between transcriptomes and identify similar transcriptomes on the basis of having ≥ 80% average nucleotide identity. Hence, for e.g. community 2 (see Figure 2), two MMETSP transcriptomes with `fastANI` similarity 2: 80% would be selected.

#### Identifying putative evolutionary relationships with OrthoFinder

In order to test the skill of metatranscriptome assembly with respect to recovering genes of shared evolutionary origin, but different current annotated taxonomic identity, we used the tool `OrthoFinder` to identify orthologous groups between the MMETSP transcriptomes, and to include genes from both highly-conserved and relatively rare gene groups in the designer metatranscriptome (***Emms and Kelly, 2019, 2015***). As summarized in Figure 2, the jEUKebox pipeline automates this process by including 10% of all of the identified single-copy orthologs reported by `OrthoFinder` (orthologous groups with a single gene representative from every transcriptome in the community). Genes are then selected for each one of the organisms in the community according to the following formula. For genes that *do* have a “highly related” partner with respect to computed similarity (see Section “Similarity computation using `fastANI`”), 75% of the contigs to be included in the designer transcriptome (as prescribed by the desired ratio of the candidate organism in the final metatranscriptomes) were taken from orthologous groups which included more than just the candidate. The remaining 25% were randomly selected from orthologous groups that only contained the candidate. For genes *without* a highly-related partner, 75% of genes were taken from exclusive orthologous groups containing only the candidate. The remaining 25% were randomly selected from orthologous groups shared with other MMETSP transcriptomes.

#### Simulation of raw reads

After creating the designer metatranscriptomes directly from informed random selection of contigs from the MMETSP transcriptomes, raw reads were simulated using the package `Rsubread` (***Liao et al., 2019***). We chose a read length of 75 base pairs to enable the `simReads` function to use its inbuilt set of quality scores to randomly determine a sequencing error for the generated raw reads. We chose a mean fragment length of 180±40 base pairs and generated a 1 million base pair library for the paired sequencing reads which were simulated using the package for each community and trial.

#### Reassembly with *euk* rhythmic

The raw reads simulated using the `simReads` function were provided as input to the eukrhythmic pipeline. The pipeline was run with default settings as described in Section “Eukrhythmic pipeline”. Four assemblers were used: `rnaSPAdes` (***Bushmanova et al., 2019***), `MEGAHIT` (***Li et al., 2015***), `metaSPAdes` (***Nurk et al., 2017***), and `Trinity` (***Haas et al., 2013***).

### Assessing reassembly quality

#### Assembly statistics

We used the salmon mapping tool to both quantify the abundance of each contig with respect to the raw reads, and to assess what proportion of the raw reads were represented in the assembled contigs.

We report descriptive statistics for the contigs assembled as a proxy for the quality of the assembled sequences. These include minimum, maximum, mean, and standard deviation of contig length, as well as the N50 metric. We use the definition of the N50 metric as the minimum length among the set of contigs which together constitute 50% of the total length of all contigs in the assembly, as reported by QUAST (***Bushmanova et al., 2016***).

#### Clustering reassembled metatranscriptomic proteins with MMETSP-derived designer metatranscriptomic proteins

To determine whether exact sequence matches were shared between the predicted proteins from the metatranscriptome assembly and the proteins from the MMETSP used to create the designer metatranscriptome, we performed `mmseqs2` clustering between the two protein sets (***Mirdita et al., 2019***). We chose the LINCLUST algorithm as implemented in `mmseqs2` due to its exceeding low false discovery rate in clustering (***Steinegger and Söding, 2017, 2018***). In accordance with what was used by authors of `mmseqs2`, we report these results using minimum coverage of the target sequence (`–cov-mode 1`) of 90% and a minimum sequence identity of 90%, at which threshold fewer clusters are produced but there is very little chance of a false negative, i.e. two 90%-similar sequences in the dataset which `mmseqs2` fails to report.

When evaluating the likelihood of contigs assembled using eukrhythmic to cluster with the designer contigs, we based the comparison on the protein predictions from `TransDecoder` (***Haas and Papanicolaou, 2021***) as clustered through `mmseqs2`. For each full nucleotide contig, we considered it to have “clustered with the designer metatranscriptome” if at least one ORF from `TransDecoder` was successfully co-clustered with a protein from the designer assembly, even though transcriptome assembly happens in nucleotide space. This enabled us to also quantify what proportion of the contigs from the eukrhythmic assembly were not assigned an ORF at all by the `TransDecoder` software.

#### Assessing metatranscriptomic proteins using BLAST all-by-all comparison

In addition to clustering, we performed an all-by-all BLAST search between the proteins from the original contigs from the MMETSP and the resulting predicted proteins from eukrhythmic. An e-value cutoff of 10^−2^ was used to catch the top match on the basis of bitscore, and then hits were classified according to their percentage identity and bitscore value.

#### Taxonomic annotations

As performed within the eukrhythmic pipeline, we generated taxonomic annotations for both the designer metatranscriptomes and the reassembled products from eukrhythmic with the `EUKulele` tool using the default reference database of contigs from all MMETSP transcriptomes and the MarRef database (***Krinos et al., 2021***; ***Keeling et al., 2014***; ***Caron et al., 2017***; ***Klemetsen et al., 2018***). We report differences in the number of annotated species and genera from `EUKulele` in the reassembled products as compared to the sequences which were prescribed to be included in the designer metatranscriptome using the jEUKebox pipeline. We also compare the `EUKulele` annotations from the designer metatranscriptomes, including false matches on the basis of poor-quality sequences being present in the database and failing to be annotated to begin with, to the annotations of the reassembled products. We perform standard linear regression on the number of annotations for each species, genus, order, and phylum from the designer metatranscriptomes as compared to the reassembled products. We also categorize taxonomic annotations according to whether they were classified correctly, incorrectly (in conflict with the original annotations), or were not classified. We performed a Welch’s 2-sample T-test for independent samples as implemented in `scipy` (***Virtanen et al., 2020***) to compare the summed abundances of correctly and incorrectly classified and unclassified sequences.

#### Functional annotations

All functional annotations were determined using `eggNOG-mapper` (***Huerta-Cepas et al., 2017***). Similarly to taxonomic annotations, reported annotations of orthology terms (KOs) from the Kyoto Encyclopedia of Genes and Genomes (KEGG) were compared between the designer metatranscriptomes via annotation of the contigs from the MMETSP and the reassembled products which were retrieved as an output of the eukrhythmic pipeline.

Standard linear regression was performed to compare the abundance of KEGG orthology terms in the designer metatranscriptomes as compared to the reassembled products from eukrhythmic. The regression and associated probability value was calculated using the implementation in base R (***R Core Team, 2021***).

### Assembling and evaluating environmentally relevant metatranscriptomes from the *Tara* Oceans project

We assembled metatranscriptomes from the *Tara* Oceans project (***Vorobev et al., 2020***; ***Sunagawa et al., 2015***) as an environmental counterpart to the simulated sequence data. Metatranscriptome samples from three distinct ocean basins were assembled from the highly-diverse small-size fraction surface samples from the *Tara* project: the North Atlantic, Southern Ocean, and Mediterranean Sea. We assembled these metatranscriptomes using default parameters to the eukrhythmic pipeline and used `MEGAHIT` and `rnaSPAdes`, which proved to be the fastest and most accurate assemblers, respectively, in both the present work and other investigations (***Ortiz et al., 2021***; ***Li et al., 2015***; ***Bushmanova et al., 2019***), and `Trans-ABySS`, another high-performing assembler for RNA sequence data (***Ortiz et al., 2021***). Three assemblers were selected so as to compare the mutual findings of the three assemblers to the unique sequence content identified by each. For the samples from the Mediterranean Sea, we also demonstrate *euk* `rhythmic`’s performance on a large co-assembly of samples from the same region, depth, and size fraction as compared to individual assembly of each sample. We assess the results of the metatranscriptome assembly via percentage mapping via `Salmon` ***Patro et al. (2017)***, and taxonomic and functional annotations as provided by `EUKulele` and `eggNOG-mapper`, respectively (***Krinos et al., 2021***; ***Huerta-Cepas et al., 2017***).

These metatranscriptomes were previously analyzed by ***Carradec et al. (2018)*** with transcribed sequences of length ≥150 bases assembled using `velvet` (***Zerbino and Birney, 2008***) included as part of the “MATOU” database (***Carradec et al., 2018***). In order to compare the contigs generated and retained from our multiassembler approach, we conducted a `blastn` (***Altschul et al., 1990***) search with e-value cutoff of 1e-10 to find the top-scoring match of “MATOU” transcribed sequences against our sequences, and compared the contigs that were successfully matched to the database using this method to those that otherwise could be functionally and/or taxonomically annotated. Identified coding sequences of length *>* 150 bases were retained for further analysis following ***Carradec et al. (2018)***.

### Reassembling and evaluating previously explored metatranscriptomes from the Narragansett Bay time series

We assembled ten samples from a 2015 metatranscriptomic study from the Narragansett Bay time series (***Alexander et al., 2015***). These samples are stored under National Center for Biotechnology Information (NCBI) project accession number SRP055134 and samples were assigned individual accession numbers collated in Supplementary Table 5. We assembled these metatranscriptomes using default parameters to the eukrhythmic pipeline and used `MEGAHIT`, `rnaSPAdes`, `metaSPAdes`, and `Trinity` (***Li et al., 2015***; ***Nurk et al., 2017***; ***Bushmanova et al., 2019***; ***Haas et al., 2013***). We compared the taxonomic and functional annotations between assemblers to the composition of major taxonomic groups reported by the 2015 study, which used raw read mapping to reference transcriptome assemblies rather than assembling the metatranscriptome itself (***Alexander et al., 2015***). We also compare the insights drawn from the simulated metatranscriptomes via `jEUKebox` to the patterns that emerge from using multiple assemblers on a previously analyzed environmental dataset.

### Data processing and visualization

Output data from the described tools were processed using Python version 3.8.3 (***Van Rossum and Drake Jr, 1995***) and R version 4.1.0 (***R Core Team, 2021***). Figures were generated using `plotnine` in Python ***Kibirige et al. (2021)*** or `ggplot2` (***Wickham, 2016***) in R with organization into panels using (***Pedersen, 2020***). Statistical analysis on the data was performed with `SciPy` (***Virtanen et al., 2020***) or with R version 4.1.0 (***R Core Team, 2021***).

**Table 4.** MMETSP members of each simulated community with the number of orthologous groups which include each organism and the assessed BUSCO completeness of each transcriptome. BUSCO completeness is a metric for the quality of the transcriptome based on the presence of shared ancestral eukaryotic genes (of a total of 255 evaluated genes). Reported orthologous groups are based on `OrthoFinder` ***Emms and Kelly (2015)*** analysis on all community members for each MMETSP group; the total number of reported orthogroups was 42,093 for MMETSP group A and 44178 for MMETSP group B.

| MMETSP Group | MMETSP ID | Organism | BUSCO Completeness | Number of Orthologous Groups |
| --- | --- | --- | --- | --- |
| A | MMETSP0027 | *Skeletonema marinoi* | 146/255 | 17057 |
| A | MMETSP0147 | *Chrysochromulina polylepis* | 119/255 | 8185 |
| A | MMETSP0448 | *Heterocapsa triquetra* | 124/255 | 14377 |
| A | MMETSP0562 | *Skeletonema dohrnii* | 134/255 | 10279 |
| A | MMETSP0604 | *Skeletonema menzelii* | 161/255 | 9599 |
| A | MMETSP0918 | *Skeletonema marinoi* | 135/255 | 9871 |
| A | MMETSP0971 | *Thalassiosira oceanica* | 152/255 | 11115 |
| A | MMETSP0994 | *Emiliana huxleyi* | 113/255 | 12435 |
| A | MMETSP0995 | *Emiliana huxleyi* | 99/255 | 12415 |
| A | MMETSP1403 | *Micromonas pusilla* | 137/255 | 3779 |
| A | MMETSP1405 | *Thalassiosira weissflogii* | 151/255 | 8800 |
| A | MMETSP1428 | *Skeletonema marinoi* | 152/255 | 10152 |
| B | MMETSP0321 | *Leptocylindrus danicus* | 169/255 | 8235 |
| B | MMETSP0369 | *Scrippsiella hangoei-like* | 155/255 | 20932 |
| B | MMETSP0397 | *Cyclophora tenuis* | 83/255 | 7306 |
| B | MMETSP0469 | *Oxyrrhis marina* | 160/255 | 9456 |
| B | MMETSP0800 | *Striatella unipunctata* | 100/255 | 8288 |
| B | MMETSP0884 | *Pelagococcus subviridis* | 112/255 | 7134 |
| B | MMETSP0896 | *Heterosigma akashiwo* | 62/255 | 3988 |
| B | MMETSP0975 | *Pelagomonadales sp.* | 171/255 | 9100 |
| B | MMETSP1117 | *Symbiodinium sp.* | 129/255 | 14718 |
| B | MMETSP1338 | *Pelagodinium sp.* | 141/255 | 17307 |
| B | MMETSP1349 | *Aplanochytrium stocchinoi* | 75/255 | 3747 |
| B | MMETSP1411 | *Thalassiosira weissflogii* | 144/255 | 6939 |

**Table 5.** Sample IDs and accession numbers for Narragansett Bay samples. Descriptive information about the sample conditions are reproduced from ***Alexander et al. (2015)***.

| Sample ID | Assembly Group | Accession Number | Experimental Context |
| --- | --- | --- | --- |
| NarBay_A | NarBay_A | SRR1810207 | Nitrate added (+N) |
| NarBay_B | NarBay_B | SRR1810208 | -N |
| NarBay_C | NarBay_C | SRR1810209 | Phosphate added (+P) |
| NarBay_D | NarBay_D | SRR1810210 | -P |
| NarBay_E | NarBay_E | SRR1810211 | No amendment |
| NarBay_S1 | NarBay_S1 | SRR1810799 | Environmental sample 1 |
| NarBay_S2 | NarBay_S2 | SRR1810204 | Environmental sample 2 |
| NarBay_S3 | NarBay_S3 | SRR1810801 | Environmental sample 3 |
| NarBay_S4 | NarBay_S4 | SRR1810205 | Environmental sample 4 |
| NarBay_S5 | NarBay_S5 | SRR1810206 | Environmental sample 5 |

**Table 6.** Sample ID and accession numbers for the 15 *Tara* Oceans metatranscriptomes assembled as part of this project, including the ocean basin they were sampled from. All analyzed samples were collected from surface water.

| Accession Number | Sample ID / Assembly Group | Ocean Basin |
| --- | --- | --- |
| ERR1712028 | SO_SRF_SMALL_ERR1712028 | Southern Ocean |
| ERR1719157 | SO_SRF_SMALL_ERR1719157 | Southern Ocean |
| ERR1740115 | SO_SRF_SMALL_ERR1740115 | Southern Ocean |
| ERR1740130 | SO_SRF_SMALL_ERR1740130 | Southern Ocean |
| ERR1740133 | SO_SRF_SMALL_ERR1740133 | Southern Ocean |
| ERR1711918 | MS_SRF_SMALL_ERR1711918 | Mediterranean Sea |
| ERR1711995 | MS_SRF_SMALL_ERR1711995 | Mediterranean Sea |
| ERR1711998 | MS_SRF_SMALL_ERR1711998 | Mediterranean Sea |
| ERR1712006 | MS_SRF_SMALL_ERR1712006 | Mediterranean Sea |
| ERR1712022 | MS_SRF_SMALL_ERR1712022 | Mediterranean Sea |
| ERR1719164 | MS_SRF_SMALL_ERR1719164 | Mediterranean Sea |
| ERR1719224 | MS_SRF_SMALL_ERR1719224 | Mediterranean Sea |
| ERR550386 | MS_SRF_SMALL_ERR550386 | Mediterranean Sea |
| ERR550396 | MS_SRF_SMALL_ERR550396 | Mediterranean Sea |
| ERR550403 | MS_SRF_SMALL_ERR550403 | Mediterranean Sea |
| ERR550404 | MS_SRF_SMALL_ERR550404 | Mediterranean Sea |

## Acknowledgments

We gratefully acknowledge Celeste Nobrega for her work on this project as a Guest Student at Woods Hole Oceanographic Institution. The Poseidon High Performance Computing cluster at Woods Hole Oceanographic Institution was used to run all analyses.

---

**Table S1.** (Supplement) Effect of clustering designer assembly on assembly size and annotations. Clustering was performed on the original “designer metatranscriptome” set of contigs from the MMETSP references using the `mmseqs2` tool ***Mirdita et al. (2019)***. *euk* `rhythmic` uses a coverage level of 0.98 and sequence identity of 1 for `mmseqs2` clustering. See Figure 1 Supplement for a graphical summary of the influence of sequence identity and coverage on the size of the recovered assembly and its functional and taxonomic annotations.

| MMETSP Group | Cluster Seq. ID | Cluster Cover-age | No. Contigs | File Size (MB) | No. Genera | No. Species | No. KOs |
| --- | --- | --- | --- | --- | --- | --- | --- |
| A | 0.6 | 0.6 | 35515 ± 6240 | 21.2 ± 3.7 | 3.6 ± 0.8 | 4.4 ± 1.2 | 4803.5 ± 403.1 |
| A | 0.6 | 1.0 | 40793 ± 6312 | 24.1 ± 3.8 | 3.8 ± 0.8 | 4.4 ± 1.2 | 4838.9 ± 403.4 |
| A | 0.75 | 0.8 | 38426 ± 6734 | 22.6 ± 4.0 | 3.6 ± 0.8 | 4.4 ± 1.2 | 4831.1 ± 404.4 |
| A | 0.8 | 0.8 | 39422 ± 6824 | 23.2 ± 4.1 | 3.8 ± 0.8 | 4.4 ± 1.2 | 4835.2 ± 404.9 |
| A | 0.99 | 0.6 | 47174 ± 6616 | 27.5 ± 4.3 | 3.8 ± 0.8 | 4.4 ± 1.2 | 4860.3 ± 399.8 |
| A | 0.99 | 0.8 | 47282 ± 6604 | 27.6 ± 4.3 | 3.8 ± 0.8 | 4.4 ± 1.2 | 4861.3 ± 399.7 |
| A | 0.99 | 0.95 | 47541 ± 6543 | 27.7 ± 4.2 | 3.8 ± 0.8 | 4.4 ± 1.2 | 4862.6 ± 399.7 |
| A | 0.99 | 0.99 | 47937 ± 6470 | 27.8 ± 4.2 | 3.8 ± 0.8 | 4.4 ± 1.2 | 4863.0 ± 399.6 |
| A | 0.99 | 1.0 | 48152 ± 6414 | 28.0 ± 4.2 | 3.8 ± 0.8 | 4.4 ± 1.2 | 4863.4 ± 399.5 |
| A | 1.0 | 0.6 | 49595 ± 6100 | 28.9 ± 4.1 | 3.8 ± 0.8 | 4.4 ± 1.2 | 4865.6 ± 398.7 |
| A | 1.0 | 0.8 | 49672 ± 6094 | 28.9 ± 4.0 | 3.8 ± 0.8 | 4.4 ± 1.2 | 4865.8 ± 398.7 |
| A | 1.0 | 0.95 | 49807 ± 6069 | 29.0 ± 4.0 | 3.8 ± 0.8 | 4.4 ± 1.2 | 4866.4 ± 398.7 |
| A | 1.0 | 0.99 | 50036 ± 6037 | 29.1 ± 4.0 | 3.8 ± 0.8 | 4.4 ± 1.2 | 4866.5 ± 398.8 |
| A | 1.0 | 1.0 | 50164 ± 6009 | 29.1 ± 4.0 | 3.8 ± 0.8 | 4.4 ± 1.2 | 4866.8 ± 398.7 |

Effect of clustering designer assembly on assembly size and annotations. Clustering was performed on the original “designer metatranscriptome” set of contigs from the MMETSP references using the `mmseqs2` tool ***Mirdita et al. (2019)***. `eukrhythmic` uses a coverage level of 0.98 and sequence identity of 1 for `mmseqs2` clustering. See Figure 1 Supplement for a graphical summary of the influence of sequence identity and coverage on the size of the recovered assembly and its functional and taxonomic annotations.

| MMETSP Group | Cluster Seq. ID | Cluster Cover-age | No. Contigs | File Size (MB) | No. Genera | No. Species | No. KOs |
| --- | --- | --- | --- | --- | --- | --- | --- |
| B | 0.6 | 0.6 | 38425 ± 5911 | 24.0 ± 4.1 | 4.9 ± 1.4 | 4.9 ± 1.4 | 5038.8 ± 312.7 |
| B | 0.6 | 1.0 | 41618 ± 6334 | 25.8 ± 4.4 | 4.9 ± 1.4 | 4.9 ± 1.4 | 5083.9 ± 311.0 |
| B | 0.75 | 0.8 | 40848 ± 6271 | 25.2 ± 4.3 | 4.9 ± 1.4 | 4.9 ± 1.4 | 5080.0 ± 310.4 |
| B | 0.8 | 0.8 | 41550 ± 6349 | 25.6 ± 4.4 | 4.9 ± 1.4 | 4.9 ± 1.4 | 5086.6 ± 309.9 |
| B | 0.99 | 0.6 | 46150 ± 6852 | 28.3 ± 4.8 | 4.9 ± 1.4 | 4.9 ± 1.4 | 5116.3 ± 306.8 |
| B | 0.99 | 0.8 | 46202 ± 6858 | 28.3 ± 4.8 | 4.9 ± 1.4 | 4.9 ± 1.4 | 5116.4 ± 306.8 |
| B | 0.99 | 0.95 | 46276 ± 6865 | 28.4 ± 4.8 | 4.9 ± 1.4 | 4.9 ± 1.4 | 5116.6 ± 306.6 |
| B | 0.99 | 0.99 | 46369 ± 6876 | 28.4 ± 4.8 | 4.9 ± 1.4 | 4.9 ± 1.4 | 5117.0 ± 306.4 |
| B | 0.99 | 1.0 | 46391 ± 6879 | 28.4 ± 4.8 | 4.9 ± 1.4 | 4.9 ± 1.4 | 5117.2 ± 306.4 |
| B | 1.0 | 0.6 | 46580 ± 6875 | 28.6 ± 4.8 | 4.9 ± 1.4 | 4.9 ± 1.4 | 5118.2 ± 306.3 |
| B | 1.0 | 0.8 | 46616 ± 6879 | 28.6 ± 4.8 | 4.9 ± 1.4 | 4.9 ± 1.4 | 5118.3 ± 306.3 |
| B | 1.0 | 0.95 | 46660 ± 6882 | 28.6 ± 4.8 | 4.9 ± 1.4 | 4.9 ± 1.4 | 5118.5 ± 306.0 |
| B | 1.0 | 0.99 | 46722 ± 6889 | 28.6 ± 4.8 | 4.9 ± 1.4 | 4.9 ± 1.4 | 5118.9 ± 305.9 |
| B | 1.0 | 1.0 | 46753 ± 6890 | 28.6 ± 4.8 | 4.9 ± 1.4 | 4.9 ± 1.4 | 5118.9 ± 305.9 |

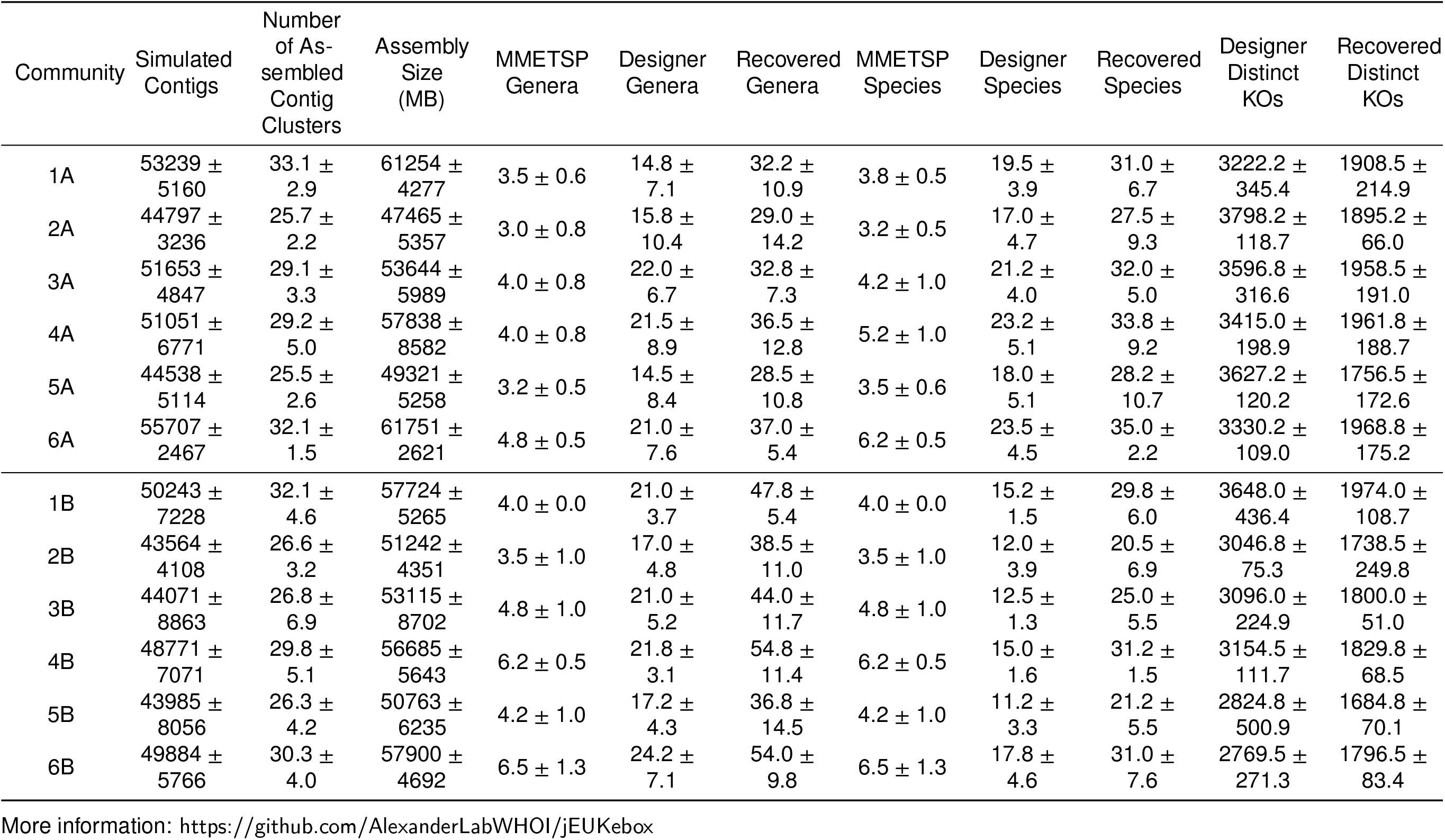
Resulting assembly size and taxonomic, functional, and core content recovery of `jEUKebox` outputs after raw read simulation and re-assembly with *euk* `rhythmic`. Four assemblers were used in this analysis which were then clustered together using default *euk* `rhythmic` settings. The mean and standard deviation of four trials of each community and list of MMETSP IDs is presented.

| Community | Simulated<br>Contigs | Number<br>of As-<br>sembled<br>Contig<br>Clusters | Assembly<br>Size<br>(MB) | MMETSP<br>Genera | Designer<br>Genera | Recovered<br>Genera | MMETSP<br>Species | Designer<br>Species | Recovered<br>Species | Designer<br>Distinct<br>KOs | Recovered<br>Distinct<br>KOs |
| --- | --- | --- | --- | --- | --- | --- | --- | --- | --- | --- | --- |
| 1A | 53239 ± | 33.1 ± | 61254 ± | 3.5 ± 0.6 | 14.8 ± | 32.2 ± | 3.8 ± 0.5 | 19.5 ± | 31.0 ± | 3222.2 ± | 1908.5 ± |
|  | 5160 | 2.9 | 4277 |  | 7.1 | 10.9 |  | 3.9 | 6.7 | 345.4 | 214.9 |
| 2A | 44797 ± | 25.7 ± | 47465 ± | 3.0 ± 0.8 | 15.8 ± | 29.0 ± | 3.2 ± 0.5 | 17.0 ± | 27.5 ± | 3798.2 ± | 1895.2 ± |
|  | 3236 | 2.2 | 5357 |  | 10.4 | 14.2 |  | 4.7 | 9.3 | 118.7 | 66.0 |
| 3A | 51653 ± | 29.1 ± | 53644 ± | 4.0 ± 0.8 | 22.0 ± | 32.8 ± | 4.2 ± 1.0 | 21.2 ± | 32.0 ± | 3596.8 ± | 1958.5 ± |
|  | 4847 | 3.3 | 5989 |  | 6.7 | 7.3 |  | 4.0 | 5.0 | 316.6 | 191.0 |
| 4A | 51051 ± | 29.2 ± | 57838 ± | 4.0 ± 0.8 | 21.5 ± | 36.5 ± | 5.2 ± 1.0 | 23.2 ± | 33.8 ± | 3415.0 ± | 1961.8 ± |
|  | 6771 | 5.0 | 8582 |  | 8.9 | 12.8 |  | 5.1 | 9.2 | 198.9 | 188.7 |
| 5A | 44538 ± | 25.5 ± | 49321 ± | 3.2 ± 0.5 | 14.5 ± | 28.5 ± | 3.5 ± 0.6 | 18.0 ± | 28.2 ± | 3627.2 ± | 1756.5 ± |
|  | 5114 | 2.6 | 5258 |  | 8.4 | 10.8 |  | 5.1 | 10.7 | 120.2 | 172.6 |
| 6A | 55707 ± | 32.1 ± | 61751 ± | 4.8 ± 0.5 | 21.0 ± | 37.0 ± | 6.2 ± 0.5 | 23.5 ± | 35.0 ± | 3330.2 ± | 1968.8 ± |
|  | 2467 | 1.5 | 2621 |  | 7.6 | 5.4 |  | 4.5 | 2.2 | 109.0 | 175.2 |
| 1B | 50243 ± | 32.1 ± | 57724 ± | 4.0 ± 0.0 | 21.0 ± | 47.8 ± | 4.0 ± 0.0 | 15.2 ± | 29.8 ± | 3648.0 ± | 1974.0 ± |
|  | 7228 | 4.6 | 5265 |  | 3.7 | 5.4 |  | 1.5 | 6.0 | 436.4 | 108.7 |
| 2B | 43564 ± | 26.6 ± | 51242 ± | 3.5 ± 1.0 | 17.0 ± | 38.5 ± | 3.5 ± 1.0 | 12.0 ± | 20.5 ± | 3046.8 ± | 1738.5 ± |
|  | 4108 | 3.2 | 4351 |  | 4.8 | 11.0 |  | 3.9 | 6.9 | 75.3 | 249.8 |
| 3B | 44071 ± | 26.8 ± | 53115 ± | 4.8 ± 1.0 | 21.0 ± | 44.0 ± | 4.8 ± 1.0 | 12.5 ± | 25.0 ± | 3096.0 ± | 1800.0 ± |
|  | 8863 | 6.9 | 8702 |  | 5.2 | 11.7 |  | 1.3 | 5.5 | 224.9 | 51.0 |
| 4B | 48771 ± | 29.8 ± | 56685 ± | 6.2 ± 0.5 | 21.8 ± | 54.8 ± | 6.2 ± 0.5 | 15.0 ± | 31.2 ± | 3154.5 ± | 1829.8 ± |
|  | 7071 | 5.1 | 5643 |  | 3.1 | 11.4 |  | 1.6 | 1.5 | 111.7 | 68.5 |
| 5B | 43985 ± | 26.3 ± | 50763 ± | 4.2 ± 1.0 | 17.2 ± | 36.8 ± | 4.2 ± 1.0 | 11.2 ± | 21.2 ± | 2824.8 ± | 1684.8 ± |
|  | 8056 | 4.2 | 6235 |  | 4.3 | 14.5 |  | 3.3 | 5.5 | 500.9 | 70.1 |
| 6B | 49884 ± | 30.3 ± | 57900 ± | 6.5 ± 1.3 | 24.2 ± | 54.0 ± | 6.5 ± 1.3 | 17.8 ± | 31.0 ± | 2769.5 ± | 1796.5 ± |
|  | 5766 | 4.0 | 4692 |  | 7.1 | 9.8 |  | 4.6 | 7.6 | 271.3 | 83.4 |

More information: https://github.com/AlexanderLabWHOI/jEUKebox

**Figure 1–Figure supplement 1.**
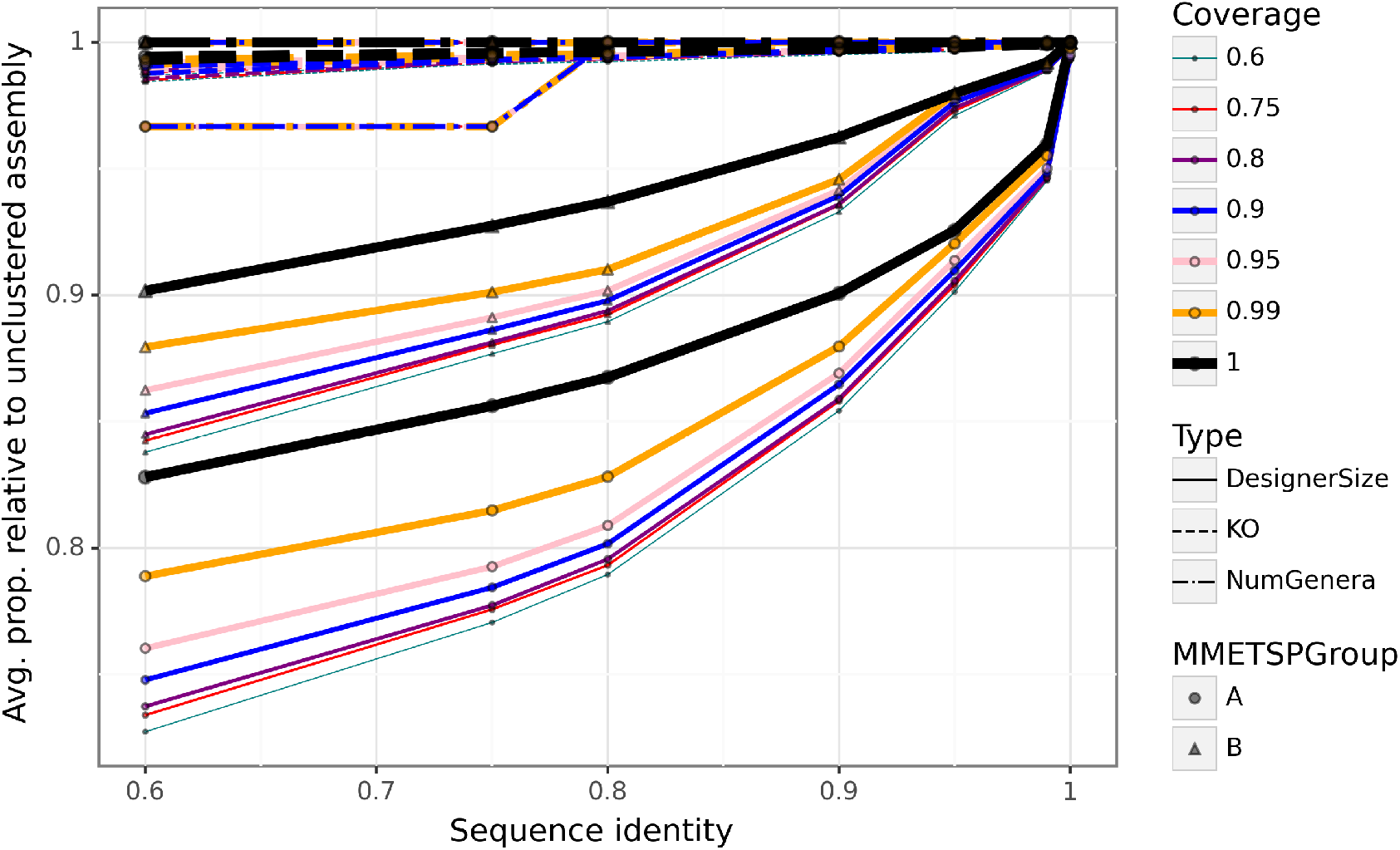
Visual summary of the effect of clustering with `mmseqs2` on recovered genera, KOs (functional annotations) and file size in bytes. While reducing the sequence identity threshold for clustering results in up to a 30% average reduction in file size, the number of recovered genera and functional annotations are only modestly impacted, especially at high coverage. An intermediate sequence identity of 0.8 and coverage of 0.8 would result in a 15-25% average reduction in file size, but leave distinct functional and taxonomic annotations unchanged.

**Figure 4–Figure supplement 1.**
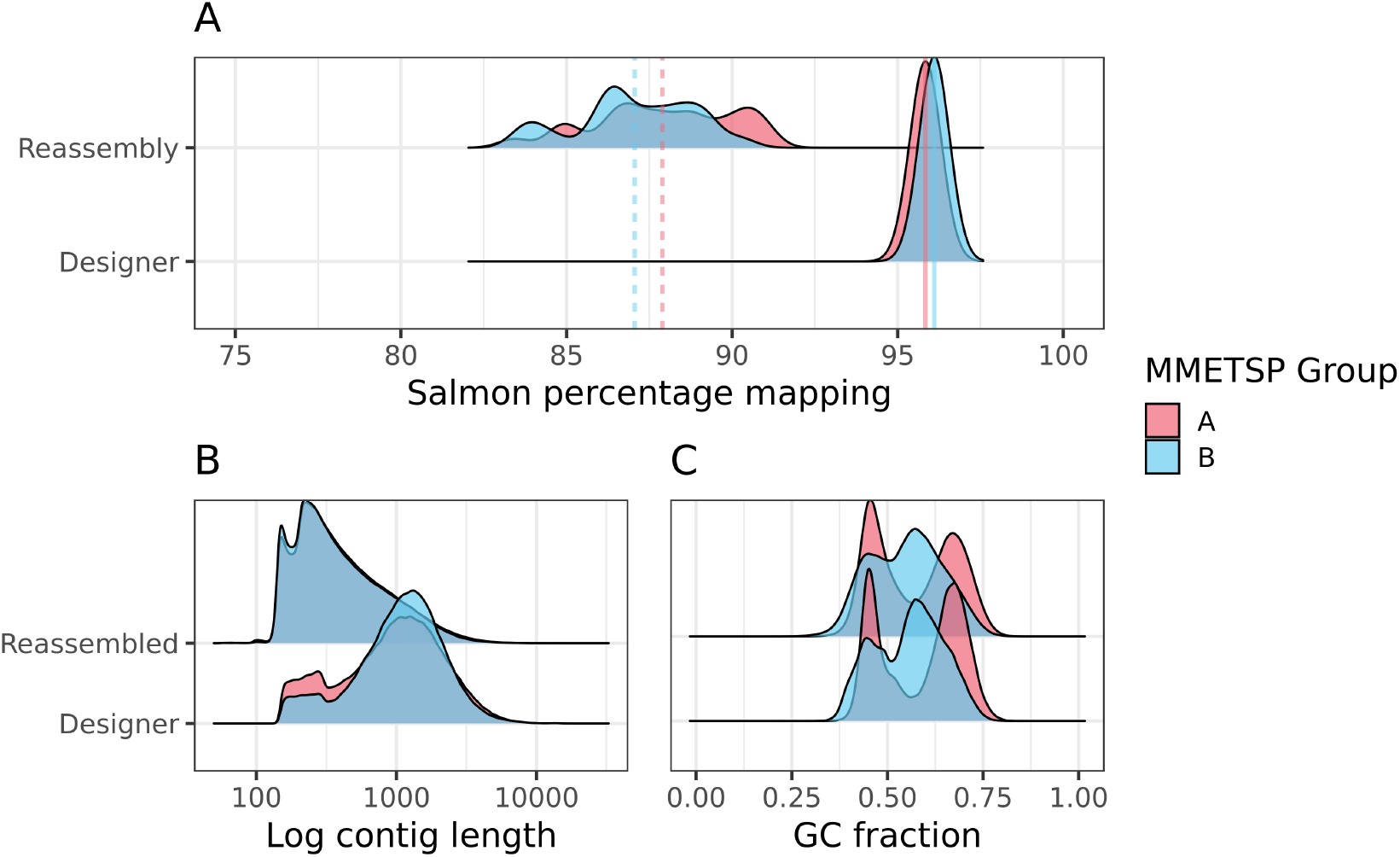
Main text figure facetted by simulated MMETSP assembly group (two different sets of organisms). Of note is that the eukrhythmic reassembly accurately recapitulates the bimodal distribution in GC-content observed in MMETSP group A’s designer metatranscriptomic sequences.

**Figure 4–Figure supplement 2.**
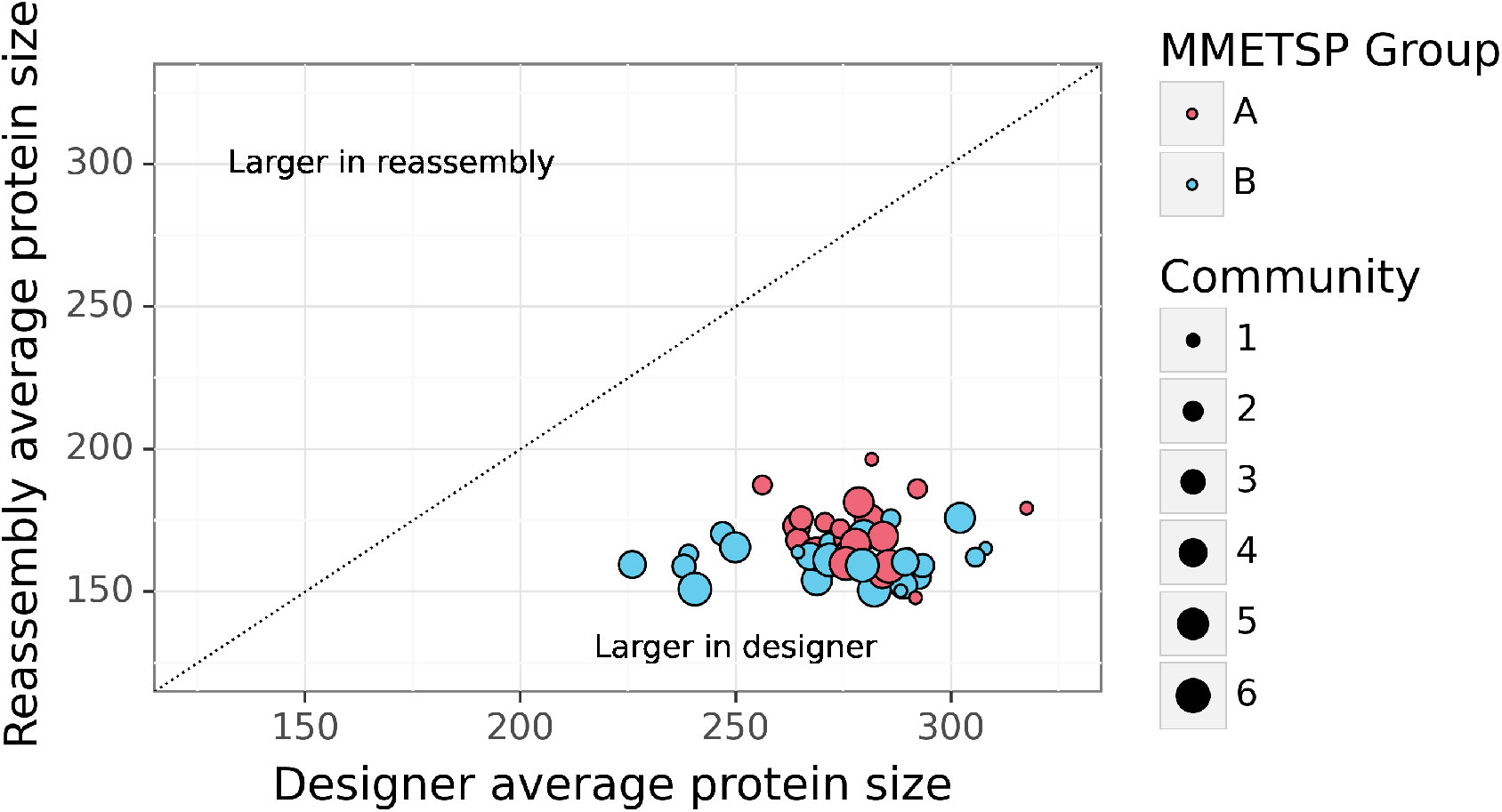
Protein sequence lengths in the reassemblies as compared to the designer. The 1-to-1 line shows where sequences would fall if the average length of recovered protein sequences via `TransDecoder` were identical between the designer assemblies and the eukrhythmic-derived reassembled products; the fact that all samples fall in the lower right half of the plot indicates that protein sequences were consistently larger in the designer assemblies as compared to the eukrhythmic reassembled products.

**Figure 4–Figure supplement 3.**
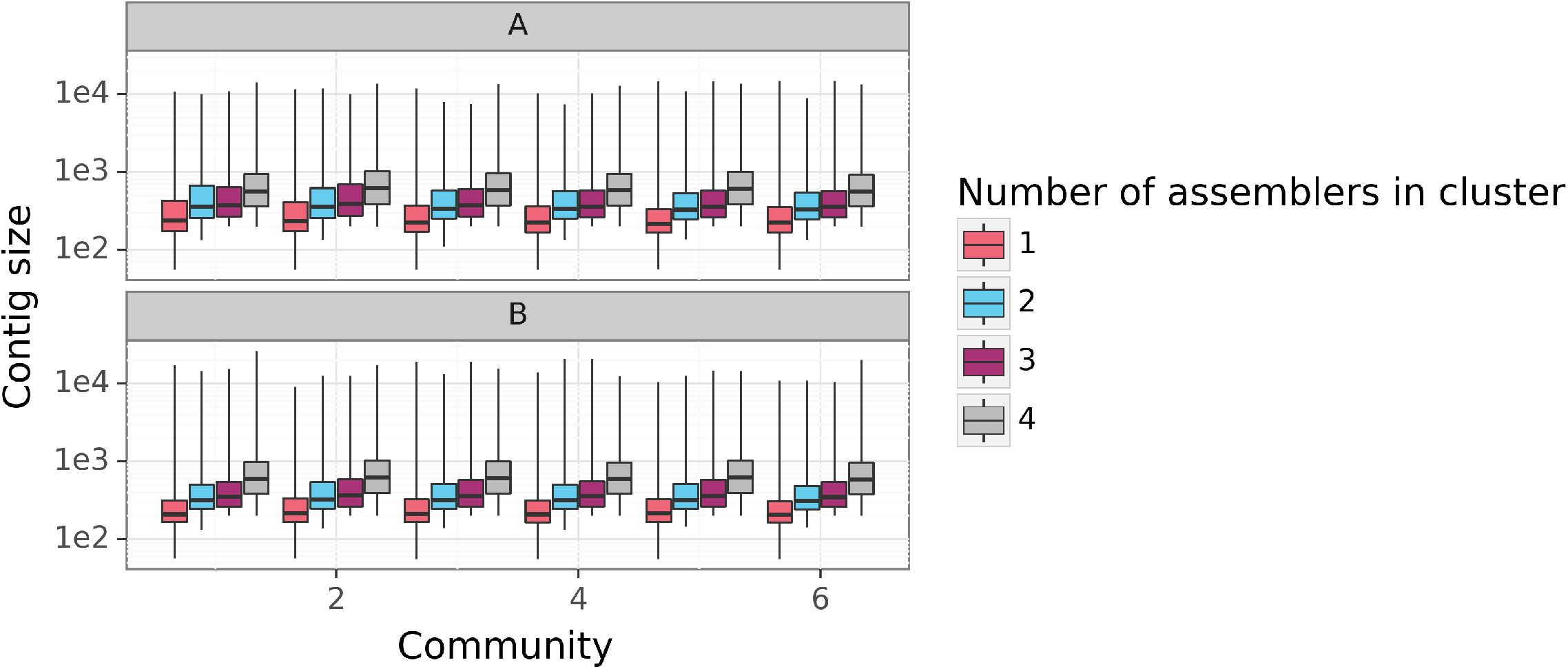
Mean contig length as a function of the number of assemblers that found a sequence that matched the given description. For the 4-assembler cluster, this means that all four assemblers tested identified a sequence that matched the sequence included in the distribution when clustered within eukrhythmic. Panel A corresponds to MMETSP Group A while Panel B corresponds to MMETSP Group B. Welch’s independent T-tests and Kolmogorov-Smirnoff tests for between-distribution goodness of fit computed on these length distributions reveals that the overall distribution of lengths for 1 vs. 2 vs. 3 vs. 4-assembler distributions are all statistically significantly different (*p <* 1*e* − 6), with greater assemblers within a cluster leading to higher average length.

**Figure 4–Figure supplement 4.**
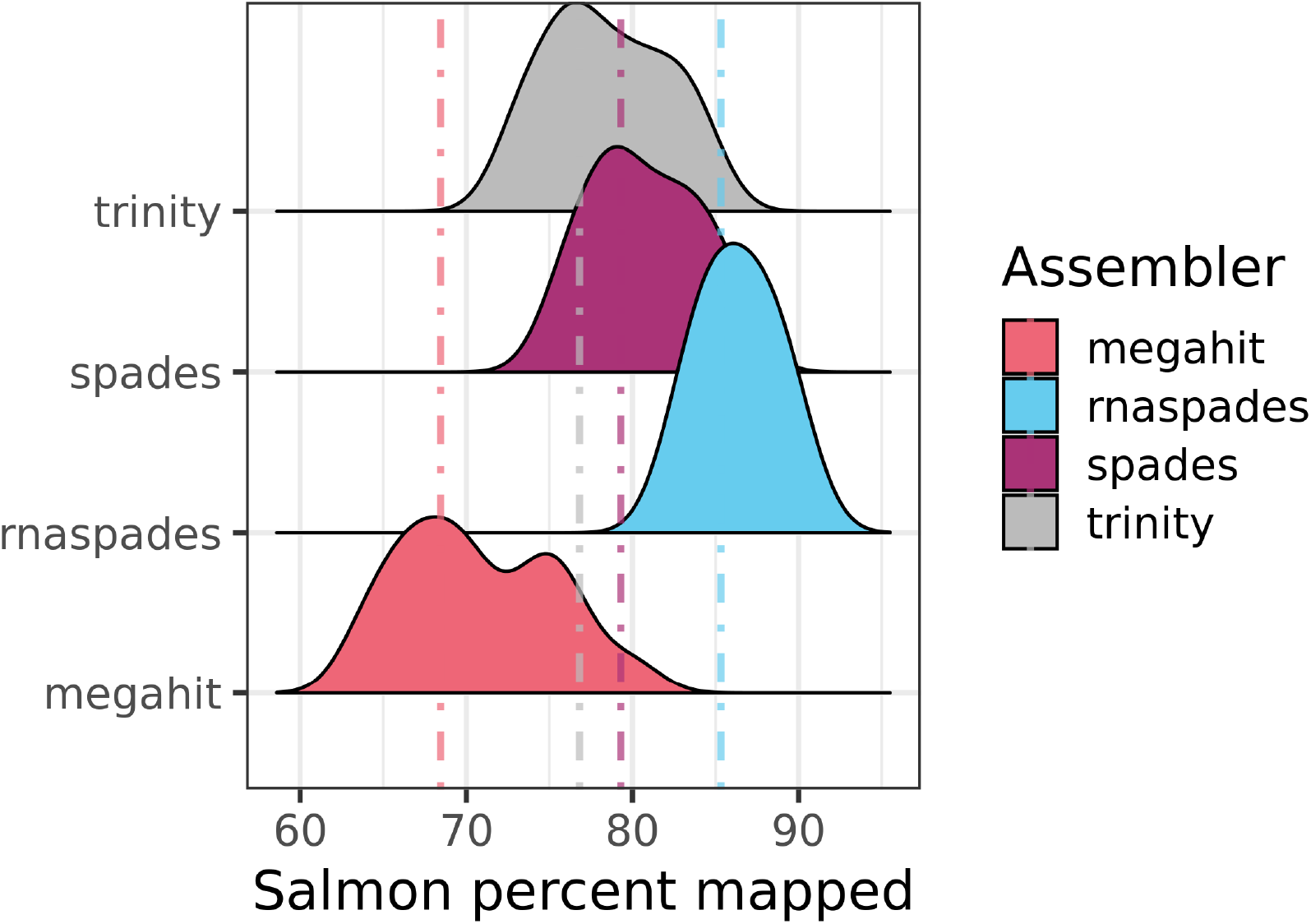
Salmon mapping percentages against the simulated raw reads when computed individually against each of the four assembly tools used by eukrhythmic. rnaSPAdes consistently outperformed the other assemblers with regard to percentage mapping, average length, and number of annotations.

**Figure 5–Figure supplement 1.**
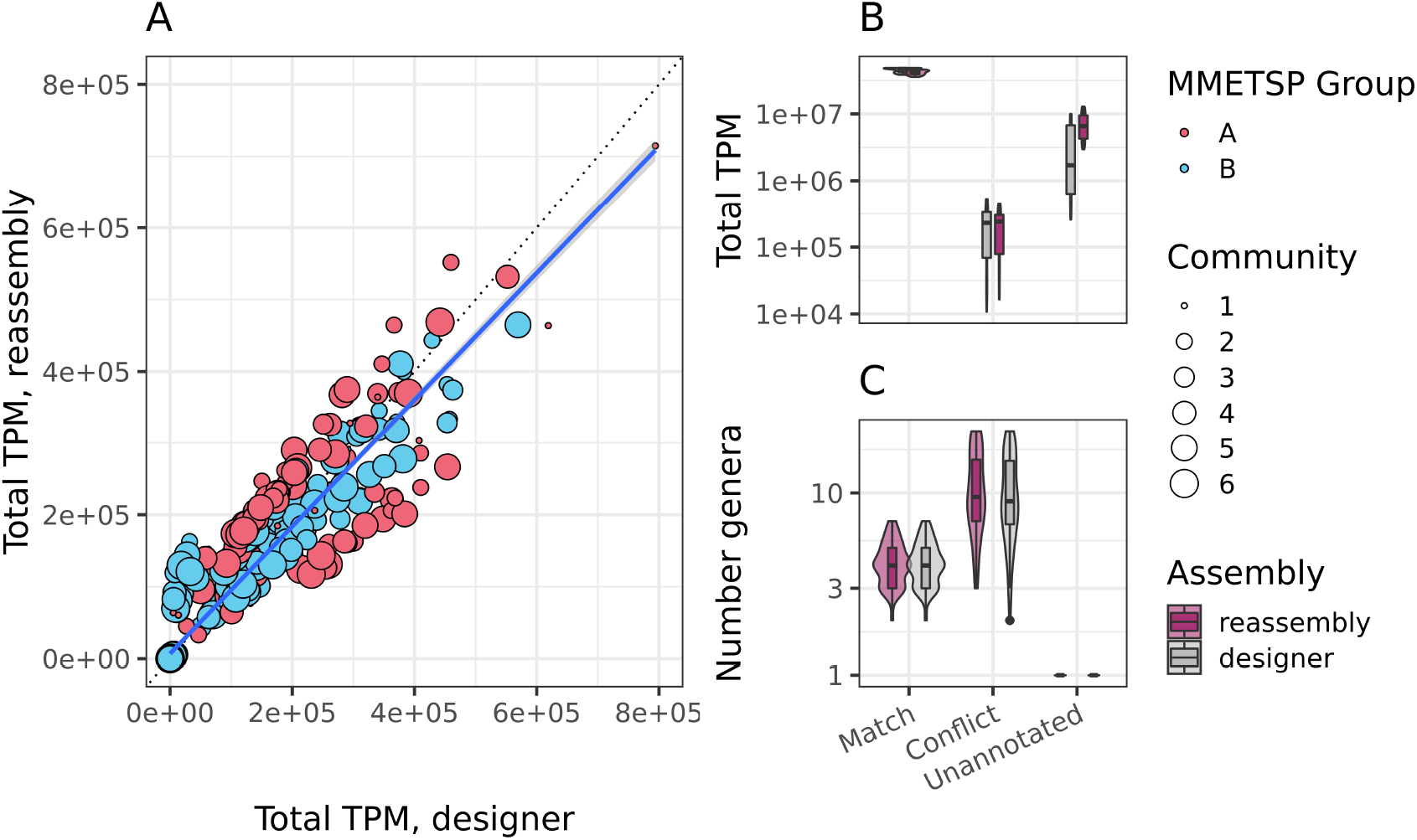
Main text figure, but with A: TPM comparison for the designer assemblies and the reassembled products from *euk* `rhythmic` labelled by distinct groupings of simulations relative to their subset of MMETSP organisms; each point is filled according to its “MMETSP group”.

**Figure 6–Figure supplement 1.**
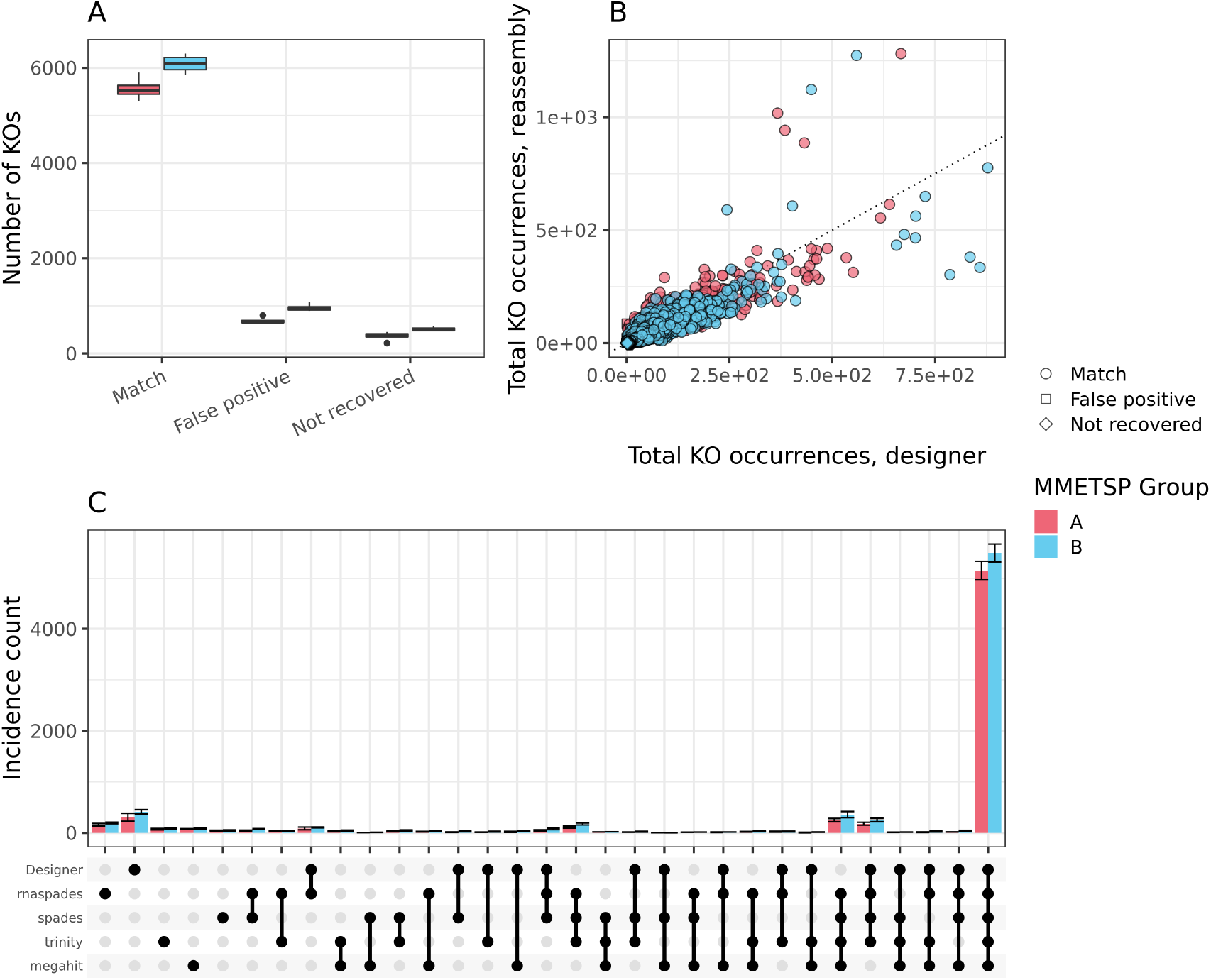
Main text figure with the results split by “MMETSP group” to demonstrate that different taxonomic groupings of organisms included in the simulation does not impact the general trends observed in the results. All three panels are divided by samples using the two “MMETSP groups” of individual organism transcriptomes.

**Figure 7–Figure supplement 1.**
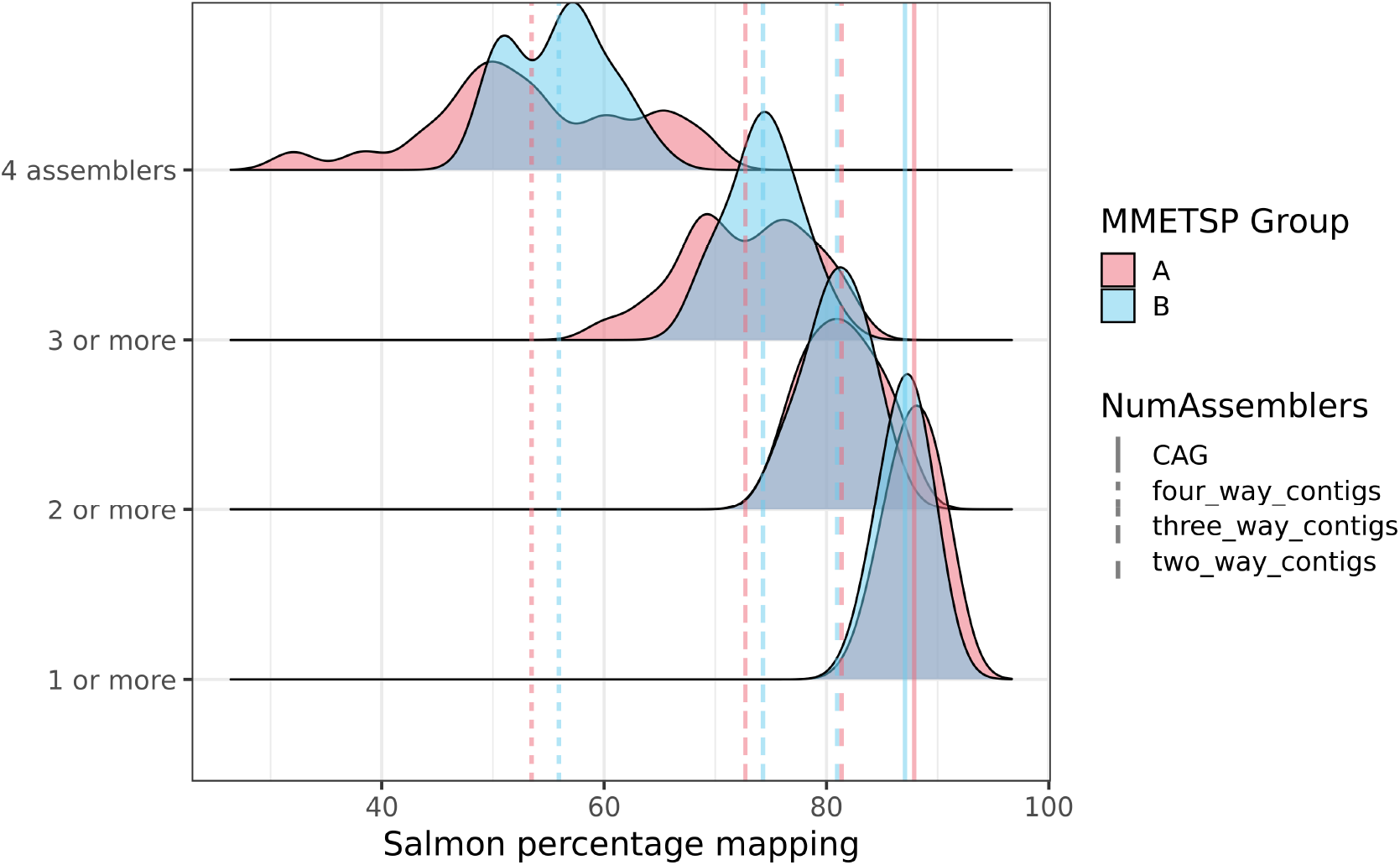
Salmon percentage mapping by MMETSP group when contig products from all assemblers as in eukrhythmic are used (bottom distribution with one or more assembler) as compared to when only contigs agreed upon by multiple assemblers are used. Average percentage mapping decreases progressively with fewer contigs being included as the criteria for inclusion are made stricter.

**Figure 7–Figure supplement 2.**
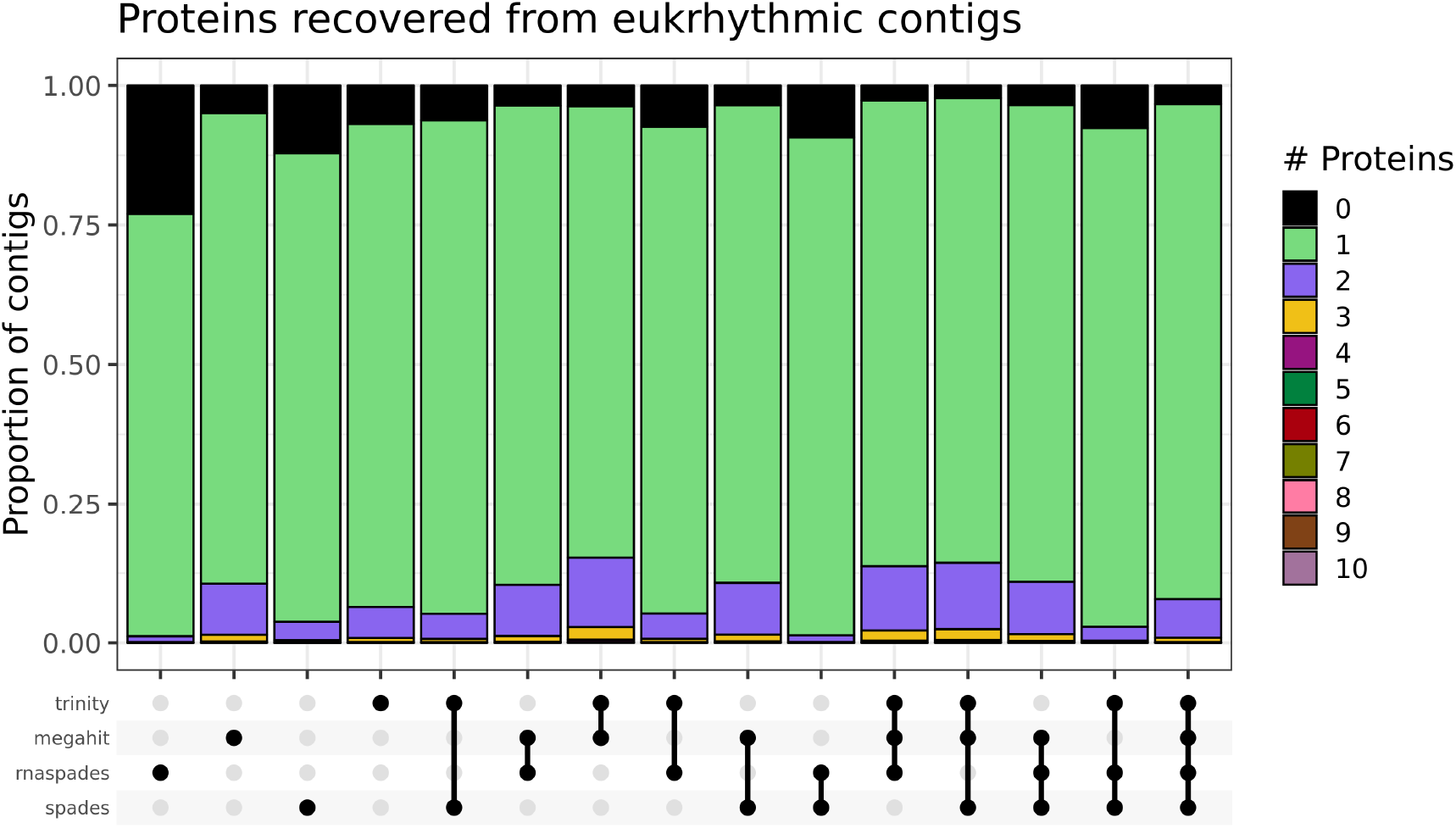
Proportion of contigs from each clustering subset that had ORFs extracted from the sequence. The vast majority of contigs had a single predicted ORF, but `rnaSPAdes` alone had the greatest number of nucleotide contigs on which an ORF could not be detected. In practice, these sequences might be assumed to be non-coding.

**Figure 7–Figure supplement 3.**
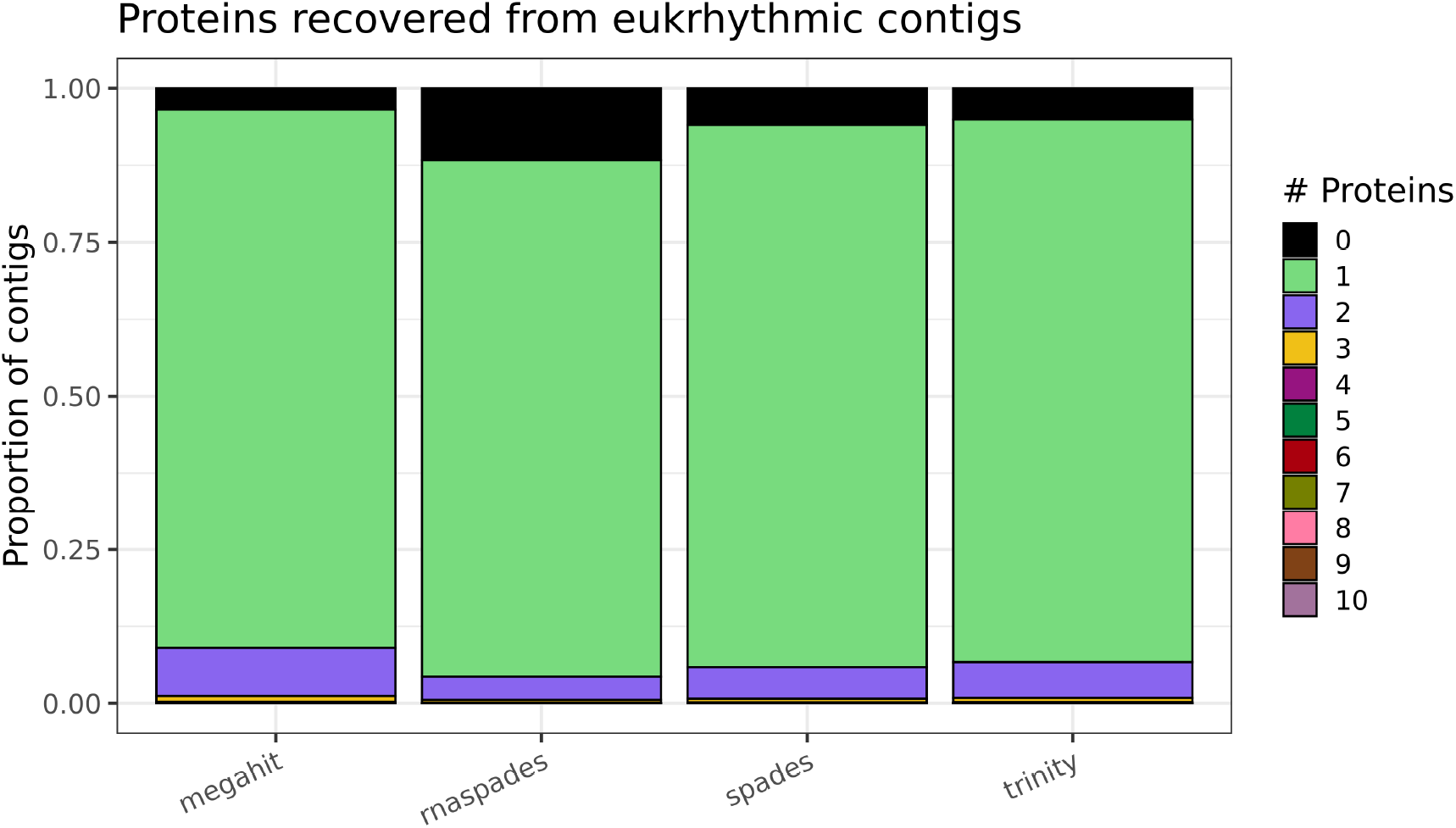
Proportion of contigs from each clustering subset that had ORFs extracted from the sequence across all contigs identified by the assembler. `rnaSPAdes` had a higher number of contigs without an identified ORF.

**Figure 7–Figure supplement 4.**
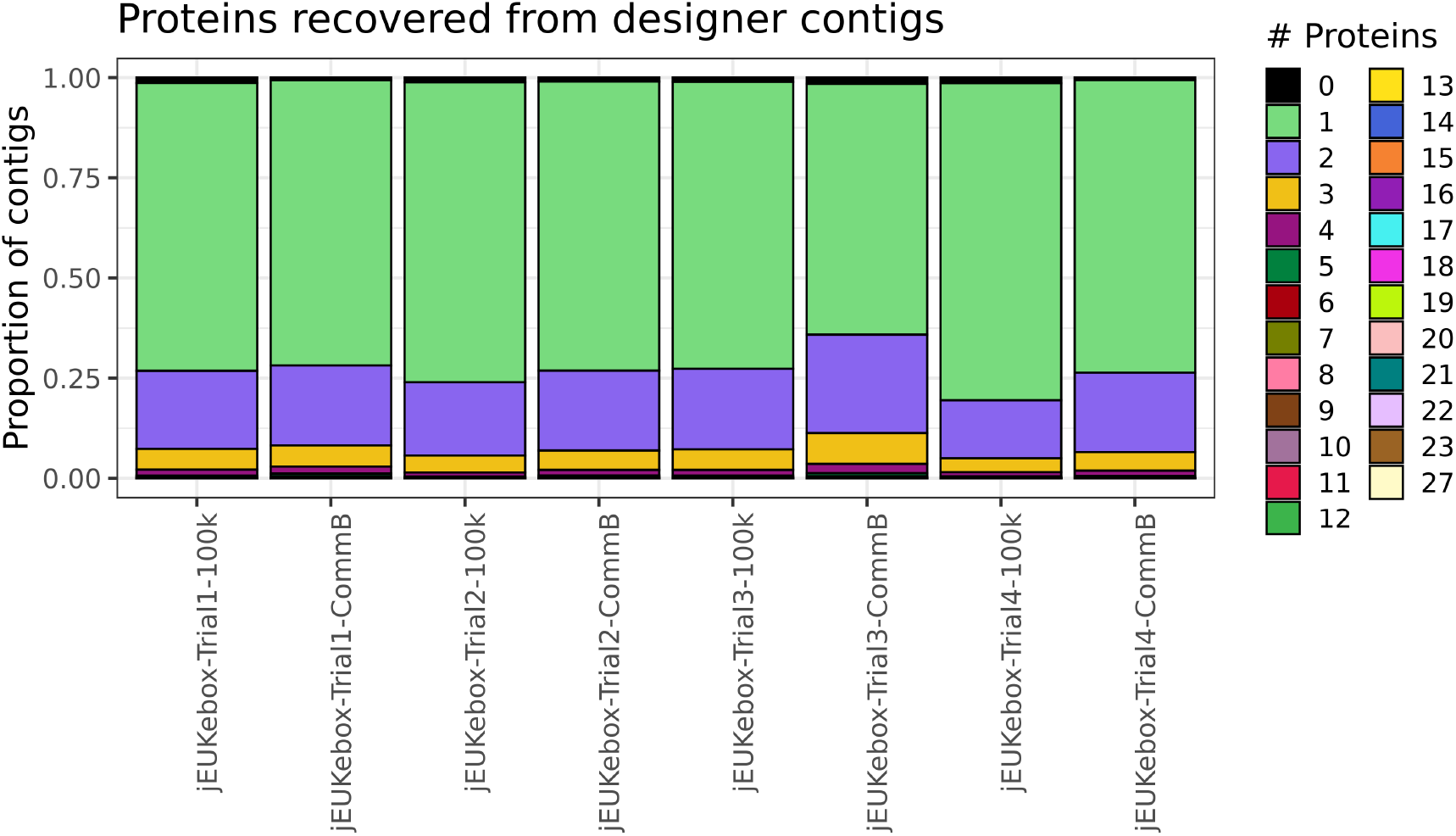
Proportion of contigs from the designer assembly that had ORFs extracted from the sequence. In general, fewer contigs did not have an identified ORF, and a higher number of contigs had multiple predicted ORFs than in the reassembled products from *euk* `rhythmic`.

**Figure 8–Figure supplement 1.**
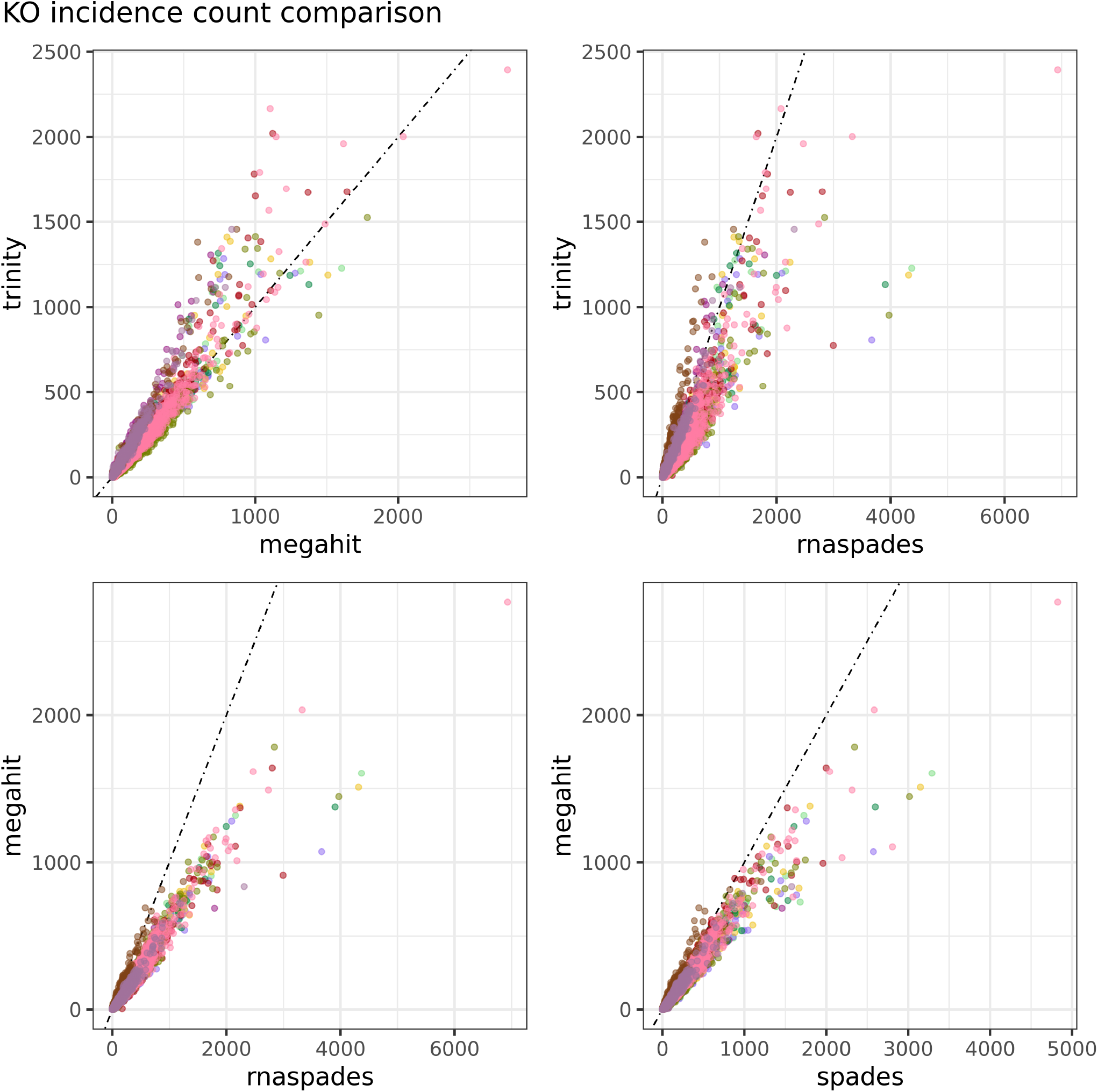
Comparison of the abundance of KO IDs within functional annotations across Narragansett Bay samples and different combinations of metatranscriptome assemblers. A black dotted line indicates a one-to-one relationship, meaning that the abundance of KOs that fall along this line are exactly as abundant using the assembler listed on the x-axis and using the assembler listed on the y-axis. On the top left, `Trinity` is compared to `MEGAHIT`, on the top right `Trinity` is compared to `rnaSPAdes`, on the bottom left `MEGAHIT` is compared to `rnaSPAdes` and on the bottom right `MEGAHIT` is compared to `SPAdes`. Each point corresponds to a single KO within a sample. While `rnaSPAdes` tended to report high abundances of each identified KO relative to the other assemblers, `MEGAHIT` reported fewer instances of each KO than the other three assemblers in most samples. This may be due to the approaches adopted by the two assemblers. Whereas in a typical assembly, *k*-mers that appear only once are assumed to be the result of error, these *k*-mers may represent real and important diversity in a set of low-abundance whole-community sequences ***Li et al. (2015)***. The `MEGAHIT` assembler is an example of metagenomics-specific software that defines “mercy k-mers” which come into play in between two *k*-mers within a single read that are sequenced *more* than once. `rnaSPAdes`, as an example, does not employ a “mercy” strategy but instead decreases the coverage threshold significantly in comparison to genomic assembly

**Figure 8–Figure supplement 2.**
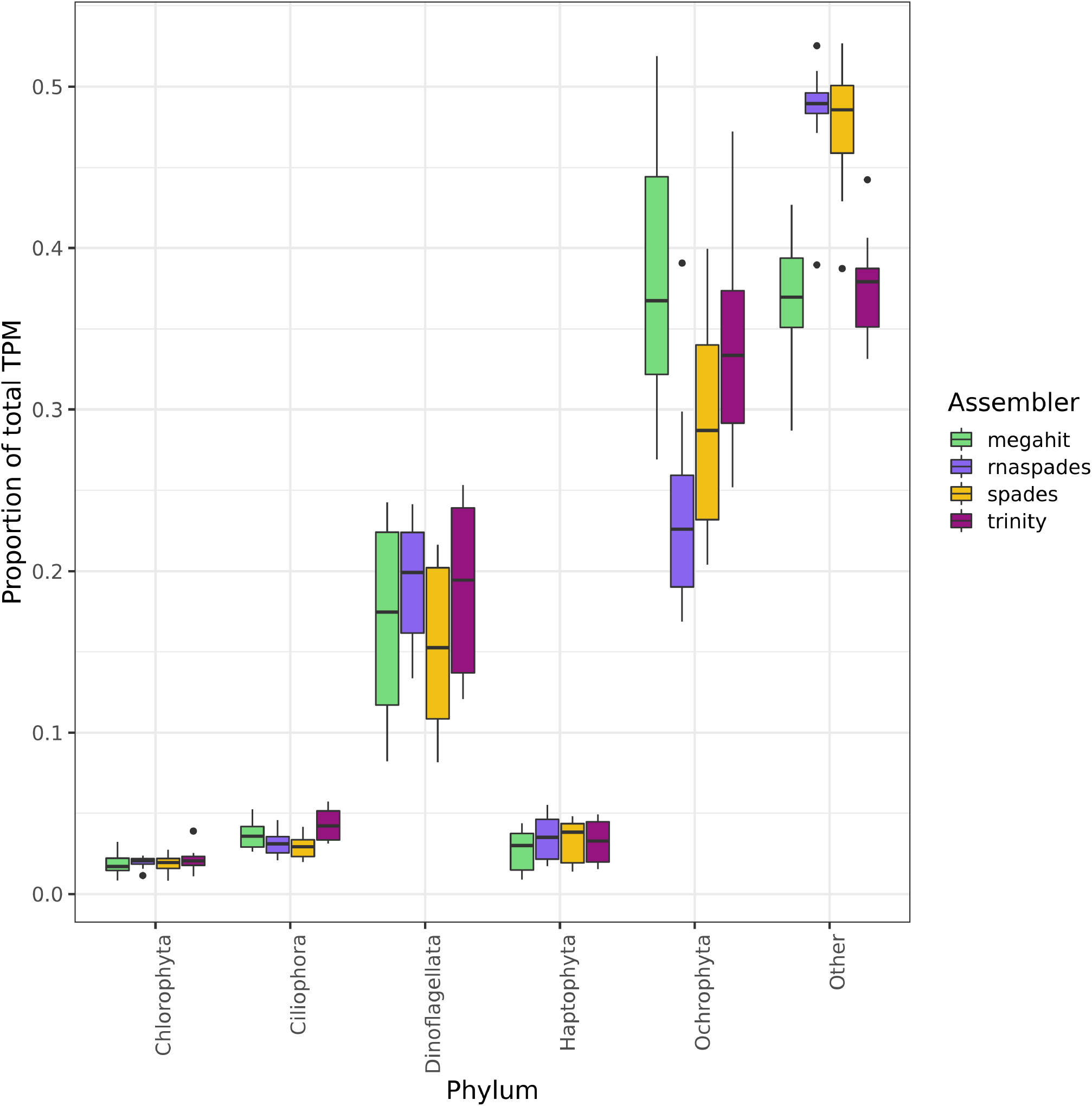
Proportion of total normalized TPM value of major taxonomic groups according to the four metatranscriptomic assemblers that were tested. `rnaSPAdes` had a lower proportion of TPM assigned to Ochrophyta (including diatoms), but further investigation appeared this to be in large part a consequence of the large number of small contigs produced by `rnaSPAdes`.

**Figure 8–Figure supplement 3.**
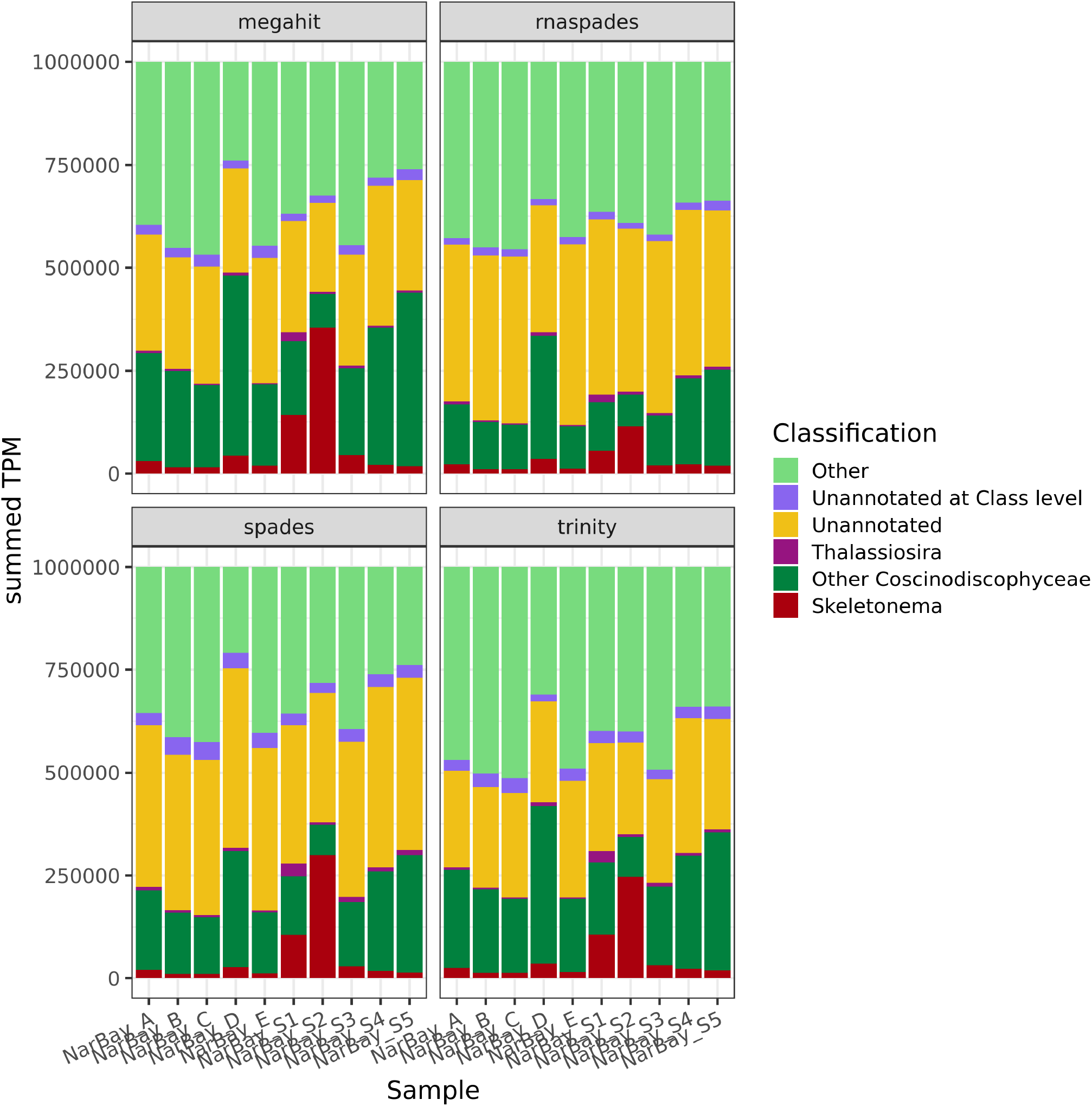
Proportion of total normalized TPM as reported by `Salmon` from the raw reads assigned to each taxonomic category in the assemblies produced by each of the four assemblers (facets). In particular, in sample S2 the taxonomic breakdown of the eukaryotic community differs importantly in the midst of a bloom of the diatom *Skeletonema*.

**Figure 8–Figure supplement 4.**
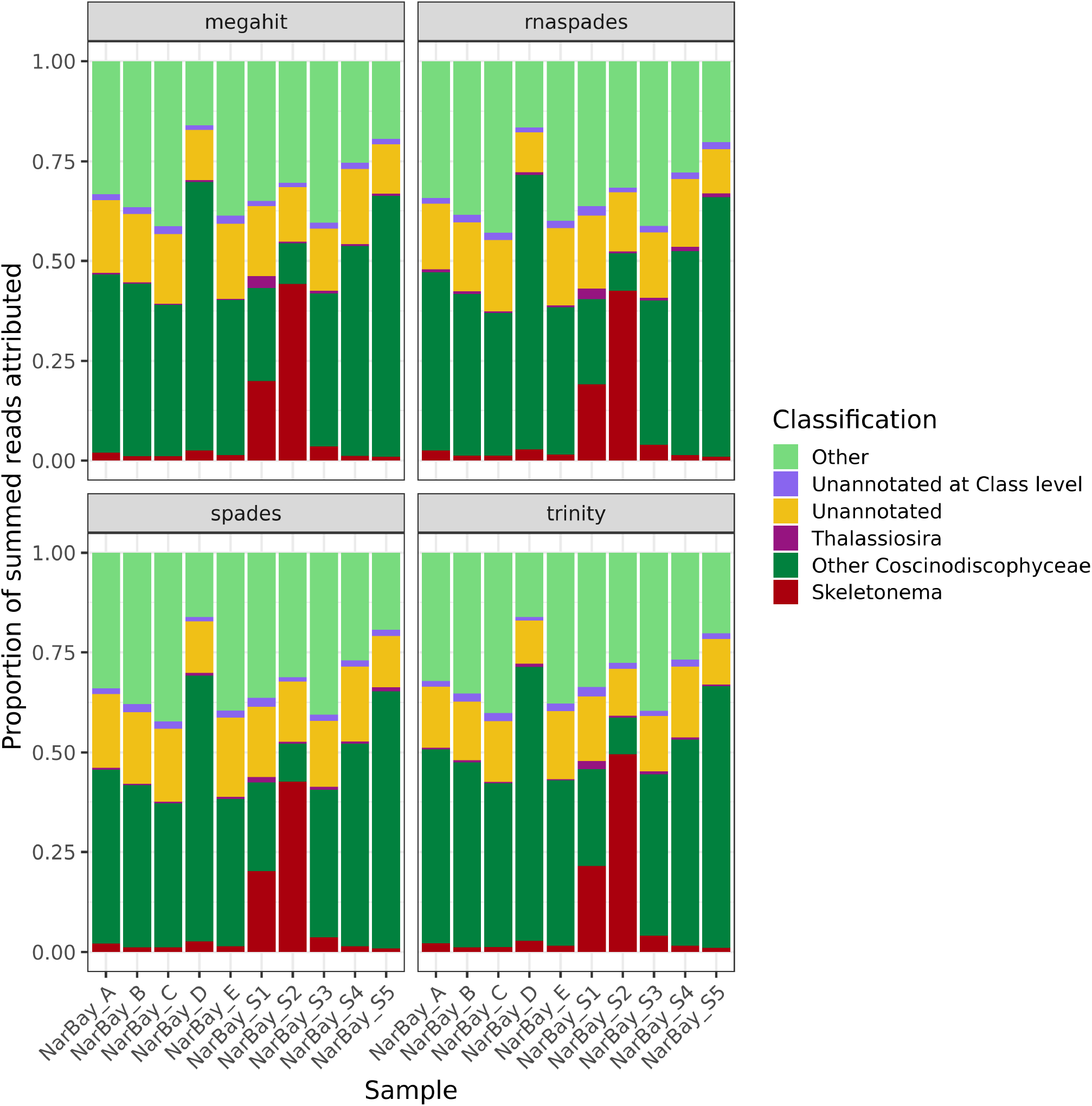
Proportion of assigned raw reads as reported by `Salmon` assigned to each taxonomic category for the four metatranscriptome assemblers, expressed as a proportion of the total.

**Figure 8–Figure supplement 5.**
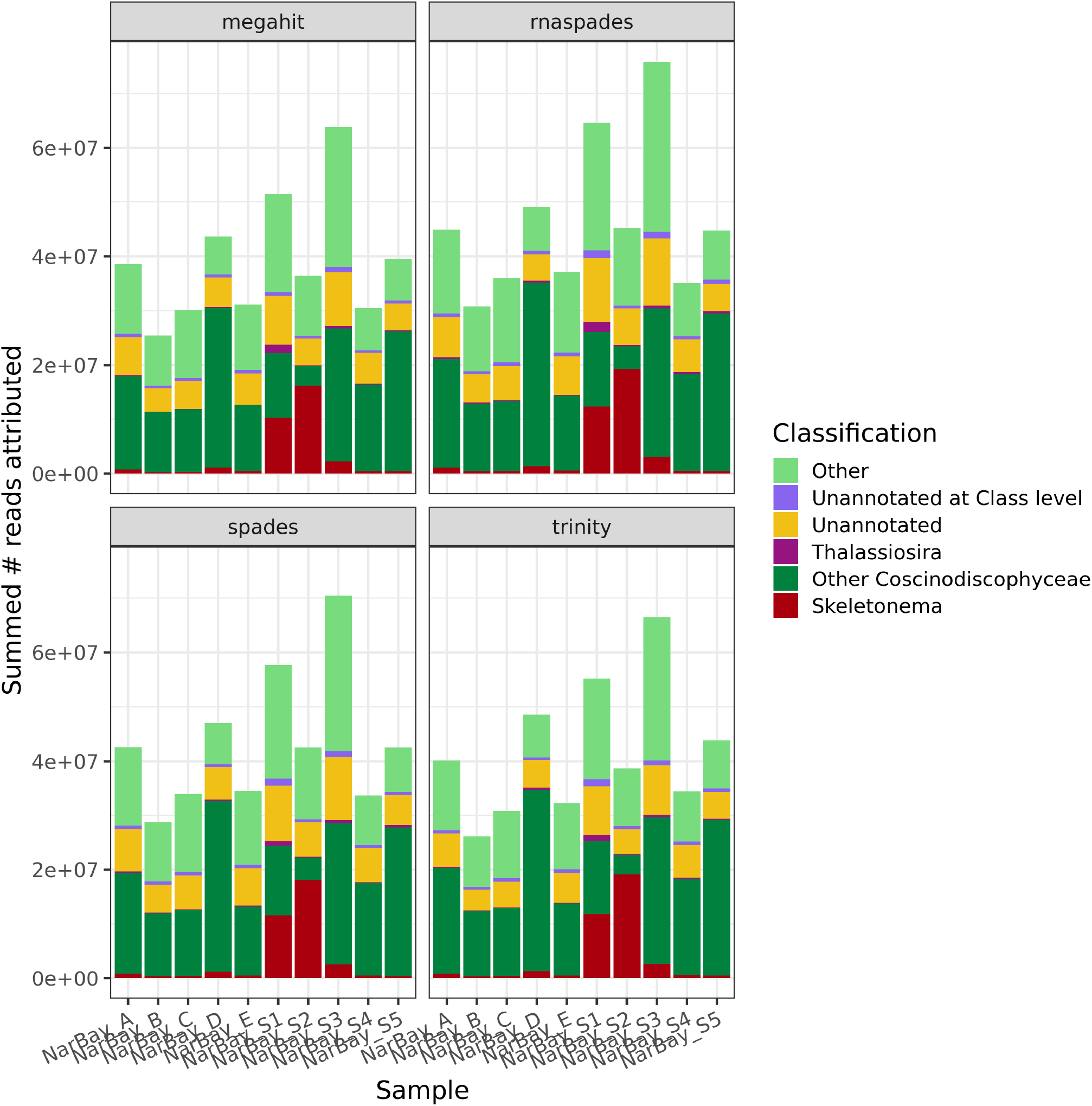
Proportion of assigned raw reads as reported by `Salmon` assigned to each taxonomic category for the four metatranscriptome assemblers, expressed as raw number rather than proportion.

**Figure 8–Figure supplement 6.**
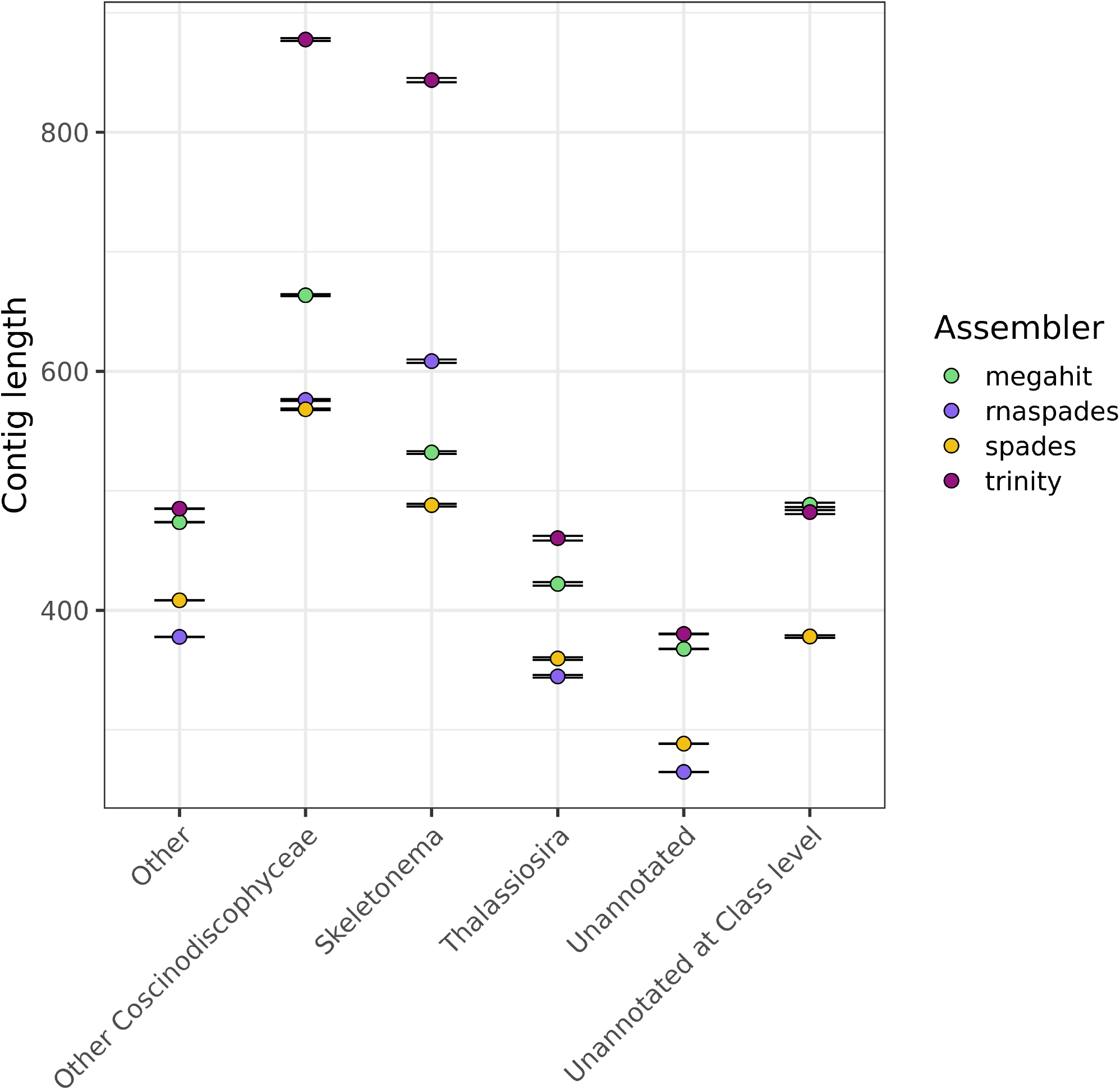
Average contig length by annotation of contigs generated by the four assembly tools across samples. Error bars show the standard error of the mean.

**Figure 8–Figure supplement 7.**
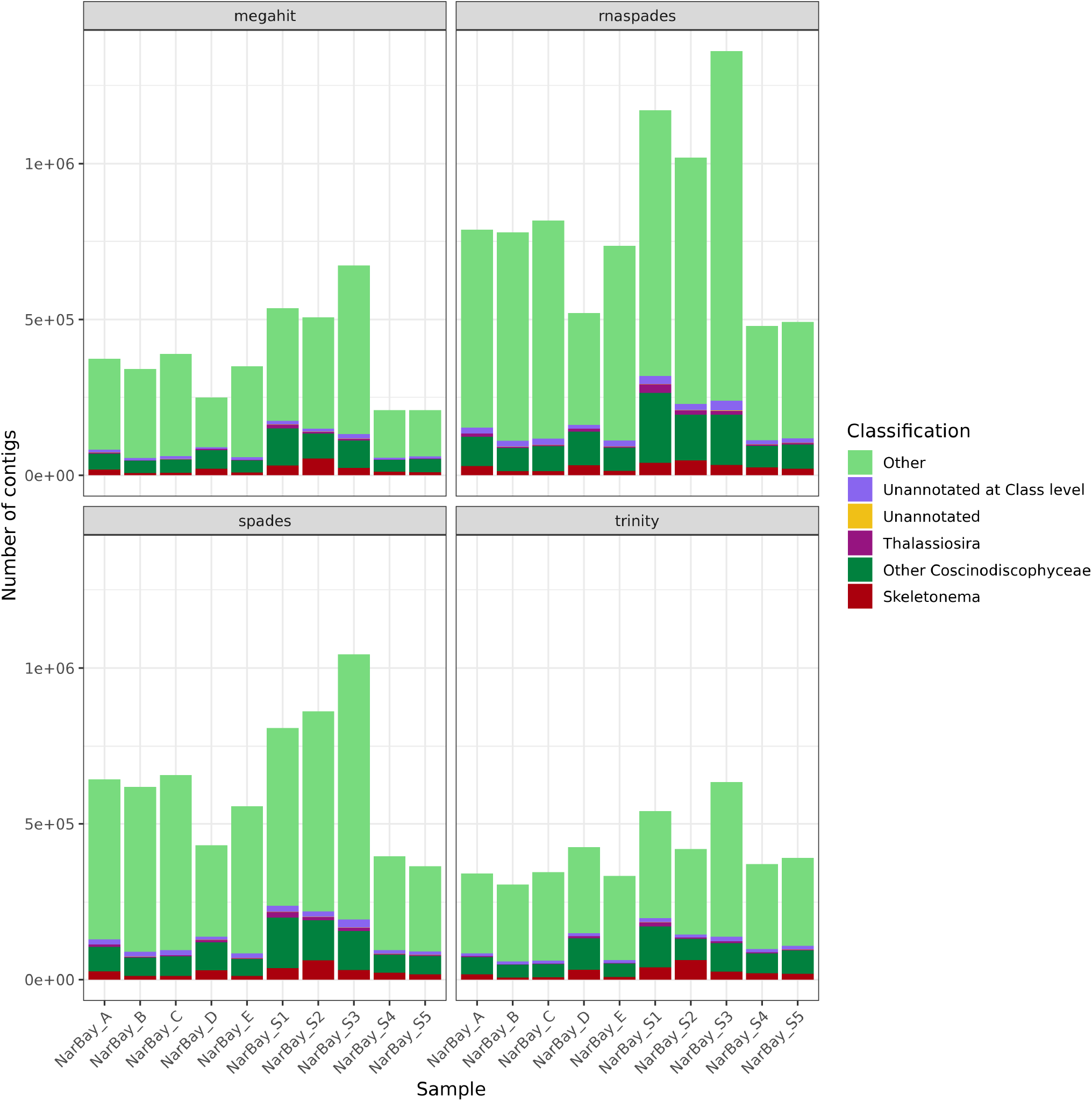
Total number of contigs generated by the four tested assemblers for each of the taxonomic groupings considered. `rnaSPAdes` tended to produce more contigs than the other four assemblers, but these contigs were often shorter and occasionally led to misleading community composition results.

**Figure 8–Figure supplement 8.**
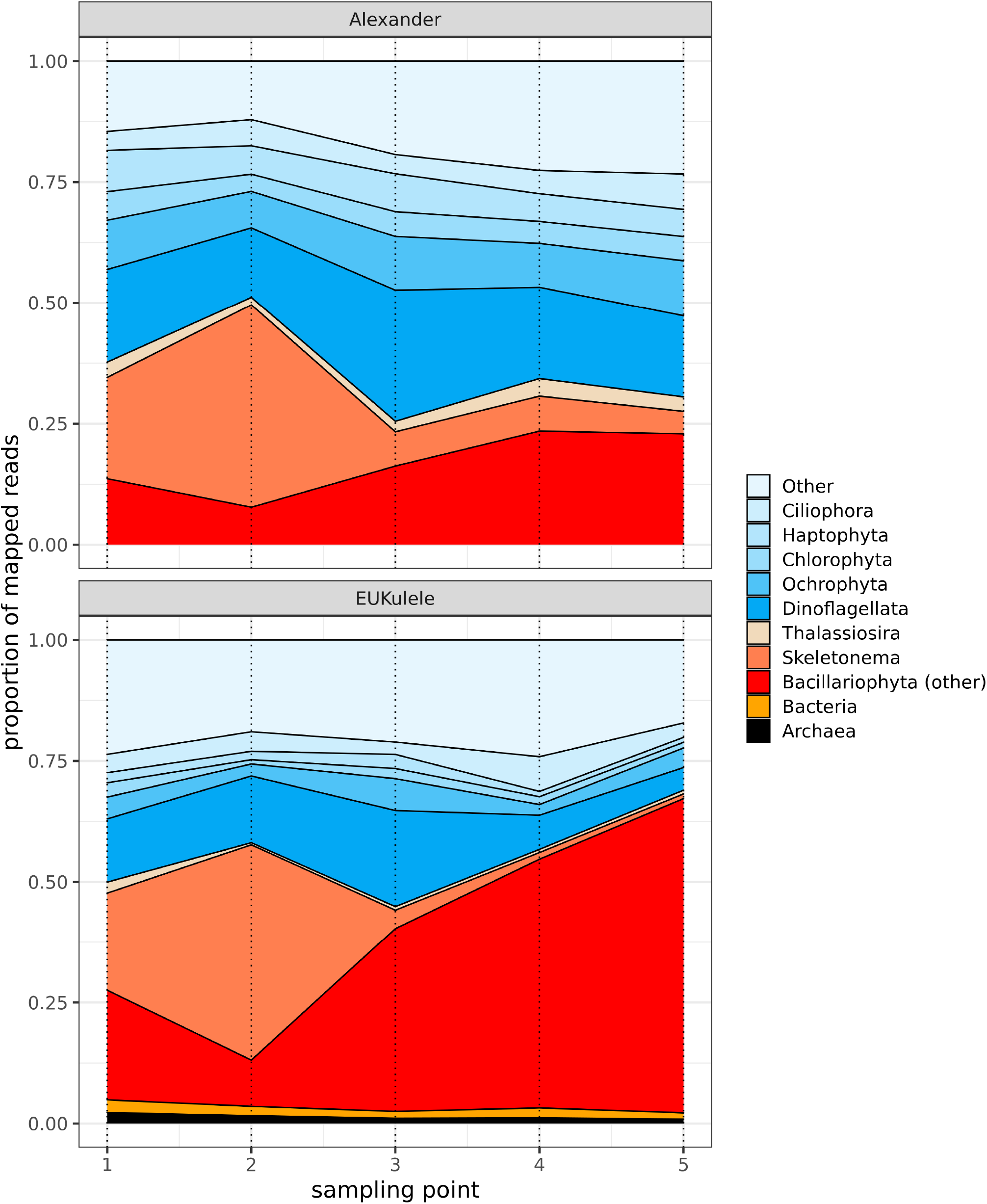
Analogous stacked abundance plots by taxonomic grouping to the relative abundance plots presented in ***Alexander et al. (2015)*** for the five in-situ sampling points collected from Narragansett Bay.

**Figure 9–Figure supplement 1.**
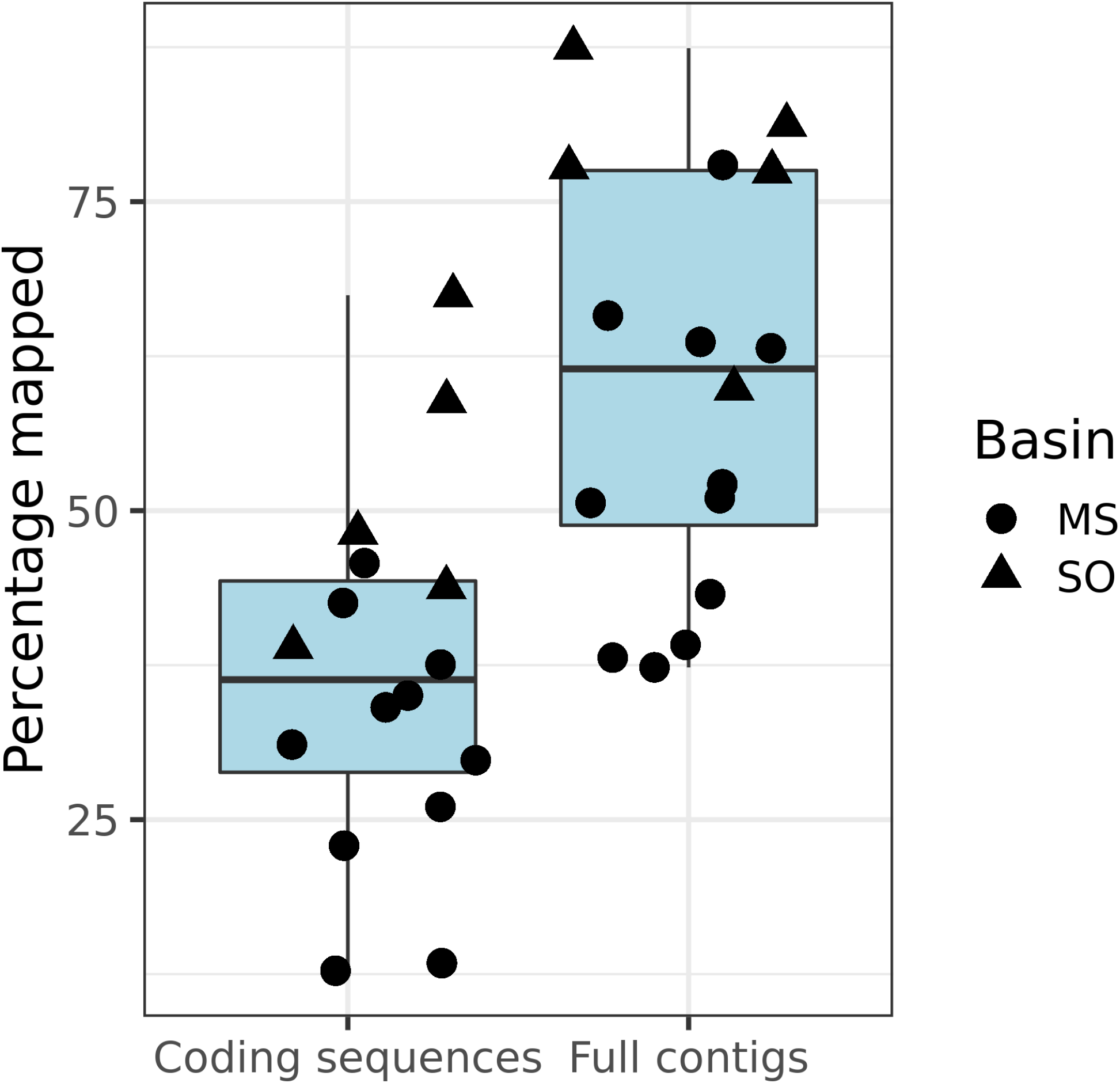
Salmon percentage mapping of coding sequences (left) vs. entire contigs (right) for the *Tara* Oceans samples. Mapping only to coding sequences from the assembly decreased the mean percentage mapped, as reported by Salmon.

**Figure 9–Figure supplement 2.**
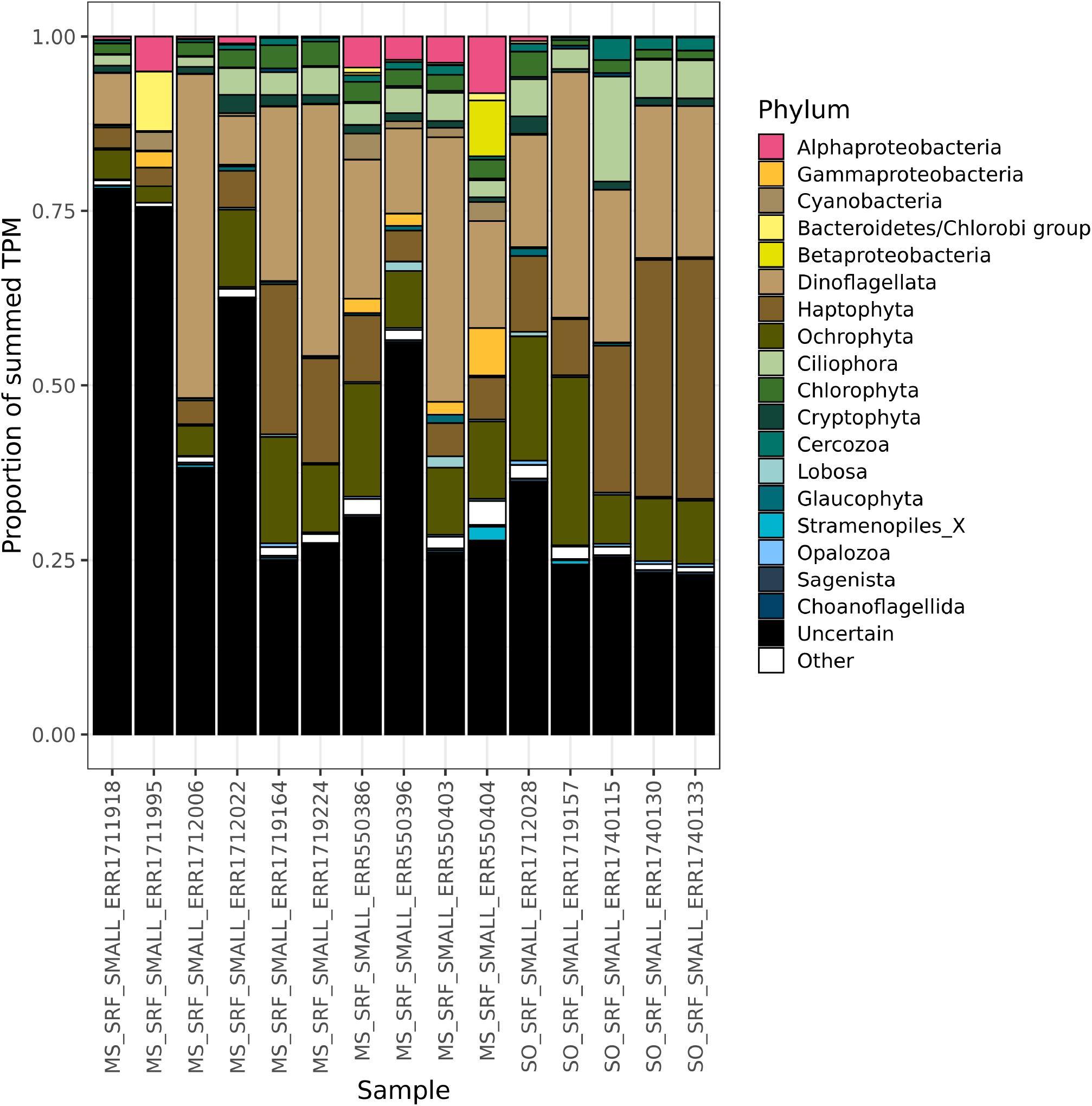
Per-sample taxonomic annotations for all *euk* `rhythmic` CDSs, including those which were and were not found to have a match to the MATOU database.

**Figure 9–Figure supplement 3.**
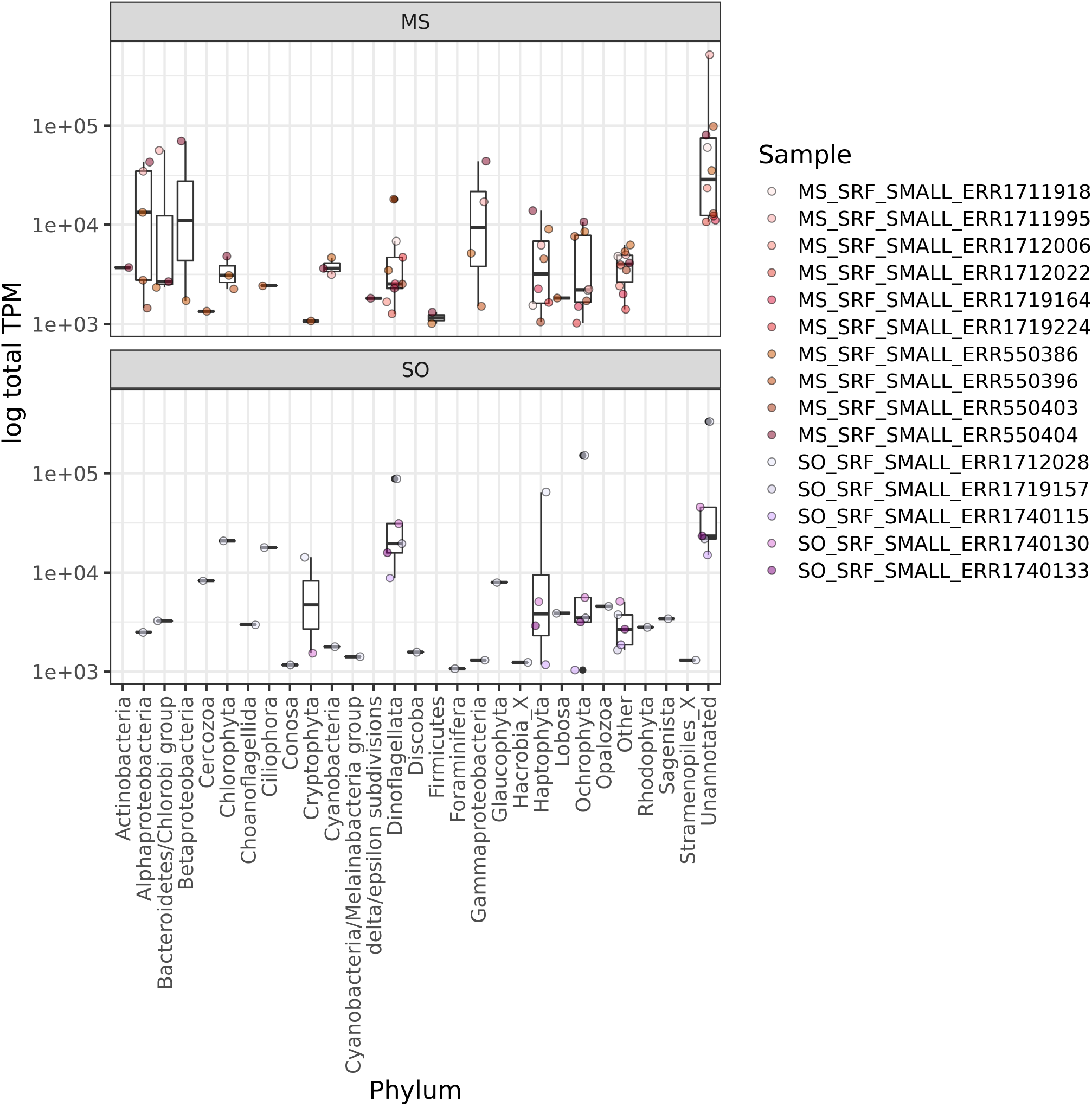
Total TPM assigned per sample to sequences that were assigned a `EUKulele` annotation, but did not have a significant `blast` match to the MATOU database from ***Carradec et al. (2018)***.

**Figure 9–Figure supplement 4.**
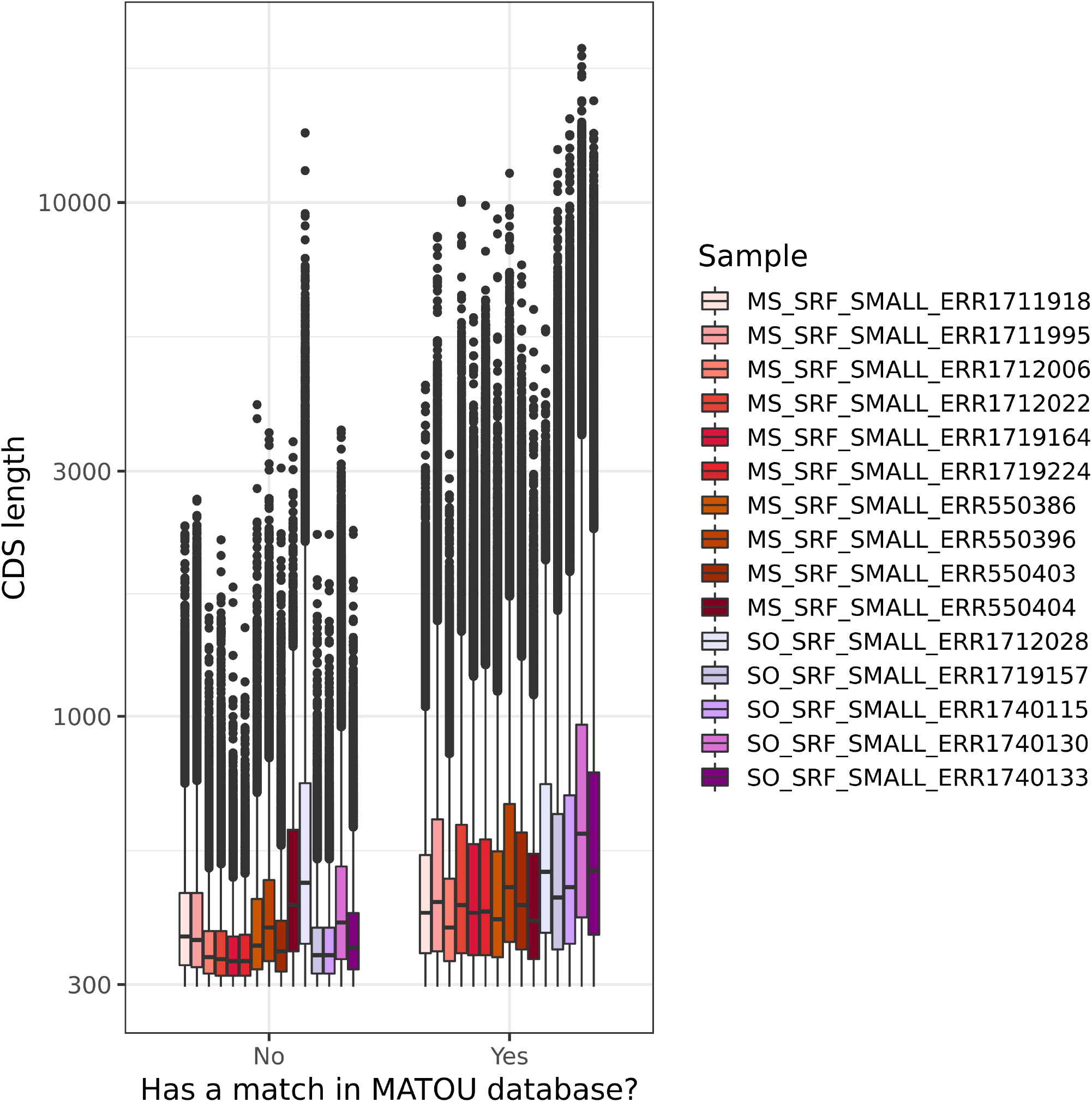
Full length distribution of coding sequences recovered by the eukrhythmic assembly and were (left) not found in the MATOU database ***Carradec et al. (2018)*** vs. (right) found in the MATOU database.

**Figure 9–Figure supplement 5.**
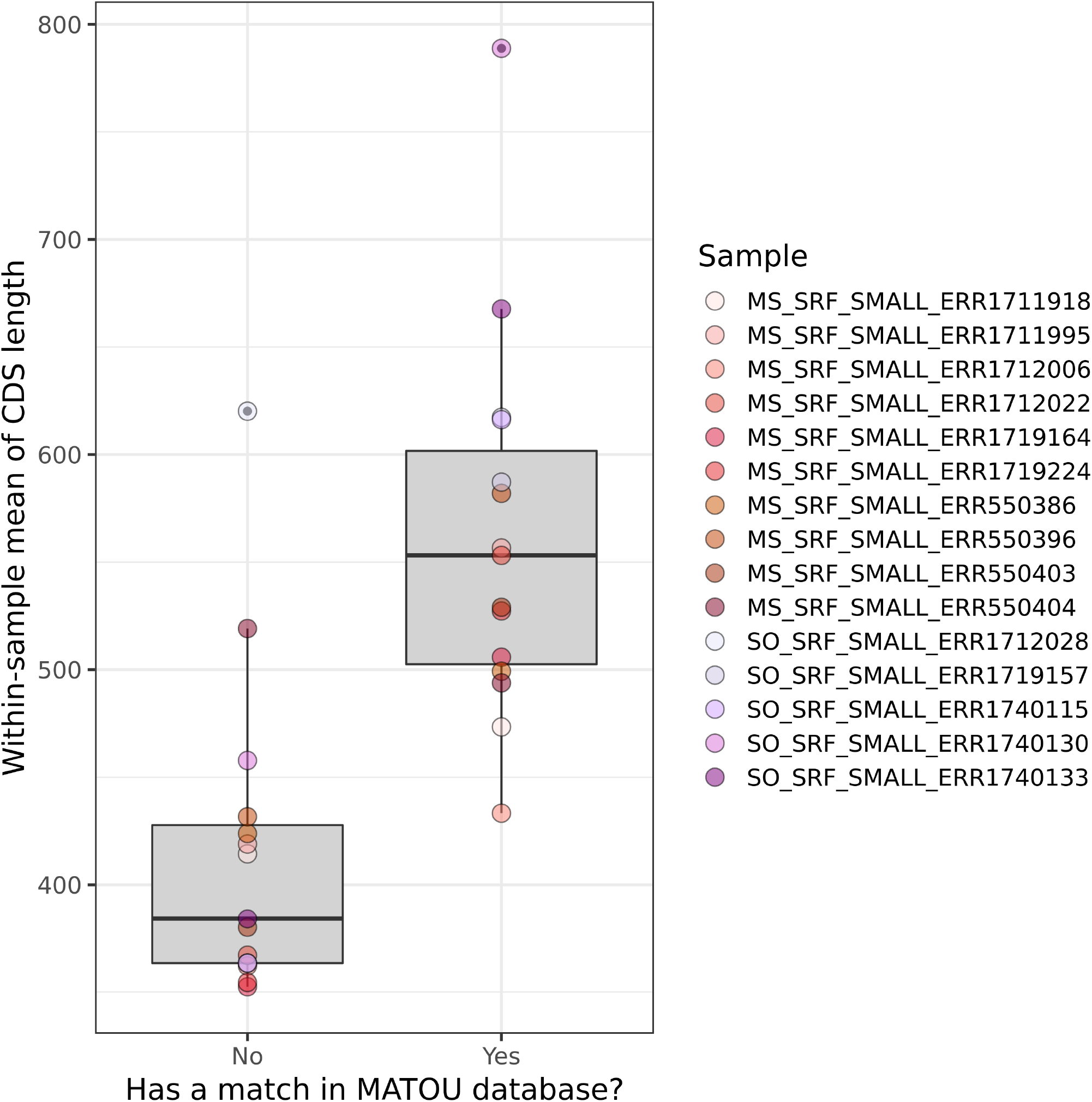
Per-sample average length of coding sequences recovered by the eukrhythmic assembly and were (left) not found in the MATOU database ***Carradec et al. (2018)*** vs. (right) found in the MATOU database.

